# Next generation restoration metrics: Using soil eDNA bacterial community data to measure trajectories towards rehabilitation targets

**DOI:** 10.1101/2021.08.12.456018

**Authors:** Craig Liddicoat, Siegfried L. Krauss, Andrew Bissett, Ryan J. Borrett, Luisa C. Ducki, Shawn D. Peddle, Paul Bullock, Mark P. Dobrowolski, Andrew Grigg, Mark Tibbett, Martin F. Breed

## Abstract

In post-mining rehabilitation, successful mine closure planning requires specific, measurable, achievable, relevant and time-bound (SMART) completion criteria, such as returning ecological communities to match a target level of similarity to reference sites. Soil microbiota are fundamentally linked to the restoration of degraded ecosystems, helping to underpin ecological functions and plant communities. High-throughput sequencing of soil eDNA to characterise these communities offers promise to help monitor and predict ecological progress towards reference states. Here we demonstrate a novel methodology for monitoring and evaluating ecological restoration using three long-term (> 25 year) case study post-mining rehabilitation soil eDNA-based bacterial community datasets. Specifically, we developed rehabilitation trajectory assessments based on similarity to reference data from restoration chronosequence datasets. Recognising that many alternative options for microbiota data processing have potential to influence these assessments, we comprehensively examined the influence of standard versus compositional data analyses, different ecological distance measures, sequence grouping approaches, eliminating rare taxa, and the potential for excessive spatial autocorrelation to impact on results. Our approach reduces the complexity of information that often overwhelms ecologically-relevant patterns in microbiota studies, and enables prediction of recovery time, with explicit inclusion of uncertainty in assessments. We offer a step change in the development of quantitative microbiota-based SMART metrics for measuring rehabilitation success. Our approach may also have wider applications where restorative processes facilitate the shift of microbiota towards reference states.

## 1. INTRODUCTION

Land degradation and transformation, with negative impacts to biodiversity and ecosystem function, are estimated to impact 75% of the Earth’s land surface, and this figure is projected to rise to over 90% by 2050 (IPBES, 2018). Ecological restoration—activity that supports rehabilitation of locally representative, sustainable, biodiverse ecosystems (Gann et al., 2019)—is seen as integral to reversing these impacts, as highlighted by the UN declaration of 2021–2030 as the Decade on Ecosystem Restoration (https://www.decadeonrestoration.org/). Restoration is technically challenging and requires considerable investment, without guaranteed success (Tibbett, 2015). With large investments in restoration (e.g. BenDor et al., 2015 estimate US$9.5 billion/yr is spent in the USA alone; Menz et al., 2013 estimate US$18 billion/yr is required to restore degraded lands globally), there is a need to improve the evidence base to guide continuous improvement in restoration outcomes and to underpin future investment.

Reference ecosystems provide an important basis for establishing targets and monitoring progress of restoration activities (Gann et al., 2019) (online Supporting Information (SI) Appendix, Figure S1). In post-mining contexts, best practice guidelines require formal mine completion criteria to be prescribed in a matter that is specific, measurable, achievable, relevant and time-bound (SMART) (Australian_Government, 2016; Manero et al., 2021). To-date, completion criteria have largely focussed on vegetation community variables, with typical ecological measures including alpha and beta diversity reflecting the number of different taxa and community composition, respectively. For example, targets may be set at a minimum threshold similarity to a reference community. Despite available guidance, many completion criteria are ambiguous or ill-defined, and can result in unclear standards for regulators, unrealistic expectations for stakeholders, and represent a key barrier to the relinquishment of minesites (Manero et al., 2021). To help move the industry towards improved definitions of completion criteria, Manero et al. (2021) suggest criteria for industry best practice, which include using multiple reference sites, monitoring and corrective actions (i.e., adaptive management), allowing innovation-guided completion criteria, and specific objectives and indicators.

Soil microbial communities (microbiota) have essential roles in organic matter decomposition, soil formation, and nutrient cycling, and therefore help regulate plant productivity and community dynamics (Harris, 2009). Patterns of land use, vegetation communities, and soil quality each help to shape soil microbiota (Bulgarelli et al., 2013; Delgado□Baquerizo et al., 2018; Turner et al., 2013). Microbiota depend on the resource and energy flows associated with aboveground biota, and therefore their monitoring may help indicate the impact of restoration interventions (Harris, 2009; Jiao et al., 2018; van der Heyde et al., 2020).

The development of low-cost, high-throughput sequencing of environmental DNA (eDNA) has enabled affordable, rapid and comprehensive assessment of soil microbiota. Applying recognised ecological assessment approaches to abundant eDNA-based microbiota data has potential to provide a novel tool for measuring trajectories and predicting time to recover towards restoration targets (Rydgren et al., 2019). Chronosequence study designs, while containing limitations (Walker et al., 2010), are commonly used to examine ecosystem recovery following restoration activities (Tibbett, 2010). However, there are few studies of soil microbiota from restoration chronosequences that explicitly visualise and evaluate patterns in ecological similarity to reference data with time since rehabilitation. It is customary for such studies (e.g., Fernandez Nuñez et al., 2021; Jiao et al., 2018; Schmid et al., 2020) to examine patterns in microbiota composition via analysis of taxonomic groups and ordination techniques which project multivariate community data into lower dimensional space (e.g., 2-d plots). These popular techniques often characterise the complexity and site-specificity of soil ecosystems. However, a focus on measuring ‘similarity to reference’ may help cut through the complexity inherent to microbiota data. Along these lines, van der Heyde et al. (2020) visualised temporal trends in ecological similarity to reference data in post-mining rehabilitation—however, in their example each rehabilitation sample was only compared to a single closest reference sample, which potentially limited insight into variability and uncertainty in microbiota recovery.

Here we provide a proof-of-concept demonstration and detailed exploration of a new complexity-reducing application of eDNA-based soil bacterial community data to assess the progress of post-mining rehabilitation using three long-term (> 25 year) chronosequence case studies from south-west Western Australia. Specifically, we aim to demonstrate the use of chronosequence-based rehabilitation trajectories, using measures of percent similarity of bacterial community structure to ecological reference sites (hereafter termed references), to assess progress of soil bacterial communities towards reference states with increasing rehabilitation age. We note that further work that links microbiota to other ecosystem components (e.g., vegetation, fauna) is important but beyond the scope of our study.

Our intended audience includes microbiome researchers working in ecosystem restoration, as well as restoration managers who are considering new methods to add to their ecological monitoring toolkit. Our approach may also have applications for monitoring and predicting microbiota recovery toward reference states in diverse fields, such as microbiota-mediated human health (Lloyd-Price et al., 2016) and microbiota-conscious urban design (Watkins et al., 2020).

Due to the potential for alternative data processing options to cause varying impacts on our rehabilitation trajectory assessments, we compare outcomes from a range of potential options relevant to microbiota data analyses. For example, compositional data analysis approaches are promoted to have greater statistical rigour compared to standard approaches (Gloor et al., 2017); grouping bacterial taxa based on sequence similarity (i.e., varying the resolution of operational taxonomic units, OTUs) might help manage noise associated with microbiome data; taxonomic grouping might assist interpretation if recognised groups can be discussed; and eliminating rare taxa (to simulate reduced sequencing depths) might allow more cost-effective and rapid analyses. We also recognise the potential for spatial autocorrelation—where measured outcomes are closer in value due to closer spatial proximity—to confound the assessment of rehabilitation age in chronosequence studies that lack appropriate spatial design and replication. Accordingly, our *a priori* research questions were: (1) can soil bacterial community data be used to establish reference-based targets? (2) can soil bacterial community rehabilitation trajectory data be used to predict the time to recover to reference targets? and (3) how are these predictions of recovery influenced by different ecological distance/similarity measures and sequence data resolution? (4) Additionally, we conduct a preliminary, illustrative examination of spatial autocorrelation, and trial an approach to highlight and ‘correct’ datasets where its influence appears excessive. We then discuss limitations and synthesise our findings to inform future work.

## 2. MATERIALS AND METHODS

### 2.1 Data collection

We used surface soil bacterial 16S rRNA marker gene data from three case study minesites (Figure 1; SI Appendix, Tables S1–S3) from south-west Western Australia. Soil sampling was undertaken in accordance with Australian Microbiome (AM) protocols (Bissett et al., 2016; https://www.australianmicrobiome.com/protocols; SI Appendix, Supplementary Methods). Each minesite experiences a Mediterranean-type climate with hot, dry summers and cool, wet winters. Post-mining rehabilitation activities typically involved deep-ripping, prior to the ‘direct return’ (where possible) of subsoil and topsoil stripped from a separate pit about to be mined, followed by revegetation with locally appropriate seed of diverse plant communities (Tibbett, 2010). Precise soil handling and storage techniques differed between the minesites and different pits within minesites. Summary information for each minesite is provided below (see SI Appendix, Supplementary Methods for more background information; other studies in-progress will provide expanded analyses of surface and subsoil data from these minesites, including additional marker gene datasets).

**FIGURE 1.**
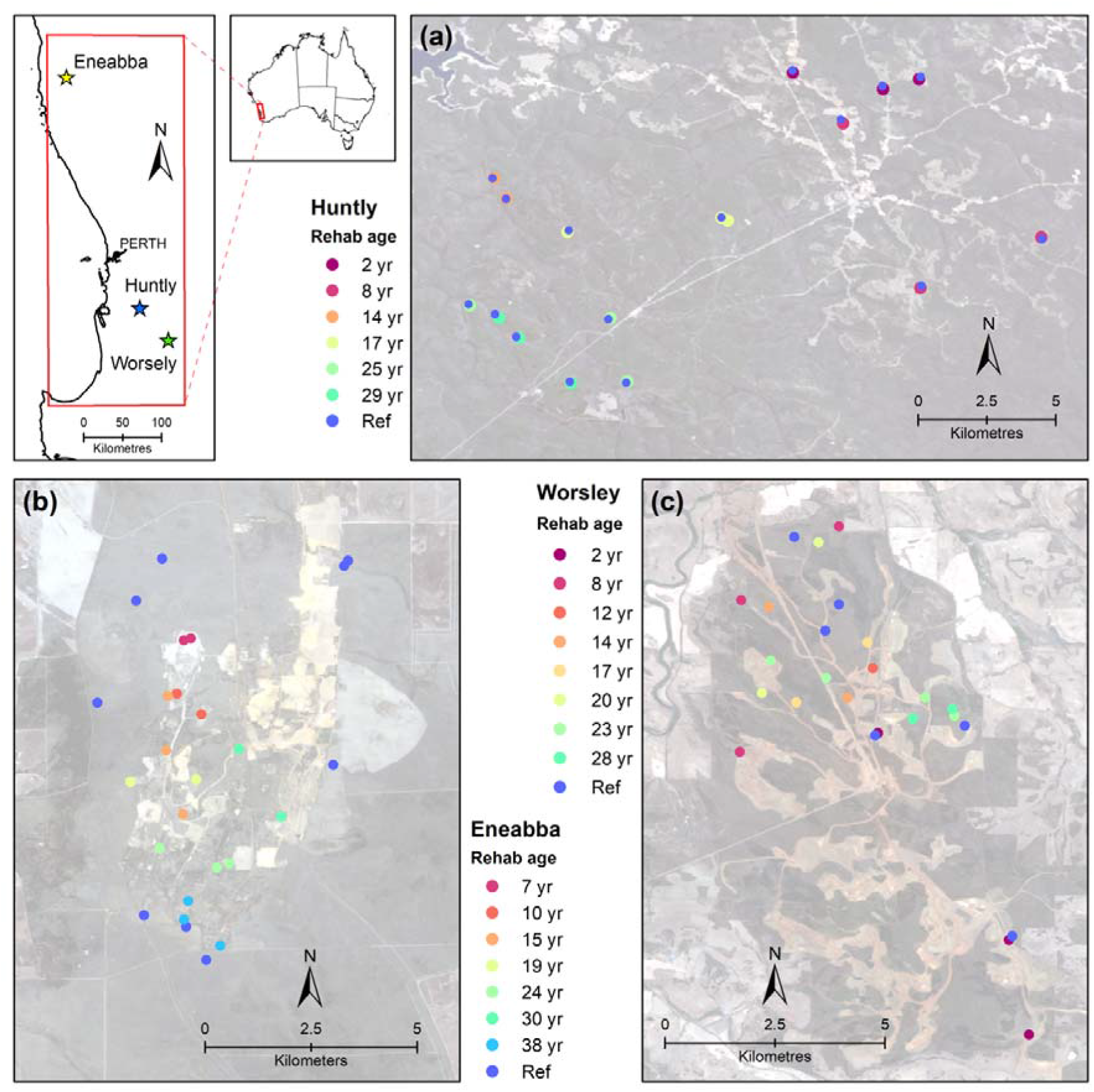
Locations of minesites and soil sampling sites: (a) Huntly, (b) Eneabba, (c) Worsley. (Imagery: Sentinel-2; https://eos.com/landviewer; EOS Data Analytics, Inc.)

Alcoa’s *Huntly* bauxite-producing minesite is approximately 100 km south-east of Perth, occurring in mixed open forest with dominant overstorey species of Jarrah (*Eucalyptus marginata*) and Marri (*Corymbia calophylla*) on lateritic, nutrient poor soils. We consider Huntly data sampled in 2016, with rehabilitation ages between 2–29 years old. Huntly’s 36 samples correspond to rehabilitation years: 1987 (n = 3), 1991 (n = 3), 1999 (n = 3), 2002 (n = 3), 2008 (n = 3), 2014 (n = 3), reference (n = 18), where each reference site was paired with an adjacent rehabilitation site.

Iluka Resource’s *Eneabba* mineral-sand minesite is approximately 280 km north of Perth, occurring in sandplain heath vegetation comprising low shrubland on undulating infertile siliceous sandplains, predominantly featuring perennial woody species from the Proteaceae, Myrtaceae, and Fabaceae families. We consider Eneabba data sampled in 2019, with rehabilitation ages between 7–38 years. Eneabba’s 26 samples correspond to rehabilitation years: 1981 (n = 3), 1989 (n = 2), 1995 (n = 3), 2000 (n = 2), 2004 (n = 3), 2009 (n = 2), 2012 (n = 2), reference (n = 9).

South32’s *Worsley* bauxite-producing minesite is located approximately 150 km south of Perth, occurring in Jarrah (*Eucalyptus marginata*) forest on lateritic, nutrient poor soils. We consider Worsley data sampled in 2019, with rehabilitation ages between 2–28 years old. Worsley’s 25 samples correspond to rehabilitation years: 1991 (n = 2), 1996 (n = 4), 1999 (n = 2), 2002 (n = 2), 2005 (n = 2), 2007 (n = 1), 2011 (n = 3), 2017 (n = 3), reference (n = 6).

Each soil sample had physico-chemical analyses performed at CSBP Laboratories (Perth, Western Australia) to quantify key soil abiotic variables as prescribed by AM protocols, including soil texture, organic carbon, ammonium, potassium, sulphur, calcium, pH, nitrate, phosphorous, and electrical conductivity.

### 2.2 eDNA sequencing, bioinformatics, and data preparation

DNA extraction, PCR and preliminary bioinformatic analyses were undertaken in accordance with AM workflows (Bissett et al., 2016; see SI Appendix, Supplementary Methods). From this workflow, denoised 16S rRNA gene amplicon sequence variant (ASV) level abundance data were produced for all minesites. Note, in this study ASVs are equivalent to zero radius OTUs (zOTUs). Further data preparation and analyses were largely undertaken in R version 4.0.3 (R-Core-Team, 2020) utilising the framework of the R phyloseq package (McMurdie & Holmes, 2013) to manage the datasets (see SI Appendix Supplementary Methods for number of sequences and ASVs studied in each minesite, initial data cleaning steps, and preparation of phylogenetic trees).

### 2.3 Data visualisation and statistical analyses

We visualised the sequence depth of samples using rarefaction curves (SI Appendix, Figure S2). We performed exploratory data analyses to visualise ASV alpha diversity, evenness, and relative abundance via heatmaps of phyla, classes, and orders in each minesite (SI Appendix, Supplementary Methods, Figures S3–S13). Alpha diversity and evenness were based on rarefied ASV abundances (as below), while relative abundances were computed using non-rarefied data. Further exploratory data analyses included preliminary visualisations of soil and landscape variables that associated with the soil bacterial community samples within each minesite (see SI Appendix Supplementary Methods, Supplementary Data, Figures S14–20).

To prepare for the computation of ‘standard’ ecological distance measures (as described by Gloor et al., 2017; e.g. Bray-Curtis, Jaccard, UniFrac), we normalised the sequence data for sampling effort by rarefying abundances of ASVs, and other taxonomic levels investigated (see below), to the minimum sample sequence depth within respective minesites (Huntly, n = 17,485 sequences; Eneabba, n = 10,142 sequences; Worsley, n = 54,122 sequences) using the *rarefy_even_depth()* function from R phyloseq.

To prepare ‘compositional’ data analysis distance measures we followed the recommendations of Gloor et al. (2017) and Quinn et al. (2019), using non-rarefied data. However, we took the pragmatic initial step of excluding taxa that contained zero counts in more than 90% of samples within each minesite, to help limit the potential for artefactual influences (as discussed later) to be introduced by the subsequent steps of zero replacement and centred log ratio transformation. Then, following Quinn et al. (2019) we used the default geometric Bayesian multiplicative model in the *cmultRepl()* function of the R zCompositions package (Palarea-Albaladejo & Martín-Fernández, 2015) to replace zeros with small numbers; before computing centred log ratio transformations using the *propr()* function from the R propr package (Quinn et al., 2017). Further steps are outlined in section 2.3.1 below.

We examined a range of alternative qualitative and quantitative beta diversity (i.e., distance or community dissimilarity) measures which were converted to similarity, to model rehabilitation trajectories and time to reach reference targets (as described further below). For the minesite with the largest number of samples (Huntly), we also investigated data pre-processing options of grouping by sequence similarity, taxonomic grouping, and excluding rare taxa. Details of the number of samples, taxa and sequences considered for all minesites, distance measures and data processing options (see below) are provided in the SI Appendix, Table S4.

#### 2.3.1 Comparison of alternative ecological and compositional similarity measures

For each minesite, we used the cleaned and rarefied ASV-level bacterial community data to derive standard ecological distance matrices using distance measures commonly employed in microbiota studies—i.e., Jaccard, Bray-Curtis, Unweighted UniFrac and Weighted UniFrac (Lozupone et al., 2007)—via the *vegdist()* function from the R vegan package (Oksanen et al., 2020). We also compared results from the Bray-Curtis measures with the compositional data analysis approach from computing Aitchison distances via *vegdist()* (i.e., these were derived from Euclidean distances between samples after centred log ratio transformation; Gloor et al., 2017). For each minesite, Bray-Curtis distances were visualised using non-metric multidimensional scaling (NMDS) ordination, while Aitchison distances were visualised using principal components analysis (PCA) (Gloor et al., 2017) (Figure 2). For the comparison between Bray-Curtis and Aitchison measures at Worsley we used the spatially filtered dataset which excluded the southernmost samples as described in section 2.3.6.

**FIGURE 2.**
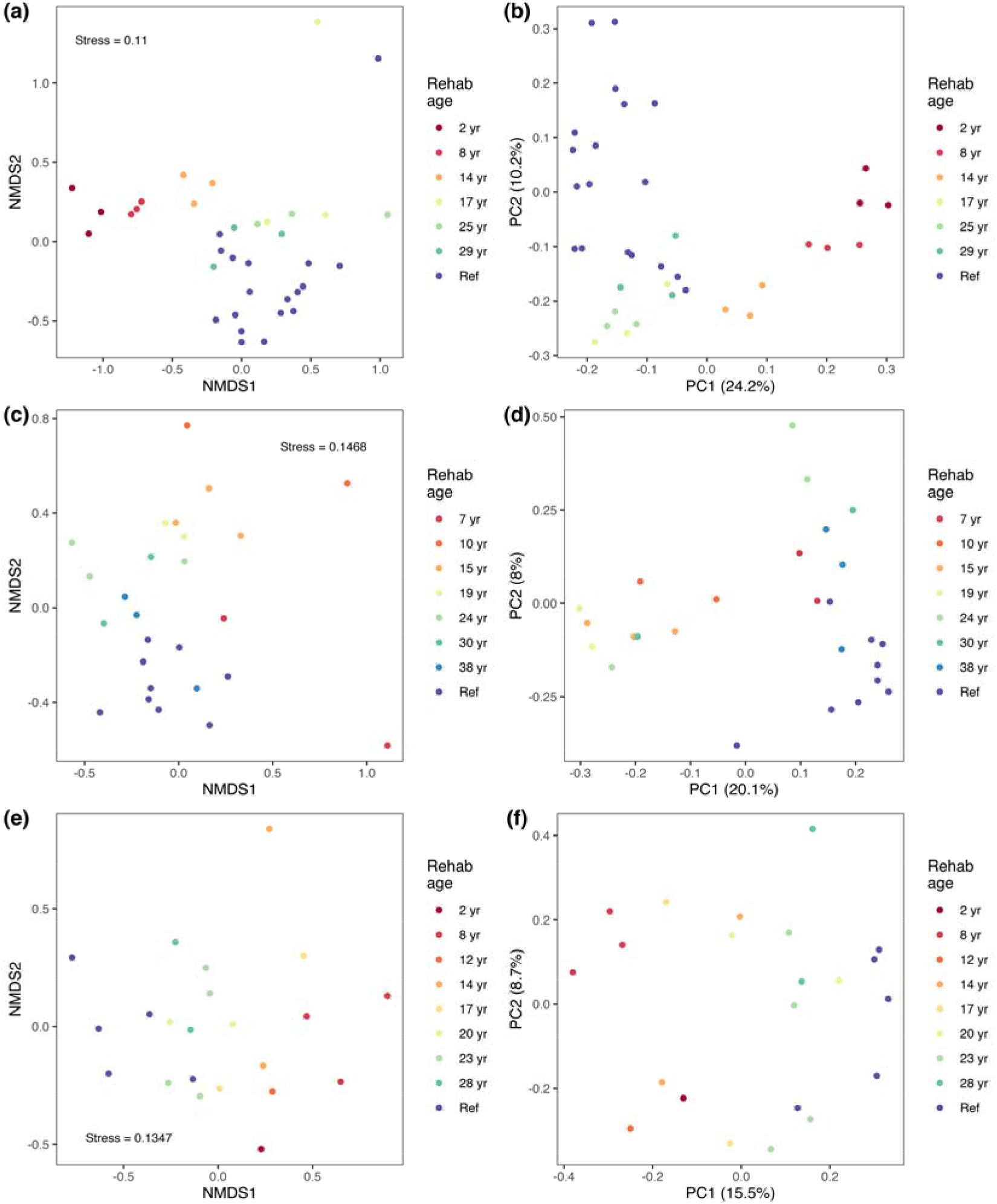
NMDS and PCA visualisations of differences in soil bacterial communities for: Huntly (n = 36 samples) using (a) Bray-Curtis distances (30,751 ASVs; 629,460 sequences) and (b) Aitchison distances (25,720 ASVs, 1,723,759 sequences); Eneabba (n = 26 samples) using (c) Bray-Curtis distances (27,115 ASVs; 263,692 sequences) and (d) Aitchison distances (24117 ASVs; 2,042,214 sequences); Worsley (excluding southernmost samples, n = 22 samples) using (e) Bray-Curtis distances (53404 ASVs; 1,190,684 sequences) and (f) Aitchison distances (43,598 ASVs; 1,782,724 sequences).

The rehabilitation trajectory analyses presented here were then derived from a subset of data contained in the above distance matrices. Specifically, only pairwise distances between samples and reference samples were considered (including distances among reference samples within minesites).

For standard measures (i.e. Bray-Curtis, Jaccard, Weighted UniFrac and Unweighted UniFrac) data were then expressed as percent similarity to reference values using:

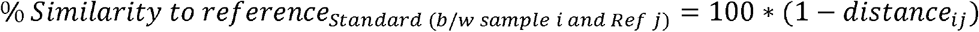

For the compositional Aitchison measures, similarity to reference was calculated using:

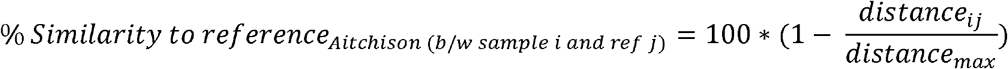

#### 2.3.2 Grouping by sequence similarity

For Huntly data only, separate R phyloseq objects were generated to represent soil bacterial community data with sequences clustered into 99%, 97%, 95%, and 90% identity OTUs (see SI Appendix, Supplementary Methods). For these analyses, OTUs were formed, abundance data were rarefied, and then Jaccard and Bray-Curtis distances and similarity to references were calculated.

#### 2.3.3 Taxonomic grouping

For Huntly data only, we examined the influence of taxonomic grouping (i.e., ASV, genus, family, order, class, and phylum) on the assessments of recovery. We also tested the influence of discarding versus retaining (at the next available classified grouping) taxa that were unclassified at each taxonomic rank, which we termed ‘pruned’ and ‘non-pruned’ data respectively. Grouping was undertaken using *tax_glom()*; and in ‘pruned’ datasets, unclassified taxa were removed using *prune_taxa()* from R phyloseq. For these analyses, taxa were grouped, abundance data were rarefied, then Jaccard and Bray-Curtis distances and similarity to references were calculated. Richness and evenness of sequences at the order, class and phylum level were also visualised based on rarefied data and plotted together with composite estimates within rehabilitation age groups from merged-sample bootstrap resampling (Liddicoat et al., 2019) (B=100).

#### 2.3.4 Excluding rare taxa

For Huntly data only, we examined the influence of excluding rare taxa, by considering all ASVs, then ASVs with >0.001 %, > 0.01%, and > 0.1% relative abundance within each minesite. For these analyses, ASVs with below the respective relative abundance threshold were removed, abundance data were rarefied, then Jaccard and Bray-Curtis distances and similarity to references were calculated.

#### 2.3.5 Rehabilitation trajectory modelling

The progress of rehabilitation was then visualised using boxplots and logarithmic models based on the similarity to reference data. Boxplots were generated from the series of similarity to reference data on the y-axis and increasing rehabilitation age on the x-axis, concluding with reference samples (e.g., Figure 3). Testing for differences in similarities to reference at each rehabilitation age (as visualised with boxplots) was performed using the Kruskal-Wallis rank sum test, followed by post-hoc Dunn tests for multiple comparisons, with Bonferroni adjusted threshold *P*-values. The multiple comparison testing used default two-sided *P*-values and alpha = 0.05 nominal level of significance.

**FIGURE 3.**
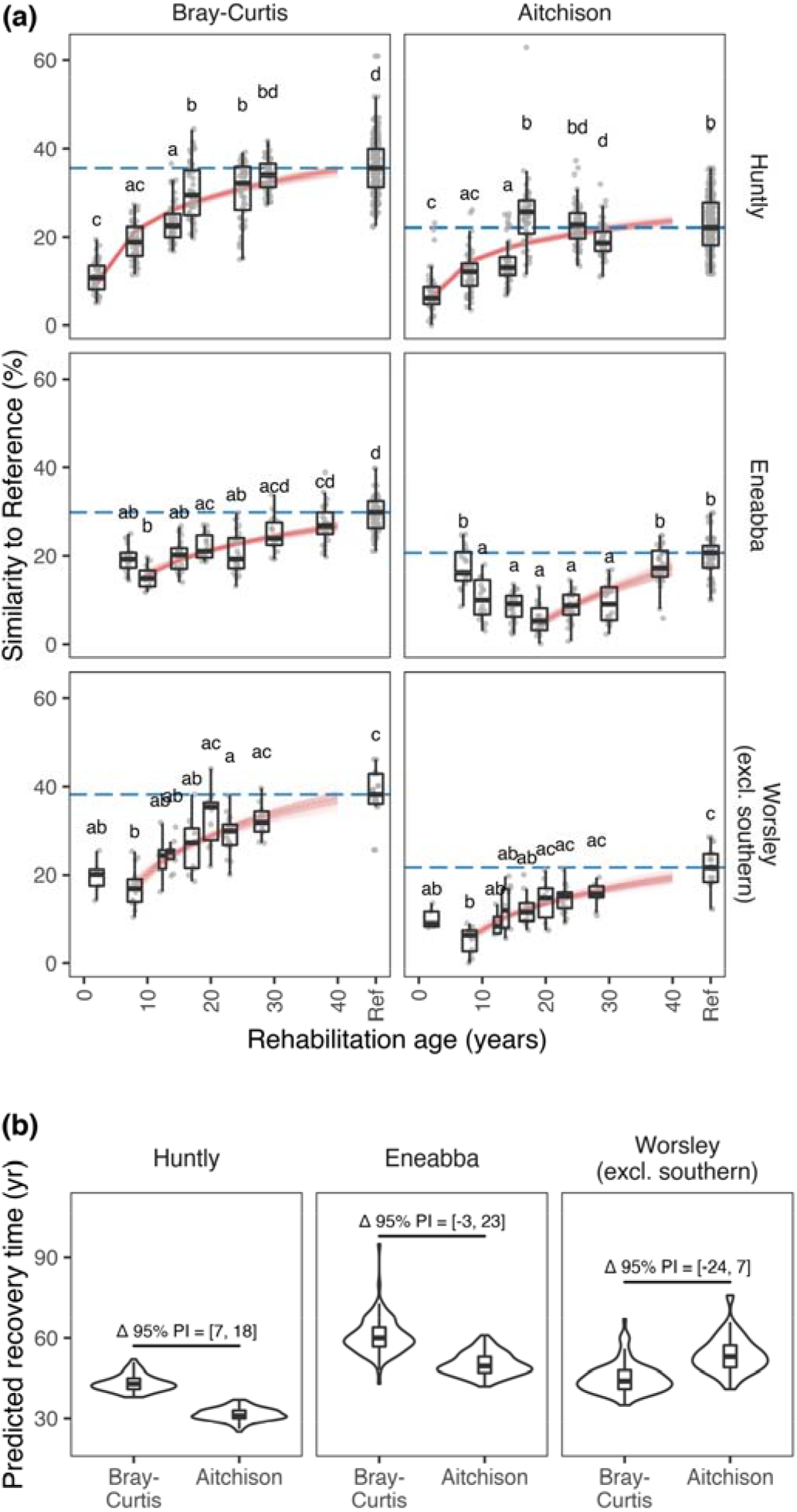
Modelled rehabilitation trajectories (a) and predicted recovery times (b) for Huntly, Eneabba, and Worsley (excluding southernmost samples) based on surface soil bacterial community similarity to reference data using Bray-Curtis and Aitchison measures. Plots are derived from the same data that underpin Figure 2. In (a), blue dotted lines denote the target median similarity among reference soils, and red lines represent logarithmic models for changing similarity to reference with rehabilitation age based on bootstrap resampling and modelling (B=100). Boxplots display the distribution of similarity to reference values across rehabilitation ages (groups not sharing a letter are significantly different). In (b), violin plots with boxplot inlays depict the distribution of recovery times from the 100 bootstrap model runs. Δ 95% prediction intervals (PI) indicate whether differences in recovery times predicted using alternative measures (Bray-Curtis versus Aitchison) are significantly different.

After observing the variation in similarity to reference values among references within each minesite (e.g., Figure 3), we defined rehabilitation targets for the purpose of this study as the median (= the central value) of among-reference similarities. This target median value varied by minesite, distance/similarity measure, and pre-processing option.

We predicted the time to reach a restoration target (= recovery time) by modelling the trend in similarity to reference with increasing rehabilitation age using bootstrapped (B = 100) logarithmic models. The median, 2.5^th^ and 97.5^th^ percentiles of predicted recovery time were evaluated. Our use of logarithmic models was consistent with the approach of Rydgren et al. (2019), except we used similarity not distance measures. Each iteration of the bootstrap involved random sampling with replacement from the available chronosequence similarity to reference data, excluding outliers identified via the *boxplot()* function in base R, and developing a predictive logarithmic model for similarity to reference out to a maximum rehabilitation age of 500 years, or until the target was reached. Models that failed to reach the target were reported with a prediction time of ‘>500 years’. Rectangular hyperbola and negative exponential models were also trialled but were abandoned after many cases failed to produce model fits. During our early analyses, we also uncovered example data that highlighted a distorting influence on our trajectory (and recovery time) modelling that appeared to be consistent with the application of ‘direct return’ soils in young rehabilitation sites. Specifically, this soil material was more similar to references than older rehabilitation sites. Including these samples with elevated similarity to references in the logarithmic modelling appeared to bias models towards flatter, longer trajectories of recovery. Therefore, to reflect the likely onset of recovery towards reference states we decided to only commence logarithmic models (via our automated modelling algorithm) from the youngest rehabilitation age group that had a next older group with increased median similarity to references. As discussed later, this check on model commencement was designed to avoid likely distortions in the modelling of recovery, in particular, due to potential biological inertia in direct return soils (Janzen, 2016).

#### 2.3.6 Exploring spatial autocorrelation

To explore the influence of spatial autocorrelation on our trajectory analyses, we produced variogram-like plots using Bray-Curtis ecological distances (between samples and references) on the y-axis, and geographic distances (between samples and references) on the x-axis. Each rehabilitation age group was modelled as a second-order polynomial, allowing the possible expression of curvilinear trendlines that mimicked variogram-like relationships (i.e., increasing then flattening). Assuming reference curves offered a natural baseline trend for spatial autocorrelation within each minesite environment, we applied a ‘correction’ to the curvilinear trendline for each rehabilitation age group by calculating the difference in mean-centred model curves (= rehabilitation age group minus reference), such that ‘corrected’ data for rehabilitation age groups expressed the same ecological distance-geographic distance curvilinear trend as seen for references (see SI Appendix Supplementary Methods for further details of the rationale and approach for this preliminary analysis). Rehabilitation trajectories and predicted recovery times were compared between ‘original’ and ‘corrected’ data, for the Bray-Curtis similarities. For the Worsley minesite, a filtered dataset, and corresponding correction, were also prepared which excluded the three southernmost samples (i.e., two 2-year old samples and an adjacent reference), which were geographically separate from the other Worsley samples (see Figure 1, and SI Appendix Table S3).

## 3. RESULTS

### 3.1 General findings

We found remarkable variability among reference samples within each minesite (Figure 3; SI Appendix, Table S5, Figures S21, S23–S25). Median among-reference similarities ranged from <20% to >95% across all measures, and between approximately 30–40% for Bray-Curtis measures, with variation depending on the specific distance measure, pre-processing option, and minesite. All rehabilitation trajectory plots indicated recovery, displaying the general pattern of increasing similarity to references with increasing rehabilitation age (Figure 3; SI Appendix, Figures S21, S23–S25), although the logarithmic models and predicted recovery times varied with distance measures, pre-processing and minesite.

### 3.2 Alternative ecological and compositional measures

Despite some differences in the expression of rehabilitation trajectories using Bray-Curtis versus Aitchison measures at Huntly, Eneabba, and Worsley (excluding southernmost samples), these measures produced comparable predictions for recovery time within each minesite (Figure 3; SI Appendix, Table S6). At Huntly, predicted recovery times differed by around 12 years, with a median recovery of 43 years for Bray-Curtis measures and 31 years for Aitchison measures. At Eneabba, the median Bray-Curtis recovery time was 60 years, while the median Aitchison recovery time was 50 years, however due to the spread of model outcomes, predictions from these measures were not significantly different (i.e., Δ 95% interval contains zero; Figure 3). Similarly, at Worsley (excluding southernmost samples), the median Bray-Curtis recovery time was 44 years, while the median Aitchison recovery time was 53 years, however predictions from these measures were not significantly different (i.e., Δ 95% interval contains zero; Figure 3).

Among standard measures we found a general increase in similarity to reference values across the ecological measures, from Jaccard (generally lowest similarities), Bray-Curtis, Unweighted UniFrac, to Weighted UniFrac (generally highest similarities) (SI Appendix, Figure S21, Table S5). The greatest y-axis span, and therefore greatest sensitivity to detect change, in similarity to reference values between the youngest rehabilitation ages and references occurred with Bray-Curtis measures (SI Appendix, Figure S21). The smallest span (or flattest curves) in similarity to reference values between the youngest rehabilitation ages and references occurred with Weighted UniFrac measures.

Except for the Unweighted Unifrac result at Huntly, Jaccard measures generally returned the longest predicted recovery times, followed by reduced or similar recovery times predicted using Bray-Curtis, Unweighted Unifrac and Weighted UniFrac measures (SI Appendix, Figure S22, Table S6). Low sample sizes (and corresponding low numbers of distance measures) represent a limitation in our analysis, and the ecologically-distant samples in the 17-year and 25-year rehabilitation age group at Huntly (Figure 2a) are likely contributing to the reduced similarity and longer rehabilitation trajectory in Unweighted UniFrac data. These 17-year and 25-year rehabilitation age group data at Huntly express reduced alpha diversity and evenness compared to other samples, however reasons for this are unclear (SI Appendix, Figures S3–S4).

### 3.3 Grouping by sequence similarity (Huntly only)

Grouping by sequence similarity resulted in progressive overall shifts towards increasing similarity to reference values from ASV-level (generally lowest similarities), 99%, 97%, 95%, to 90%-identity clustered OTUs (generally highest similarities) (SI Appendix, Figure S23). Predicted recovery times with more broadly clustered OTUs followed continuous and seemingly predictable patterns of: (i) increasing recovery times with Jaccard measures, and (ii) decreasing to steadying recovery times with Bray-Curtis measures (SI Appendix, Figure S26a, Table S6).

### 3.4 Taxonomic grouping (Huntly only)

Moving from ASV to genus-level data resulted in a pronounced shift towards increasing similarity to reference, with similar although somewhat flatter rehabilitation trajectory curves at higher taxonomic groupings (SI Appendix, Figure S24). Visually, there appeared to be little effect on the rehabilitation trajectory plots from pruning unclassified taxa (SI Appendix, Figure S24). Using Jaccard measures, moving from ASV-level to grouping at genus-level or higher groupings dramatically increased predicted recovery times, compared to other measures (SI Appendix, Figure S26b, Table S6). Also, pruning of unclassified groups reduced the smoothness or continuity in Jaccard-predicted recovery times (SI Appendix, Figure S26b). Using Bray-Curtis measures, we found a non-linear pattern of recovery times across the taxonomic groupings, with shorter times to reach the target in genus, family, and order-level groups, and longer recovery times in other groupings (SI Appendix, Figure S26b; see SI Appendix, Figures S5–S13 for relative abundances of order, class, and phylum-level taxa for each minesite). Richness and evenness of bacterial communities varied across rehabilitation age groups and taxonomic groupings (e.g., data for phylum, class, and order-level are shown in SI Appendix, Figure S27), which may help explain the somewhat erratic results from taxonomic grouping.

### 3.5 Excluding rare taxa (Huntly only)

Removing rare taxa to the point of retaining ASVs with >0.01% relative abundance produced results from the Jaccard analysis that appeared to mimic results from the Bray-Curtis analysis (SI Appendix, Figure S25). When only more common ASVs with >0.1% relative abundance were retained, both the Jaccard and Bray-Curtis results appeared to reflect over-simplified communities, resulting in shorter predicted recovery times. However, including only ASVs with >0.001% relative abundance resulted in a dataset with approximately 60% of the original taxa and 95.8% of total sequences after rarefying (i.e., 17,941 compared to 30,751 ASVs and 603,072 compared to 629,460 sequences; SI Appendix, Table S4) and produced only a small increase in predicted recovery times for both Jaccard and Bray-Curtis measures (SI Appendix, Figure S26c, Table S6).

### 3.6 Correcting for spatial autocorrelation

We modelled the slope-trends of the relationships between ecological distance to references and geographic distance to references, within rehabilitation age classes, for each of the minesites using Bray-Curtis measures (see SI Appendix, Huntly and Eneabba: Figures S28–S29; Worsley: Figure 4). We also applied a ‘correction’ for the spatial autocorrelation, such that rehabilitation age groups were adjusted to display the same ecological-geographic slope trend as found in references (refer to the ‘c’ panels in SI Appendix, Figures S28–S29; Figure 4). Figure 4d–f also includes the Worsley ‘filtered’ dataset and corresponding correction, where the three southernmost geographically separate samples were excluded. Rehabilitation trajectory plots, and predicted recovery times, using corrected data were compared to the original uncorrected data (see Figure 5 and SI Appendix, Table S6).

**FIGURE 4.**
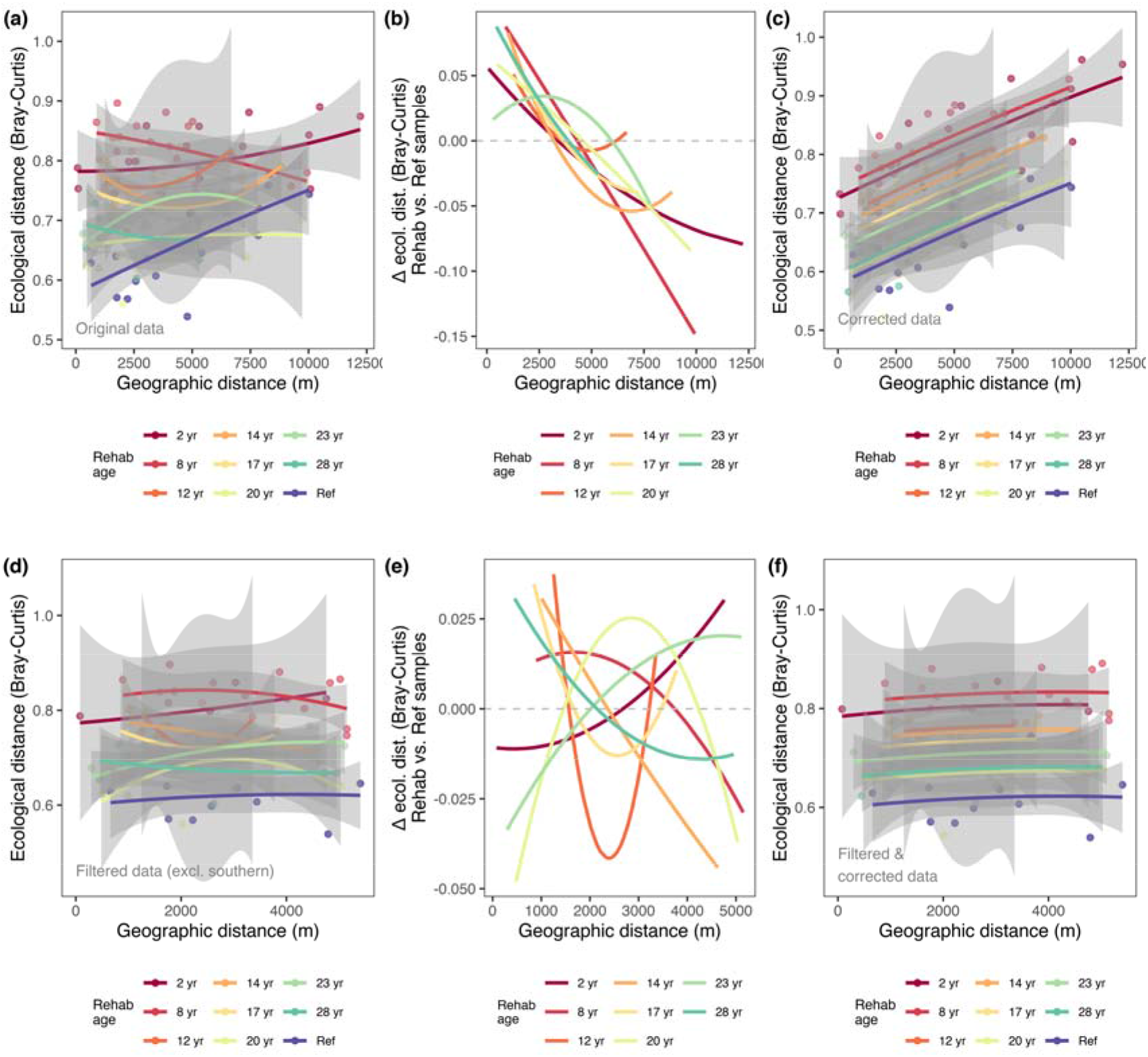
Exploring spatial autocorrelation in the Worsley (a–c) and filtered Worsley (excluding southernmost samples) (d–f) datasets, based on Bray-Curtis distance measures. (a, d) Ecological distance to reference versus geographic distance to reference for rehabilitation age groups. (b, e) Mean-centred difference in ecological distance to reference between rehabilitation age groups and among references. (c, f) Corrected ecological distance to reference versus geographic distance to reference for rehabilitation age groups, to match the slope-trend of ecological to geographic distances as found among references.

**FIGURE 5.**
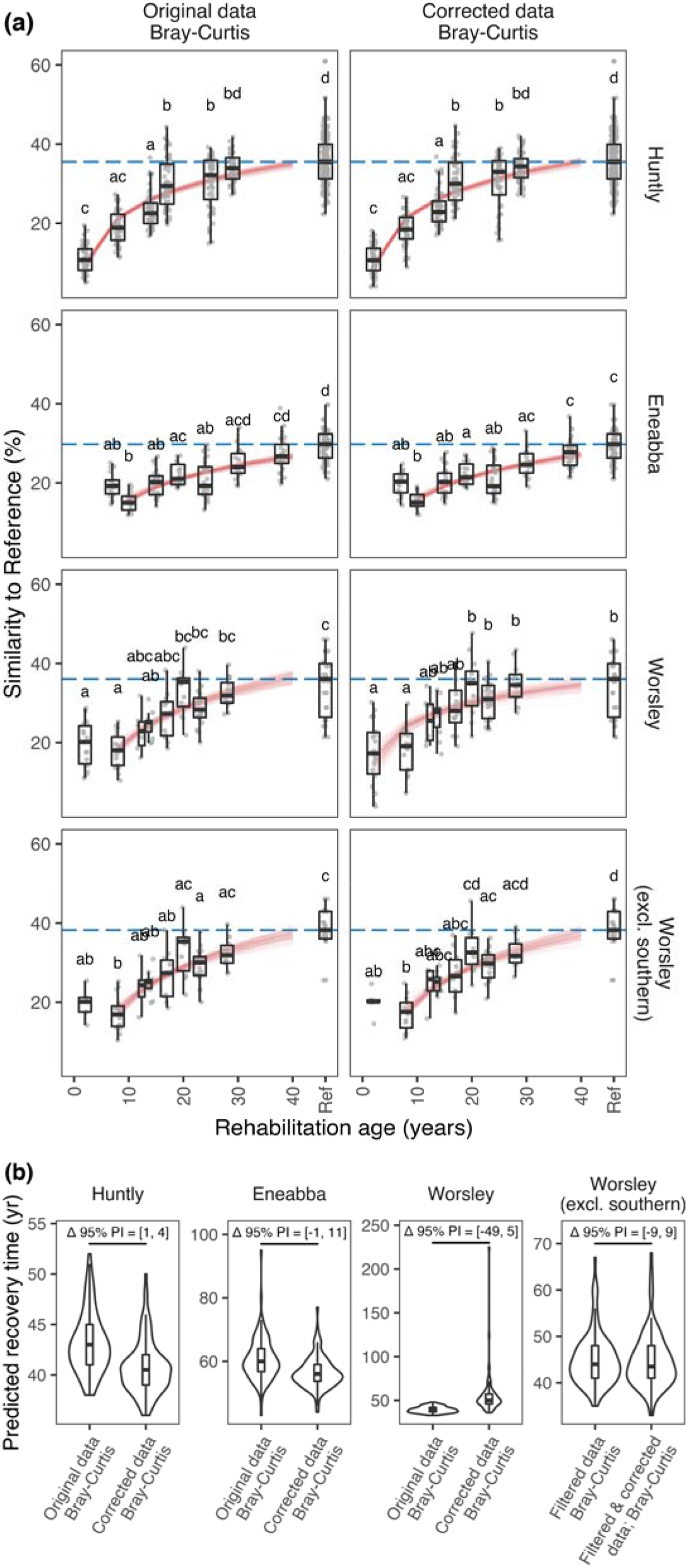
Modelled rehabilitation trajectories (a) and predicted recovery times (b) for Huntly, Eneabba, Worsley, and Worsley (excluding southernmost samples) based on surface soil bacterial community similarity to reference data using Bray-Curtis measures, with and without correction for spatial autocorrelation. Other features are as described in Figure 3.

Worsley displayed a strong ecological distance-geographic distance trend in among-reference data indicating excessive spatial autocorrelation (note the upward sloping ‘Ref’ line in Figure 4a), and the greatest divergence of all the minesites in predicted recovery times between original and corrected data (Figure 5; SI Appendix, Table S6). Notably, the spatial autocorrelation correction at Worsley caused such an adjustment in similarity to reference values that the youngest rehabilitation age group was included in the logarithmic trajectory models in the corrected data, but not in the original data. However, with exclusion of the southernmost Worsley samples (i.e., the filtered dataset), the signal of spatial autocorrelation disappeared (i.e., absence of upward sloping lines in Figure 4d, f) and predicted recovery times for filtered and filtered-corrected data displayed almost identical distributions (Figure 5; SI Appendix, Table S6).

## 4. DISCUSSION

### 4.1 Standard vs. compositional data analysis

Our rehabilitation trajectory models produced comparable predictions for recovery times using Bray-Curtis (standard) and Aitchison (compositional) measures. At Huntly, Eneabba, and Worsley (excluding southernmost samples) median recovery times differed by around a decade (i.e., 43 vs. 31 years, 60 vs. 50 years, 44 vs. 53 years respectively), however for two out of three minesites the distribution of bootstrap model predicted recovery times was not significantly different. We suspect that both Bray-Curtis (standard) and Aitchison (compositional) measures will provide slightly different perspectives to the trajectory modelling (discussed below), while neither method is perfect.

Compositional data analysis has been recently promoted as a more robust approach for analysing microbiome datasets (Gloor et al., 2017; Quinn et al., 2019), however it is not without limitations particularly for sparse datasets (containing many zeros), and where low sequence counts are commonly encountered (Lovell et al., 2020). In particular, replacement of zeros with small positive numbers has potential to cause distortions in data that will affect the relative abundance of small counts to a greater degree than large counts (Lovell et al., 2020). Distortions in data due to zero replacement are also increased where there are large numbers of zeros present (Martín-Fernández et al., 2015). Therefore, our approach to exclude taxa that contained zero counts in more than 90% of samples within each minesite represents a compromise between losing representation of less common taxa and potentially introducing spurious log ratio abundance patterns within the compositional data analysis. Following log ratio analysis, only the relative information is of interest; for example, counts of 1,2,3 become equivalent to counts of 100, 200, 300. However, advocates of these approaches have suggested that it is up to the analyst to decide whether the relative, rather than the absolute, structure of the parts is of primary interest (Martín-Fernández et al., 2015). Also, the replacement of absolute zeros (representing true absences; as opposed to zeros due to rounding or resulting from insufficiently large samples) with small numbers is potentially inappropriate (Martín-Fernández et al., 2015), and creates a theoretical challenge to performing log ratio analyses on soil microbiota data from diverse environments where many absolute zeros (true absences) are likely.

### 4.2 Alternative standard ecological measures

Bray-Curtis measures produced the greatest range in similarity values between young rehabilitation and reference samples, and therefore are likely to offer the greatest sensitivity to quantify the progress of recovery of soil bacterial communities towards reference states. In contrast, Weighted UniFrac offered limited sensitivity to detect changes with rehabilitation age (i.e., shallow trajectory curves) and may result in under-prediction of recovery times. Low variation in Weighted Unifrac similarities likely reflects a level of consistency of high proportions of somewhat closely related organisms across the samples. Jaccard distances represent the proportion of unshared taxa out of the total number of taxa recorded in two groups (Anderson et al., 2006). Unweighted UniFrac uses phylogenetic information and calculates the fraction of the branch length in a phylogenetic tree that leads to descendants in either, but not both, of the two communities (Lozupone et al., 2007). These qualitative measures reflect the survival and presence of taxa (Jaccard) and related lineages (Unweighted UniFrac), where loss of sequences may reflect extreme or limiting environmental conditions (e.g., soil abiotic factors) or limited geographic distribution. Meanwhile, Bray-Curtis and Weighted UniFrac measures emphasise abundant organisms. Similarity to reference generally increased with increasing abundances of shared taxa for Bray-Curtis, and shared lineages of related sequences for Weighted UniFrac. The quantitative measures often reflect the growth or decline of certain organisms due to factors such as nutrient availability and variation in environmental conditions (Lozupone et al., 2007).

### 4.3 Grouping by sequence similarity

Grouping near identical sequences will reduce the denominator used in calculating Jaccard distances. For a given number of unshared taxa between samples, using broader OTU clusters will make the proportion of unshared taxa (compared to all taxa) larger when there are a smaller number of total taxa present. Our data suggest this shifting Jaccard calculation can impact some samples strongly (e.g., note the 17-year age group in SI Appendix, Figure S23) resulting in a gradual increase in predicted recovery times with broader (reduced identity threshold) OTU clusters. On the other hand, broader OTU clusters will aggregate some sequences into already large groups and will tend to further emphasise abundant groups. Consequently, our Bray-Curtis data suggest broader OTU clustering will make the target similarity easier to reach and predicted recovery times reduced accordingly.

### 4.4 Taxonomic grouping

We do not recommend grouping 16S rRNA data by taxonomy to quantify recovery in soil bacterial communities due to the erratic behaviour of predicted recovery times.

### 4.5 Excluding rare taxa

We show that filtering out of rare taxa to a limited extent (>0.001% relative sequence abundance) produces a relatively small increase in predicted recovery times for both Jaccard and Bray-Curtis measures. In our case, this filtering removed many ASVs but only a low percentage of total sequences. Interestingly, this low level of exclusion of rare taxa does not appear to moderate the assessment by producing reduced recovery times. At the low level of exclusion, our analysis using rarefied data and similarity to reference measures may help mitigate some of the impacts and concerns of removal of rare sequences experienced elsewhere (e.g., Schloss, 2020). This raises the prospect to reduce sequencing depth, and potential for shifting investment towards more robust assessments that incorporate a larger number of samples with reduced sequencing depth and cost per sample.

### 4.6 Influence of ‘direct return’ soils in young rehabilitation sites

For reasons discussed here and below, we suggest it is prudent for these similarity to reference trajectory assessments to exclude young rehabilitation sites with ‘direct return’ soils that display elevated similarity to reference—as we implemented in our automated trajectory modelling algorithm. In earlier preliminary work at Eneabba and Worsley, we observed that the inclusion of young rehabilitation samples that were overly similar to references resulted in seemingly biased, longer predictions of recovery time. The industry best practice of ‘direct return’ of topsoil to new rehabilitation sites is based on objectives to minimise soil degradation and expedite ecosystem recovery. However, our use of monotonic logarithmic models applied to a data series that contains young rehabilitation sites with elevated similarity to reference values, followed by older sites with reduced similarity to reference values, results in the seemingly perverse outcome of a flatter, longer modelled trajectory of recovery. The enhanced ecological similarity to reference in young rehabilitation sites with ‘direct return’ soils reflects a biological inertia, or temporary carryover effect, from unmined areas where the soils originate, and confounds the relationship between soil microbiota development and rehabilitation age. For ‘direct return’ soils, we speculate the time taken for local influences to become dominant in shaping the resident microbiota may be in the order of 1-10 years, varying on a case-by-case basis, e.g., due to soil factors including organic matter and clay content, as well as the magnitude of environmental influences. Soil microbiota will be shaped by influences including local rainfall, temperature, aspect, soil water availability and transport (e.g., run-on, lateral flow), and vegetation communities via plant-soil feedbacks. Existing deeper soil and substrate may also influence rehabilitation surface soils via upward movement of water, nutrients, and some microbiota through mechanisms including: hydraulic redistribution by plant root systems (Neumann & Cardon, 2012); potential microbiota uptake and transfer via xylem into the phyllosphere (Deyett & Rolshausen, 2019; Fausto et al., 2018) and subsequent leaf litter; and capillary rise in heavier textured soils under conditions of soil water evaporation. Other factors affecting the similarity to reference of direct return soils include their source location (are they taken from sites that are generally closer to other reference sites or adjacent to rehabilitation sites?), the depth of fresh topsoil applied, the condition of subsurface layers (e.g., fresh vs. stockpiled), and the depth and method of tillage or mixing of the soil surface and subsurface layers following soil return. Our approach to automate the commencement of logarithmic models once there is at least an initial increase in similarity to reference values provides an objective approach to help overcome the potential model-biasing effect of biological inertia that is found in some direct return soils.

### 4.7 Spatial autocorrelation

We found signals of excessive spatial autocorrelation where strong slopes were detected in plots of ecological distance to reference versus geographic distance to reference, and where substantial differences were detected in the logarithmic models and/or predicted recovery times between original and corrected datasets. Excluding geographic outliers in the filtered Worsley analysis also removed a clear spatial autocorrelation signal in the data, which indicates the importance of sampling designs. If rehabilitation sites reflect environmental settings or imported soils that are overly similar or dissimilar to references (i.e., different to natural background rates of spatial autocorrelation), this may unduly bias predicted recovery times. Where possible, we recommend a sampling approach that resembles the approach used at Huntly, where each reference site was spatially paired with an adjacent rehabilitation site. This approach helps capture variation among references (within a given minesite) relevant to the broader range of rehabilitation sites; and provided there is adequate spatial replication and geographic outliers are avoided, then undue influence from spatial autocorrelation should be avoided.

Our analysis of spatial autocorrelation should be viewed as introductory and illustrative. For ‘direct return’ soils at young rehabilitation sites, our approach is deficient because we do not account for their previous location. Although, we anticipate localised influences would dominate the shaping of resident soil microbiota in rehabilitation sites after a few years, as discussed above.

Plant-soil-microbiota feedbacks represent a complicating factor for disentangling effects of soil abiotic condition, rehabilitation age, and residual/unexplainable spatial autocorrelation in restoration chronosequence studies. This is because chronosequence studies (which presume a ‘space-for-time’ proxy relationship between treatments and outcomes) typically do not collect sufficient data to determine whether soil conditions have influenced rehabilitation outcomes, plants have conditioned soils, or both situations have occurred. Studies that have considered plant-soil feedbacks in restored Jarrah forest (Huntly) sites have shown differential correlative effects of rehabilitated soil biotic and abiotic properties (Orozco-Aceves et al., 2017). Also, plant-soil feedbacks behave differently in unmined versus rehabilitated soils (Orozco-Aceves et al., 2015). Further work is required to build understanding of this topic (e.g., via longitudinal studies).

### 4.8 Other limitations

There are important limitations in our study, in addition to those already discussed. The robustness of our study would be improved with more samples per minesite to help better capture minesite-wide variation. We did not consider soil microbiota patterns at depth, which are also important. Also, major changes to rehabilitation practices over time will disrupt the ‘space-for-time’ substitutive modelling approach that is relied upon in chronosequence studies such as ours. For any restoration chronosequence study careful sample selection is required to avoid confounding factors as much as possible (Walker et al., 2010). There are potential limitations in our study associated with the phylogenetic trees we used to generate UniFrac distances (see SI Appendix, Supplementary Methods for details). Tree-building often represents a compromise between accuracy in representing phylogenetic relationships and computing time, and it was beyond the scope of our study to test the sensitivity of our UniFrac-based analyses to the quality of trees used. We used logarithmic models which assume a monotonic recovery function, however other models that account for variable trends over time, and varying success for different rehabilitation techniques or sites, may offer improved estimates of recovery time. We suggest these limitations should be investigated in future studies.

## 5. CONCLUSIONS

We provide a proof-of-concept demonstration of an innovative, chronosequence-based, similarity to reference trajectory assessment method, to quantitatively track progress in soil microbiota with post-mining rehabilitation. Through incorporating microbiota survey data from multiple reference sites of varying character, we revealed substantial variation among reference ecosystems within each minesite that can inform realistic rehabilitation targets. Our method reduces the complexity associated with microbiota data and enables prediction of recovery time to reach reference-based targets with explicit inclusion of uncertainty in assessments. Also, the use of soil microbiota data provides another line of evidence, which in conjunction with wider minesite information, could assist in the examination of potential impediments to the progress of rehabilitation, thereby helping to inform adaptive management. From our investigations, we recommend using ASV-level Bray-Curtis similarities which appear to offer a relatively sensitive and stable basis for modelling rehabilitation trajectories. We recommend wherever possible to maximise sample sizes, employ spatial pairing of reference and rehabilitation sites, and to exclude geographically-distant, non-representative sampling areas. We used an automated modelling routine to exclude young rehabilitation sites with ‘direct return’ soils that displayed elevated similarity to reference values, which would have biased the trajectory modelling. Further fine-tuning to identify possible minor reductions in sequencing depths (eliminating some rare taxa) offers promise to reduce per sample costs, enabling investment in more samples, to help deliver more robust assessments. This work represents an important step towards a reduced-complexity microbiota-based monitoring and evaluation framework consistent with many best practice principles for setting, monitoring and managing towards mine completion criteria recommended by (Manero et al., 2021). We anticipate that our approach could be expanded to other eDNA sequence-based survey data (e.g., fungal ITS and eukaryote 18S rRNA marker genes, functional potential from shotgun metagenomic data), and may have application in wider contexts where there is interest in monitoring restorative processes that facilitate a shift in microbiota towards reference states.

## ACKNOWLEDGEMENTS

We acknowledge the contribution of the Australian Microbiome consortium in the generation of data used in this publication. The Australian Microbiome is supported by funding from Bioplatforms Australia and the Integrated Marine Observing System (IMOS) through the Australian Government’s National Collaborative Research Infrastructure Strategy (NCRIS), Parks Australia through the Bush Blitz program funded by the Australian Government and BHP, and the CSIRO. This research was also supported by the Australian Research Council (LP190100051).

## AUTHOR CONTRIBUTIONS

CL, SLK, MT and MFB conceived the ideas and designed the study; SLK, RJB, LCD, PB, MPD, AG collected the data; CL, SLK, AB, MFB analysed and interpreted the data with contributions from all authors; CL led the writing of the manuscript. All authors contributed critically to the drafts and gave final approval for publication.

## DATA AVAILABILITY STATEMENT

Data and code are available at: https://data.bioplatforms.com/organization/about/australian-microbiome and https://github.com/liddic/resto_traj

