## Supporting Information for "Next generation restoration metrics: Using soil eDNA bacterial community data to measure trajectories towards rehabilitation targets"

**for the article:**

This Supporting Information file includes:

Supplementary Methods
Supplementary Data
Figures S1–S29
Tables S1–S6
Supplementary References

Data and code availability:

Data used in this study are available from:
https://data.bioplatforms.com/organization/about/australian-microbiome
(refer to sample identifiers in Table S1–S3)

Code used in this study is available from:
https://github.com/liddic/resto_traj

### Supplementary Methods

#### Minesite descriptions

#### (i) Huntly

Site background information is sourced from the Honours thesis of (Ducki 2020).

Alcoa's Huntly bauxite minesite is located approximately 100 km south-east of Perth, Western Australia. The site occurs in highly biodiverse 'open forest' (Specht, Roe & Boughton 1974), with dominant overstorey species of jarrah (*Eucalyptus marginata*) and marri (*Corymbia calophylla*) trees that grow upward of 30-metres, mid-storey species including she-oak (*Allocasuarina fraseriana*), banksia (*Banksia grandis*), and grasstree (*Xanthorrhoea preissii*), and diverse understorey of woody shrubs and herbaceous perennials (Bell & Heddle 1989). The trees and shrubs are sclerophyllous, deep rooted, and adapted to the Mediterranean climate and fire regimes (Bell 2001). Fire represents a frequent disturbance in the jarrah forest, and defines characteristics of vegetation community succession, therefore knowledge of vegetation recovery in response to fire has helped to inform ecological restoration activities (Koch & Ward 1994; Norman *et al.* 2006). Restoration processes undertaken at Huntly are outlined in (Koch 2007).

The area is underlain by deeply weathered lateritic, nutrient deficient soils (Churchward & Dimmock 1989), with surface soils typically comprising acidic, yellow-mottled ironstone gravels and sands. Many plant species have evolved specialised root-microbial associations due to the region's infertile soils, promoting greater acquisition of limited nutrients (Brundrett & Abbott 1991). Soil microbial communities play a significant role in the decomposition of leaf litter and soil nutrient cycling, which occurs slowly in the jarrah forest (Jasper 2007). Supplementary fertiliser application such as di-ammonium phosphate (DAP) has been a common component of rehabilitation practices in the jarrah forest (Koch 2007). However, fertiliser rates have been reduced progressively since 1991 (Standish *et al.* 2015).

The post-mining chronosequence sites sampled in 2016 capture rehabilitation ages from 2–29 years. Due to the sequence of extraction and rehabilitation activities undertaken at Alcoa there is a general progression in age from older sites to the south, and younger sites to the north. A feature of the Huntly sampling is the pairwise comparisons between rehabilitation sites and adjacent reference forest.

#### (ii) Eneabba

Site background information is sourced from the Honours thesis of (Borrett 2020).

Iluka Resource's Eneabba mineral sands minesite is located approximately 280km north of Perth, Western Australia. Eneabba has a Mediterranean-type climate, and vegetation is adapted to regular fire disturbance events (Enright *et al.* 2012). The vegetation is known as kwongan sandplain heath, which consists of low shrubland on undulating infertile siliceous sandplains (Enright *et al.* 2012). Vegetation communities feature perennial woody species from Proteaceae, Myrtaceae, and Fabaceae (Mucina *et al.* 2014). Kwongan vegetation is highly diverse and endemic, and is seen as one of the most diverse vegetation types of all Mediterranean-type ecosystems (Cowling *et al.* 1996).

The study location is underlain by geologically old and weathered soils and regolith (Mucina *et al.* 2014), resulting in soils that are extremely nutrient poor. In the silicious sandy soils of the area, phosphorus is considered the most limiting nutrient (Enright & Lamont 1992). Vegetation height in the kwongan heath also reflects varying access to soil moisture, where low heaths on shallow sands have less water than taller vegetation on dune rises (Enright & Lamont 1992).

Sampling data from 2019 captured rehabilitation ages from 7–38 years, while native non-mined reference samples were also gathered from areas bordering the mining operations. The different rehabilitation ages encompassed a considerable amount of variability in topsoil storage time and mulching practices.

#### (iii) <u>Worsley</u>

Site background information is sourced from the Honours thesis of (Peddle 2020).

South32's Worsley Alumina bauxite minesite is located approximately 150 km south of Perth, Western Australia, where bauxite has been mined since 1984. The Worsley mine is located in the northern Jarrah (*Eucalyptus marginata*) forest within the Southwest Australian Floristic Region, an international biodiversity hotspot. The northern Jarrah forest is a dry sclerophyllous 'open forest' (Specht, Roe & Boughton 1974) dominated by jarrah (*Eucalyptus marginata*) and marri (*Corymbia calophylla*) trees with an understory dominated by vegetation from the Fabaceae, Asteraceae, Proteaceae, and Myrtaceae families (Koch & Samsa 2007). The site has a Mediterranean climate and lateritic, nutrient poor soils (Tibbett *et al.* 2020).

Samples were collected in October and December 2019 and captured rehabilitation ages from 2–28 years old, with a randomised selection of rehabilitated and reference sites across the minesite. Six replicates of uncleared, reference sites were selected across the minesite to compare with the rehabilitated sites.

### <u>Overview of rehabilitation practices</u>

The mining process involves removal of all vegetation and topsoil, and overburden is stripped away to access the ore. Topsoil storage is generally limited to around six months or, alternatively where possible, is directly transferred from an active mining area to a rehabilitation area (a process termed 'direct return'). Limiting topsoil storage times represents industry best-practice to help limit degradation processes on soil biotic and abiotic properties (Golos & Dixon 2014). Following ore extraction, mined areas are contoured to mimic surrounding topography, and then a cover of subsoil and topsoil is applied, before being furrowed and seeded with a mix of local native plant species. More detailed information on typical practices is available from (Tibbett 2010).

### Soil sampling protocols

Soil sampling followed procedures specified by the Biomes of Australian Soil Environments (BASE; Bissett *et al.* 2016) (https://www.australianmicrobiome.com/protocols/). Briefly, at each sampling 'site' a 25 x 25 m plot was identified and GPS coordinates were taken for the southwest corner. After scraping away surface leaf litter, surface (0–10 cm) soils were collected using sterilised equipment from 9–10 subsamples chosen to represent site heterogeneity across the 25 x 25 m plot. (Subsurface 20–30 cm subsamples were also taken however they were not considered in this study.) The subsamples were pooled into a sterile plastic bag, and then homogenised. From each pooled/composite sample, a 500 g subsample of soil was taken for chemical analysis and a 50 mL subsample for DNA extraction. Chemical analyses were performed at CSBP Laboratories (Perth, Western Australia) quantifying soil organic carbon, ammonium, potassium, sulphur, calcium, pH, nitrate, phosphorous, and electrical conductivity. The 50 mL sample was frozen on-site and sent packed on dry ice to the Australian Genome Research Facility (AGRF) in Adelaide, South Australia for DNA extraction (described below).

### DNA extraction, PCR-amplification and preliminary bioinformatic analysis

DNA was extracted from each sample in triplicate using the Qiagen DNeasy Powerlyzer Powersoil Kit following manufacturer's instructions and quantified using fluorometric nucleic acid quantification. Soil bacterial 16S rRNA was amplified using the 27F (Lane 1991; 5'-AGAGTTTGATCMTGGCTCA-3') and 519R (Lane *et al.* 1985; 5'-GWATTACCGCGGCKGCTG-3') primer set before sequencing (300bp PE) on the Illumina MiSeq platform. Sequence data used for this work was generated by the Australian Microbiome (AM) using their amplicon analysis workflow (Bissett *et al.* 2016; https://www.australianmicrobiome.com/protocols/16sanalysisworkflow/) and were downloaded as amplicon sequence variant (ASV) abundance tables from the AM portal (dates accessed, Huntly: 01 Feb 2021; Eneabba: 16 Nov 2020; Worsley: 16 Nov 2020) (27 Nov 2020) (https://www.australianmicrobiome.com/; AM sample IDs are listed in Table S1-S3). Note, in this study ASVs are equivalent to zero radius operational taxonomic units (zOTUs).

Briefly, paired end reads were merged using Flash2 (Magoč & Salzberg 2011), merged sequences were then further screened to remove those with ambiguities, long homopolymer runs, or too short/long using the *screen.seqs()* function from Mothur (Schloss *et al.* 2009). Reads passing filter were dereplicated and denoised to zOTUs using the UNOISE3 algorithm (Edgar 2016) in USEARCH (Edgar 2010). All reads were then mapped to zOTUs to construct a sample by zOTU (representative sequence) count table. zOTUs (which we refer to as ASVs) were assigned taxonomy in subsequent processing described below.

### Initial data cleaning

The AM workflow above produced 16S rRNA ASV-level abundance data for all minesites. The starting total number of sequences comprised: for Huntly, n = 1,791,794 sequences (36,061 ASVs); Eneabba, 2,271,627 sequences (50,845 ASVs); and Worsley, 2,097,206 sequences (64,026 ASVs).

Further data preparation was undertaken in R version 4.0.3 (R-Core-Team 2020) utilising the framework of the R *phyloseq* package (McMurdie & Holmes 2013) to manage the microbiome datasets.

Taxonomy was assigned across all ASVs using the *assignTaxonomy()* function from the R *dada2* package (Callahan *et al.* 2016), using the Silva version 138 prokaryotic taxonomic training database (silva_nr99_v138_train_set.fa.gz; https://zenodo.org/record/3986799#.X5kj-aozaUl; Aug 2020). This training database uses a 99% identity criterion to cluster highly identical sequences (https://www.arb-silva.de/documentation/release-138/) and where possible assigns taxonomy to the genus-level.

Initial data cleaning was undertaken to discard ASVs that were: not identified as belonging to bacteria, unidentified at the phylum level, assigned as chloroplast or mitochondria, or did not occur in at least two samples. Additionally, at Eneabba 7 sites were excluded with low sequence counts (≤ 5897 sequences; i.e., X138414, X138422, X138474, X138446, X138420, X138416, X138440). Total cleaned sequences and samples comprised: at Huntly, n = 1,772,249 sequences (31,746 ASVs) from 36 samples; Eneabba, 2,155,211 sequences (33,636 ASVs) from 26 samples; and Worsley, 2,049,625 sequences (54,671 ASVs) from 25 samples.

Phylogenetic trees were built to support Unweighted- and Weighted-UniFrac analyses via the following steps. Multiple sequence alignments were performed using MAFFT (v7.475; Katoh & Standley 2013) with the fast strategy (FFT-NS-2) suited for larger sequence alignments. Ambiguous and uninformative alignment regions were masked using Gblocks (v0.91b; Talavera & Castresana 2007) with options: *-t=d -b2=<60% of number of sequences> -b3=10 -b4=5 -b5=h*; refer to http://molevol.cmima.csic.es/castresana/Gblocks/Gblocks_documentation.html). For Huntly data the original MAFFT-alignment of 1755 base positions was masked down to a Gblocks-alignment of 336 positions; Eneabba data was masked from 1749 to 372 positions; and Worsley data was masked from 2610 to 346 positions. We note that inclusion of a masking step is consistent with mainstream workflows such as in QIIME2 (Bolyen *et al.* 2019; https://docs.qiime2.org/2021.4/tutorials/phylogeny/#reducing-alignment-ambiguity-masking-and-reference-alignments). Then maximum likelihood phylogenetic trees were inferred using IQTREE2 (v2.1.2, Oct 2020; Minh *et al.* 2020) using a GTR + I +G model (i.e., general time-reversible model with rate heterogeneity allowing for a proportion of invariable sites plus discrete Gamma model; http://www.iqtree.org/doc/iqtree-doc.pdf) with a maximum of 200 search iterations.

### Potential limitations in our study with the use of phylogenetic trees

We performed the de novo alignment using MAFFT and Gblocks after earlier attempting to build IQTREE2 trees based on SILVA Incremental Aligner (SINA; Pruesse, Peplies & Glöckner 2012) derived alignments, which used SILVA-curated trees from the SILVA SSU Ref NR 99 database (SILVA_138.1_SSURef_NR99_12_06_20_opt.arb.gz from https://www.arb-silva.de/download/arb-files/) followed by gap-filtering via the Mothur *filter.seqs()* function. However, this attempted tree-building using gap-filtered SINA alignments failed to find converging results with our soil bacterial datasets in a timely manner with our available high performance computing resources.

Ultimately we used de novo phylogenetic trees to generate UniFrac distances and were not able to utilise existing SILVA-curated trees, due to computational challenges. Tree-building often represents a compromise between accuracy in representing phylogenetic relationships and computing time, and it was beyond the scope of our study to test the sensitivity of our UniFrac-based analyses to the quality of trees used.

### OTU clustering

Clustering of operational taxonomic units (OTUs) at a number of %-identity thresholds was achieved by first ordering the ASV-level data from highest abundance to lowest abundance, then we applied the VSEARCH (Rognes *et al.* 2016; https://github.com/torognes/vsearch) *cluster_smallmem* algorithm to produce respective uclust-like (Edgar 2010) format outputs for each minesite which were then analysed in R to generate the clustered phyloseq data objects.

### Exploratory data analysis

*Alpha diversity and evenness*
During exploratory data analysis, we visualised the alpha diversity of rarefied samples, for all minesites, expressed as the effective number of ASVs, or exp( Shannon's Index) (Jost 2006), and also as composite samples within rehabilitation-age groups using merged-sample bootstrap resampling from the unrarefied data (Liddicoat *et al.* 2019) (SI Appendix, Figure S3). Similarly, the evenness of ASVs across rehabilitation-age groups and minesites was calculated using Pielou's evenness index (J' = Shannon's diversity index (H') divided by the natural logarithm of species richness; Pielou 1966) based on rarefied data, and using merged-sample bootstrap resampling from unrarefied data (SI Appendix, Figure S4).

*Relative abundance heatmaps*
The relative abundance of phyla, classes, and orders that were present at each minesite (using unrarefied data) were indicated in heatmaps, created using the *plot_heatmap()* function in the R *phyloseq* package SI Appendix, Figures S5–S13.

*Associations with key soil variables*
We used constrained correspondence analysis (CCA) via the *ordiR2step()* function from the R *vegan* package on scaled (i.e., mean-centred and divided by standard deviation) data for available soil variables, latitude and longitude, to visualise the sample-wise variation in bacterial communities that was associated with variation in soil and landscape properties for each minesite. The *ordiR2step* function performs automatic stepwise model-building using permutation tests. We excluded highly correlated soil variables prior to the CCA analysis after identifying them using the *findCorrelation()* function from the R *caret* package (Kuhn 2020) with a cut-off Pearson's r value of 0.75. Correlated soil variables were visualised using correlation plots. The significance of terms identified by the

CCA was tested using permuted ANOVAs on the reduced *ordiR2step* model with 999 permutations and testing each term separately. Variables identified from the stepwise modelling and CCA analysis were also plotted to visualise their variation across rehabilitation ages. Correlation plots and CCA plots are presented in SI Appendix, Figures S14–S19; and results are described in SI Appendix Supplementary Data. Biologically-associated soil variables identified through the CCA were then plotted to show variation across rehabilitation ages (SI Appendix, Figure S20). This analysis should be considered preliminary, as it involved an automated step to eliminate correlated variables. Future more detailed analyses should consider wider minesite-specific information and expert knowledge to optimise variable selection in the CCA analyses.

**Rationale and approach for preliminary exploration of spatial autocorrelation**

To examine potential implications of spatial autocorrelation on our rehabilitation trajectory analyses, we visualised the relationship between ecological distance and the geographic separation of samples, for rehabilitation age and reference groups. Similar to the rehabilitation trajectory plots, we restricted analyses to the set of data that involved distances to references. We lacked the quantity of data to produce variograms, which are a common tool in spatial statistics to model spatial autocorrelation. However, we did produce variogram-like plots with ecological distance (between samples and references) on the y-axis, and geographic distances (between samples and references) on the x-axis. Because spatial autocorrelation is a natural phenomenon, we assumed the curve representing reference samples (i.e. among-reference distances) offered a natural baseline curve for spatial autocorrelation in that minesite environment. Each rehabilitation age group was modelled as a second-order polynomial, allowing the possibility of curvilinear trendlines which could permit (although did not constrain) the data to be expressed in variogram-like relationship (i.e., increasing then flattening). We expected the curves for various rehabilitation ages to express different mean y-values (i.e., increased ecological distance at younger ages) but considered differences in the shape of the curves to represent a departure from the assumed natural pattern of spatial autocorrelation for that minesite. Then for each rehabilitation age group we calculated the difference in mean-centred model curves (rehabilitation age group minus reference) and applied a correction to the data such that each rehabilitation age group expressed the same ecological distance-geographic distance curvilinear trend as seen for references. Restoration trajectories and predicted recovery times were then compared between 'original' and 'corrected' data, using the corresponding Bray-Curtis similarities. As discussed in the main paper, the presence of 'direct return' soils in young rehabilitation sites, which display elevated similarity to reference, represents a limitation in our first-pass analysis, because we do not account for the previous spatial location of 'direct return' soils. However, we also recommend to exclude such samples (i.e., young 'direct return' soils with elevated similarity to reference) from the trajectory modelling. With older rehabilitation sites, we suggest that localised factors will become dominant in shaping microbiota within a few years, and thereby overcome the influences associated with the previous spatial location of 'direct return' soils.

### Supplementary Data

**Biologically-associated soil and landscape variables**

[Refer to SI Appendix Figures S14–20]

At Huntly, sample bacterial communities associated with conductivity (dS/m) (ANOVA df = 1, $\chi^2$ = 0.3540, F = 2.6589, P = 0.001), Colwell phosphorus (mg/kg) (df = 1, $\chi^2$ = 0.2668, F = 2.0038, P = 0.001), exchangeable aluminium (meq/100g) (df = 1, $\chi^2$ = 0.2543, F = 1.9102, P = 0.002), and copper (DTPA extraction; mg/kg) (df = 1, $\chi^2$ = 0.2181, F = 1.6387, P = 0.014). Conductivity generally increased with rehabilitation age although levels are considered non-saline. The youngest samples contained high phosphorus levels. Elevated exchangeable aluminium levels (correlated with lower pH) were found in 17-yr and 25-yr rehabilitation age samples. Samples from the 14-yr, 17-yr and 25-yr rehabilitation ages contained higher levels of copper.

At Eneabba, sample bacterial communities associated with organic carbon mass-concentration (%) (df = 1, $\chi^2$ = 0.3637, F = 1.3695, P = 0.001), and clay content (%) (df = 1, $\chi^2$ = 0.3212, F = 1.2092, P = 0.020). Reference samples had higher organic carbon content, and younger sites were generally sandier.

At Worsley, soil bacterial communities associated with exchangeable aluminium (meq/100g) (df = 1, $\chi^2$ = 0.3364, F = 2.0597, P = 0.001), iron (DTPA extraction; mg/kg) (df = 1, $\chi^2$ = 0.2342, F = 1.4336, P = 0.004), and latitude (decimal degrees) (df = 1, $\chi^2$ = 0.2189, F = 1.3399, P = 0.016). Young rehabilitation samples generally had low exchangeable aluminium levels and low iron levels, and these elements increased in older rehabilitation and reference samples.

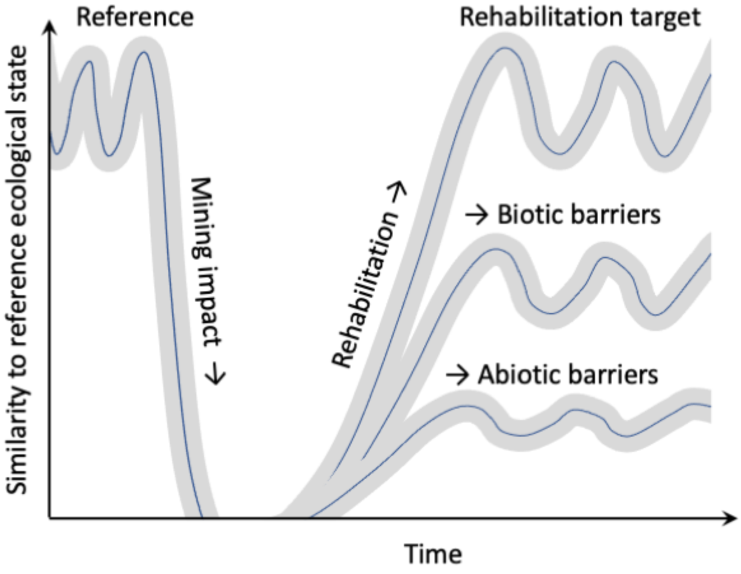


FIGURE S1. Hypothetical recovery towards reference ecological state with post-mining rehabilitation. Note there is natural variation due to dynamic (e.g. seasonal) conditions and also variation among references. Abiotic (e.g. limiting soil conditions) and biotic barriers (e.g. failure to re-establish key ecosystem components) can impede full recovery. Adapted from Manero, Standish and Young (2021) and Australian_Government (2016).

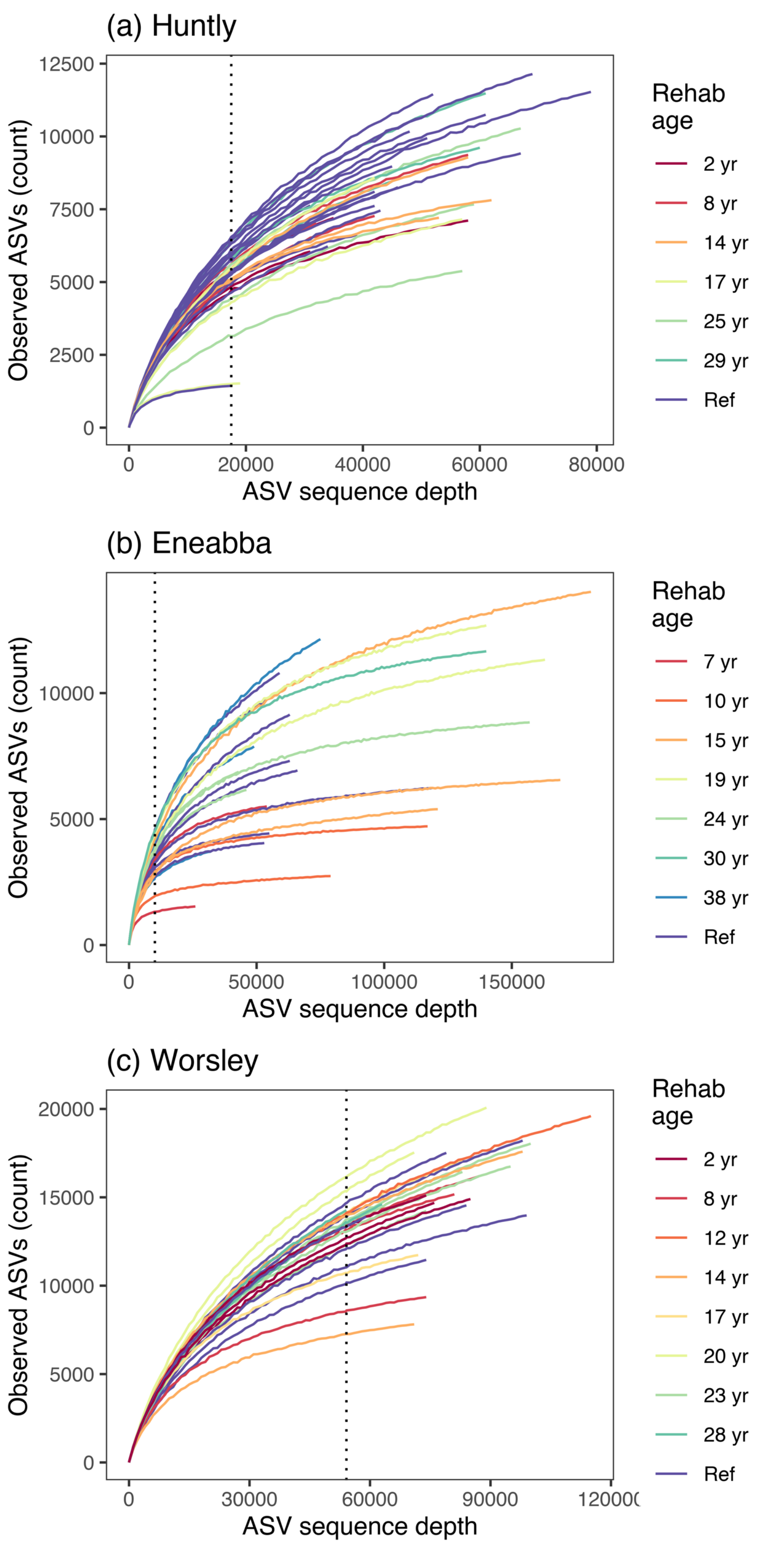


FIGURE S2. Rarefaction curves for (a) Huntly, (b) Eneabba, and (c) Worsley surface soil bacterial 16S amplicon datasets. Minimum sample sequence read depths are denoted in the dotted vertical lines (i.e., Huntly: 17,485 sequences; Eneabba: 10,142 sequences; Worsley: 54,122 sequences).

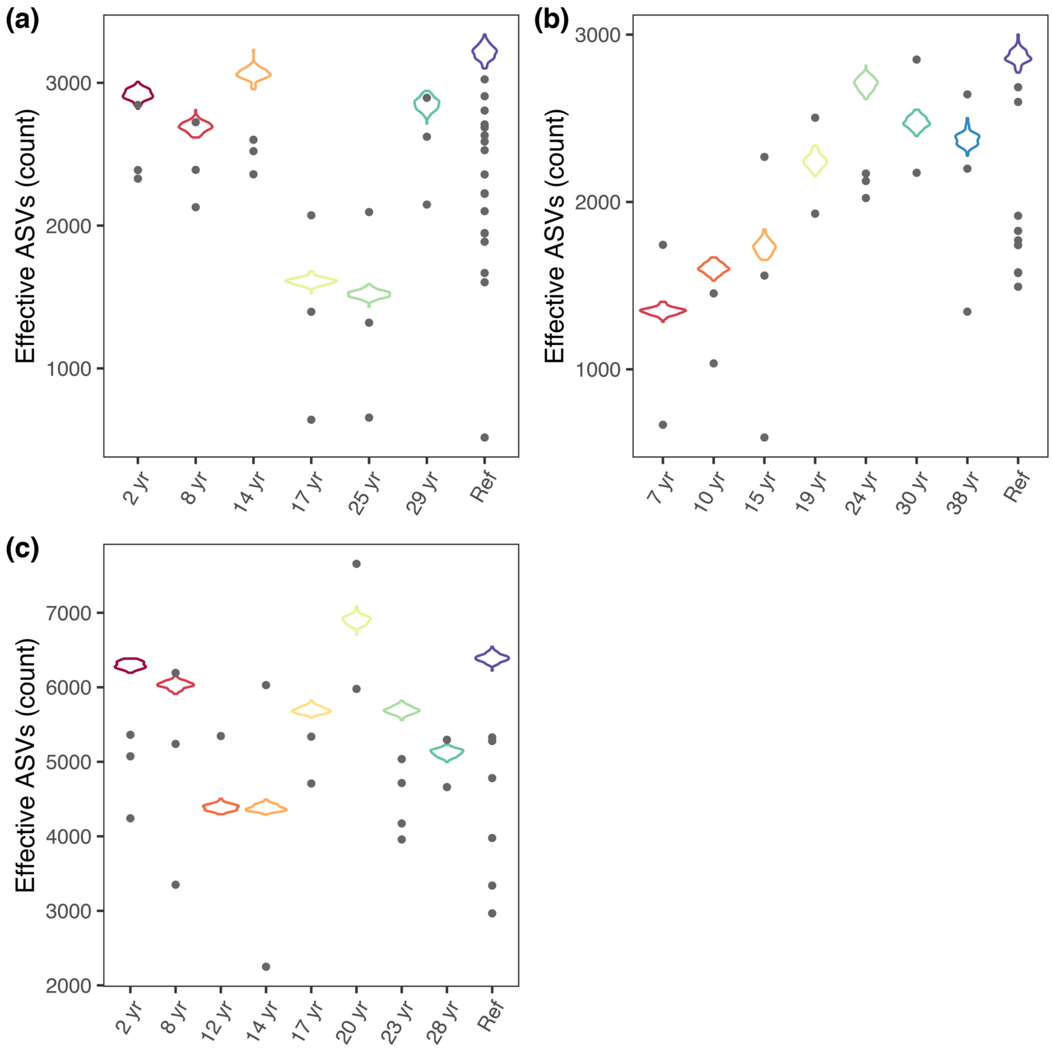


FIGURE S3. Effective number of ASVs (alpha diversity) based on exp(Shannon's index) of samples from (a) Huntly, (b) Eneabba, and (c) Worsley. Violin-plot distributions represent the overall alpha diversity across samples within a rehabilitation age group calculated using merged-sample bootstrap resampling (B = 100). Grey dots represent alpha diversity within each sample calculated from one-off rarefying to the minimum sample sequence read depth (i.e., Huntly: 17,485 sequences; Eneabba: 10,142 sequences; Worsley: 54,122 sequences).

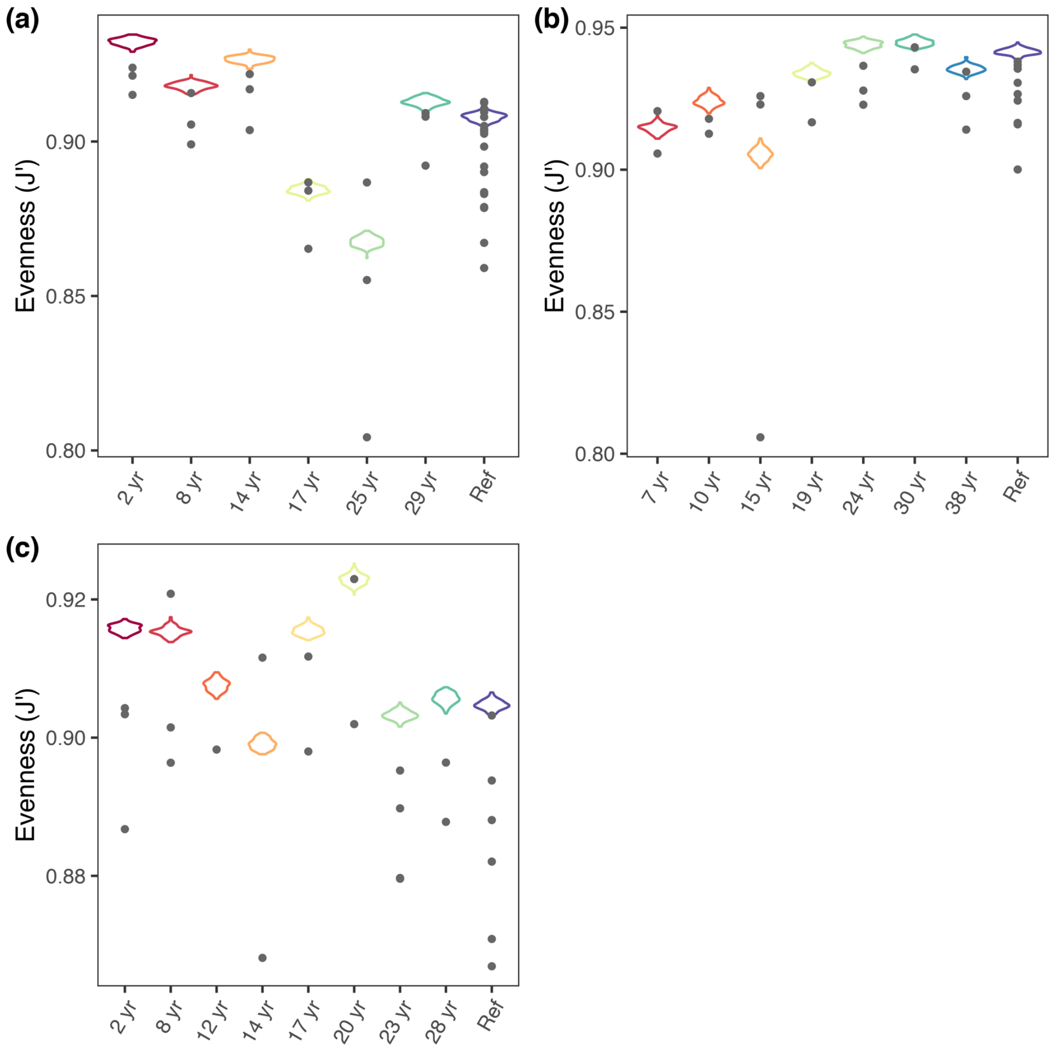


FIGURE S4. Evenness of ASVs based on Pielou's evenness index (J') of samples from (a) Huntly, (b) Eneabba, and (c) Worsley. Violin-plot distributions represent the overall evenness across samples within a rehabilitation age group calculated using merged-sample bootstrap resampling (B = 100). Grey dots represent evenness within each sample calculated from one-off rarefying to the minimum sample sequence read depth (i.e., Huntly: 17,485 sequences; Eneabba: 10,142 sequences; Worsley: 54,122 sequences).

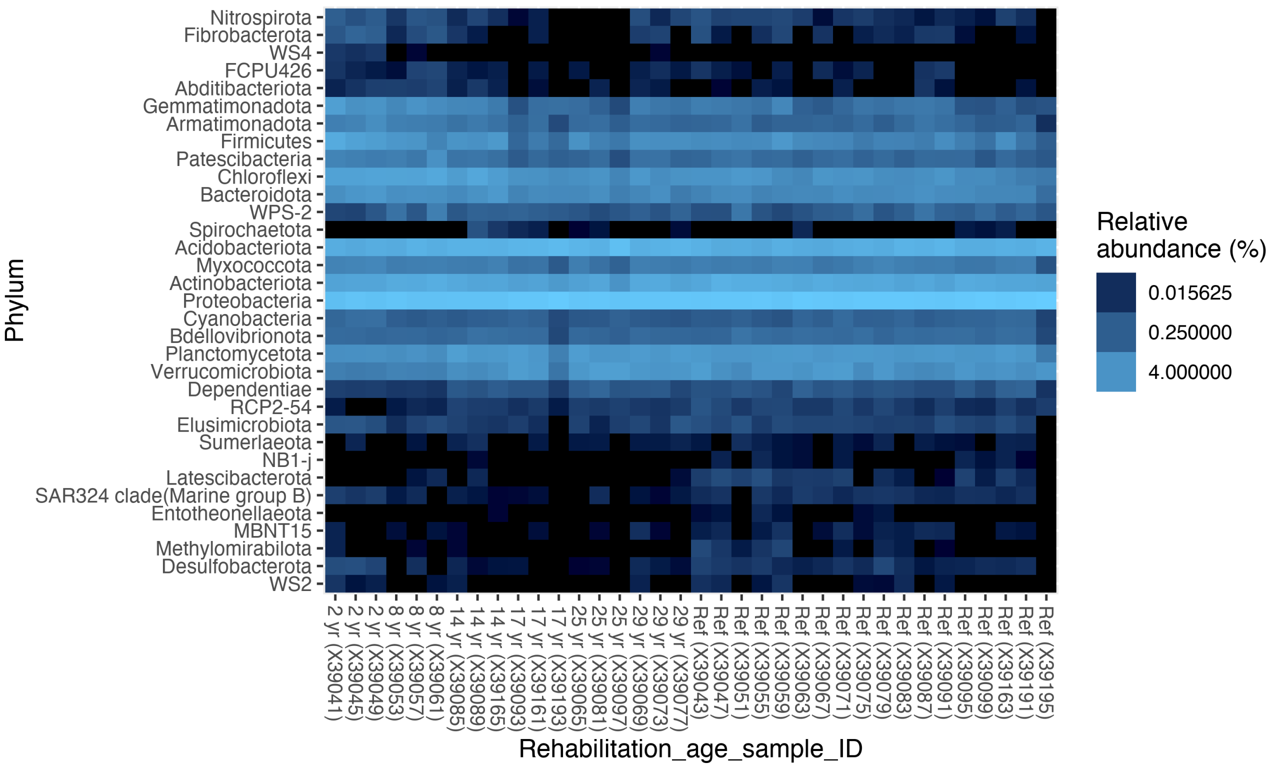


FIGURE S5. Huntly phylum-level (n = 33) heatmap of sequence relative abundance (%).

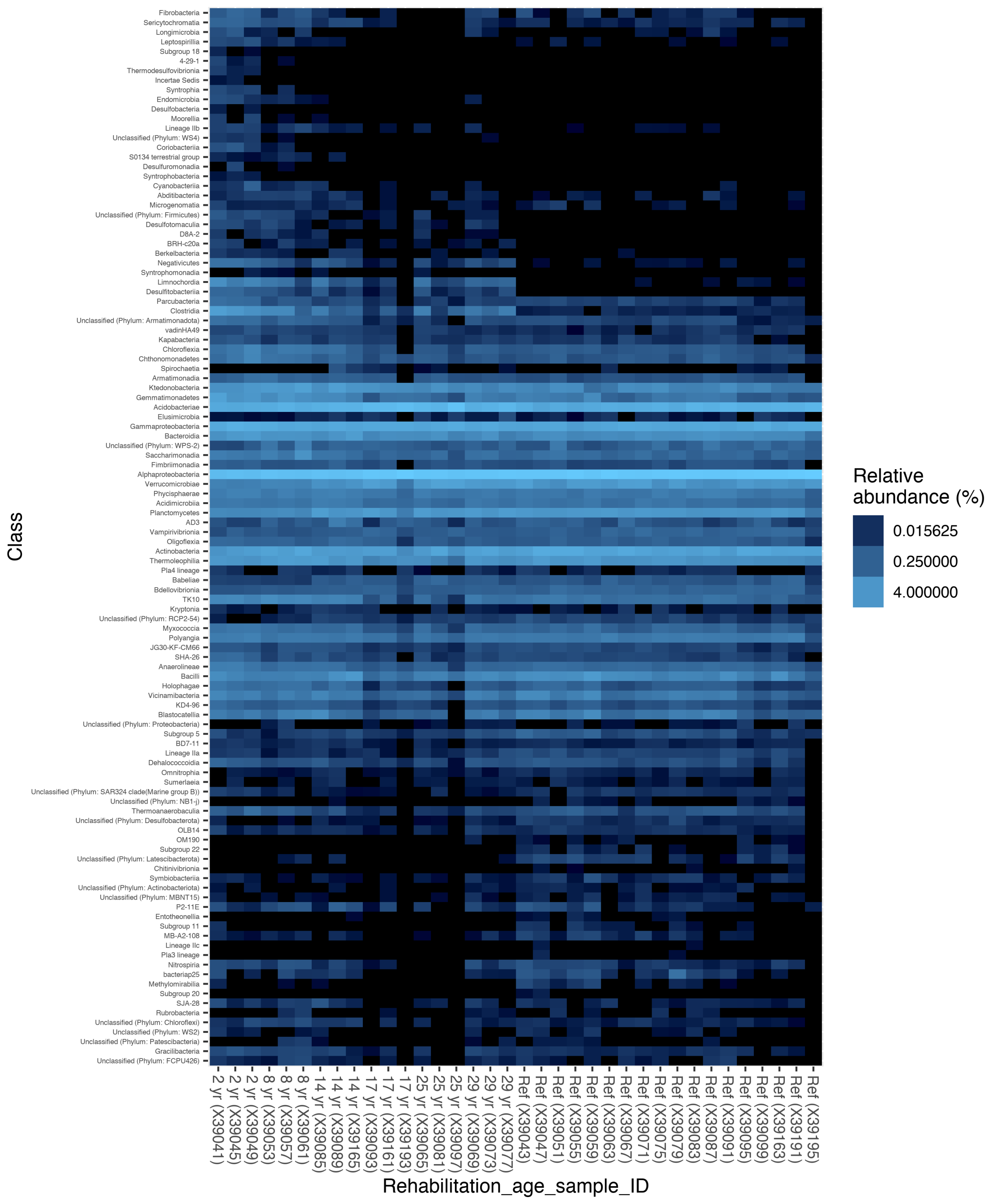


FIGURE S6. Huntly class-level heatmap of sequence relative abundance (%). From a total of 110 class-level groups, 16 are not assigned to a class but are labelled at the next available taxonomic level.

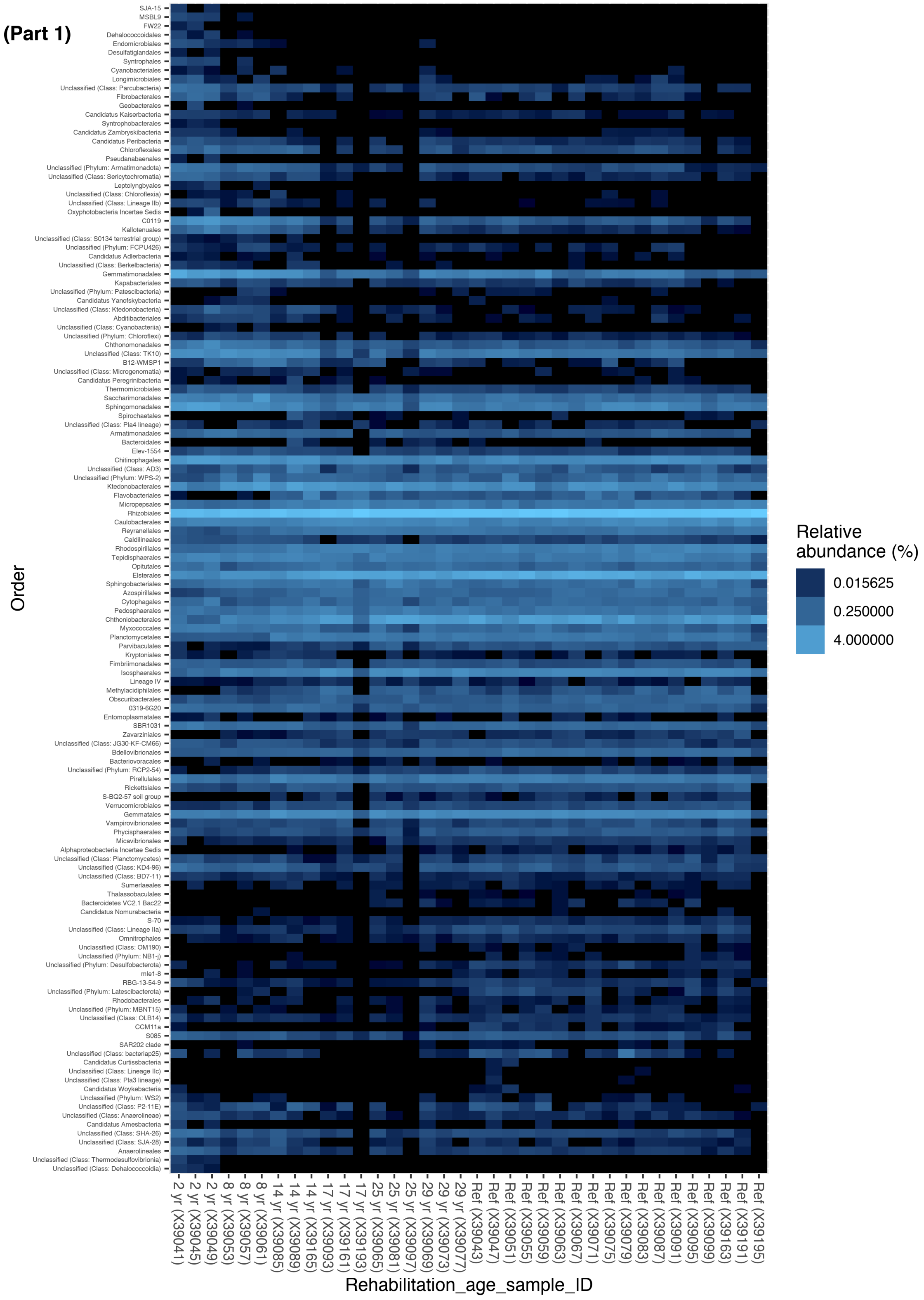


FIGURE S7. Huntly order-level heatmap (Part 1) of sequence relative abundance (%). 266 order-level groups are identified, with taxa 1-132 in this panel, 133-266 in the Part 2 (next page). 64 taxonomic groups are not assigned to an order but are labelled at the next available taxonomic level.

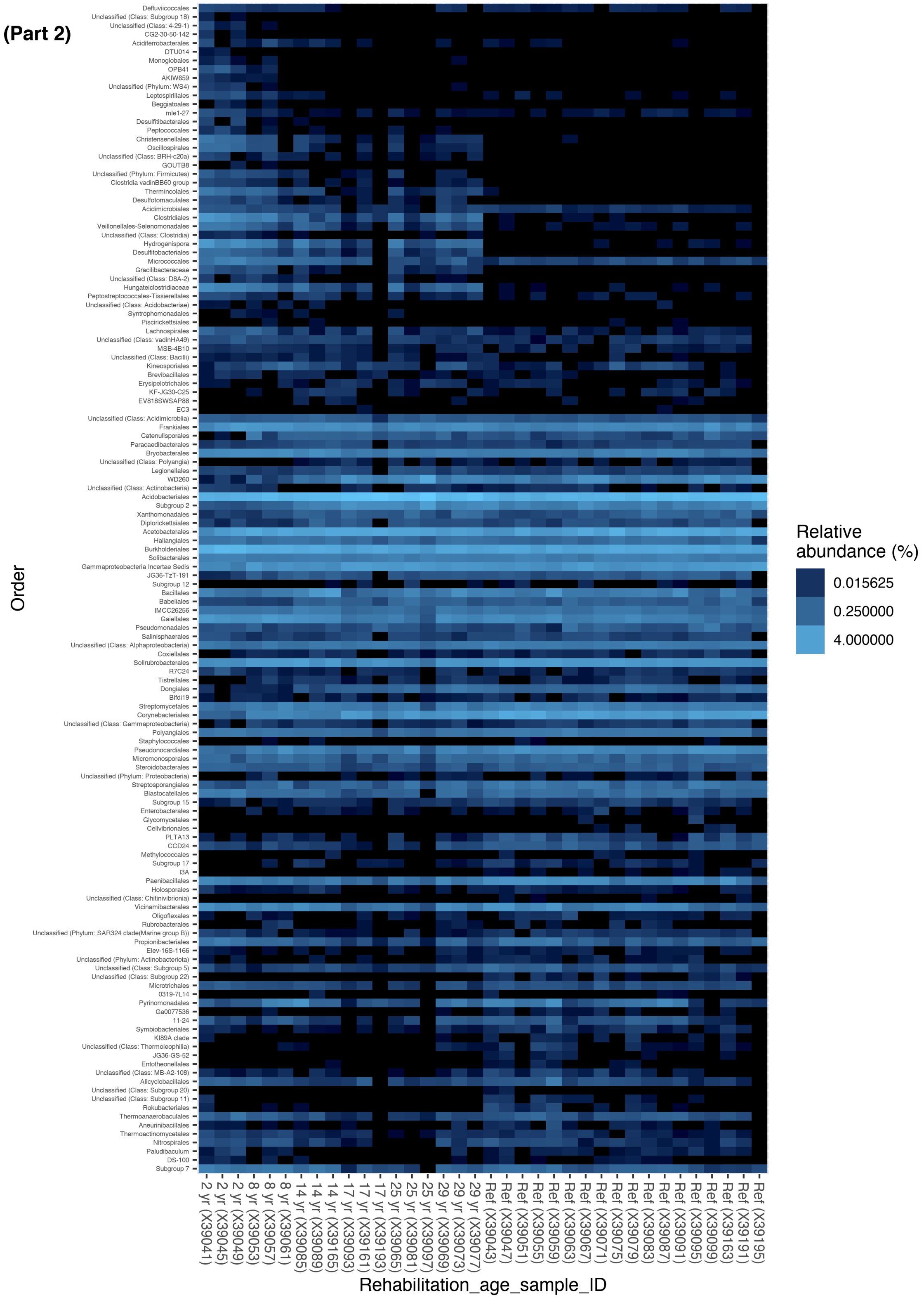


FIGURE S7 (Continued). Huntly order-level heatmap (Part 2) of sequence relative abundance (%). Refer to description in Part 1 (previous page).

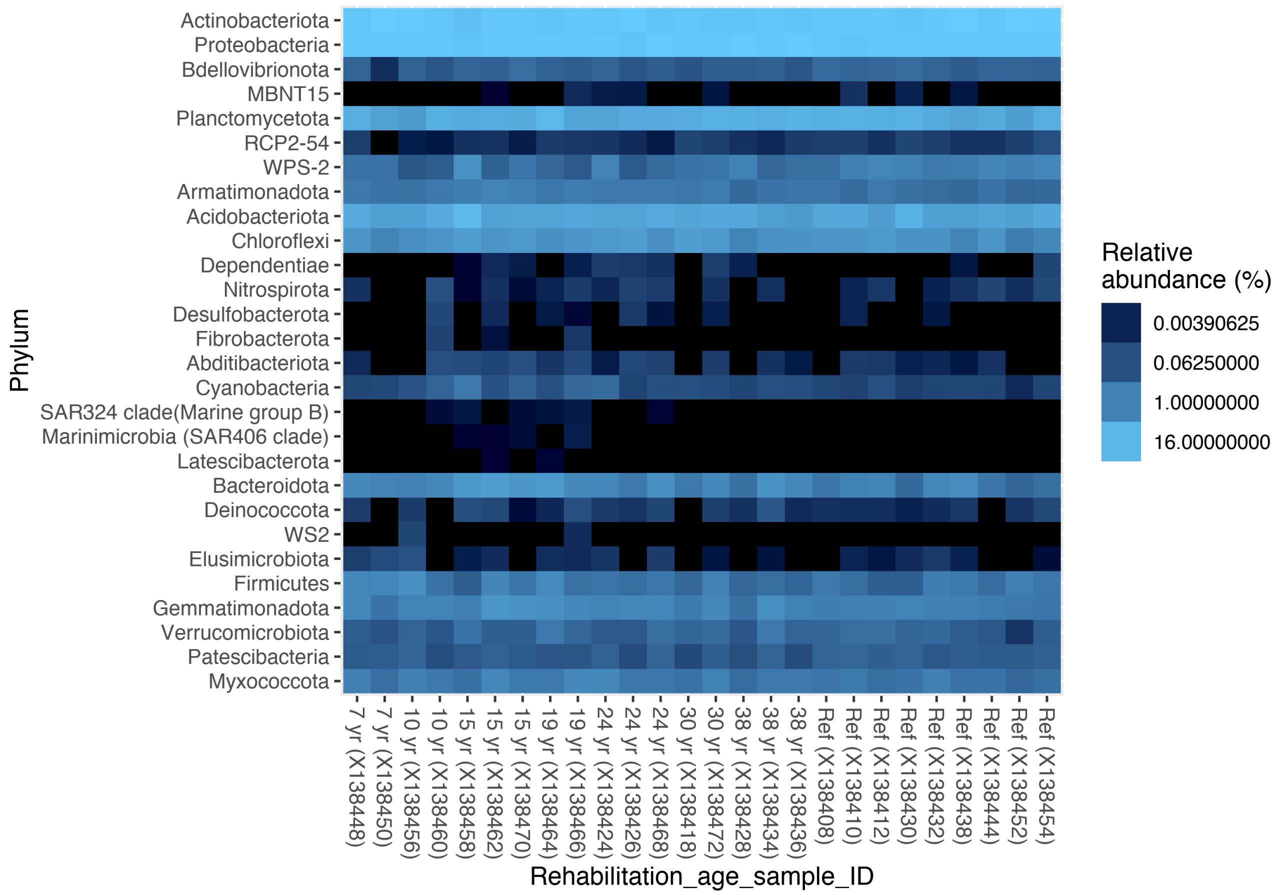


FIGURE S8. Eneabba phylum-level (n = 28) heatmap of sequence relative abundance (%).

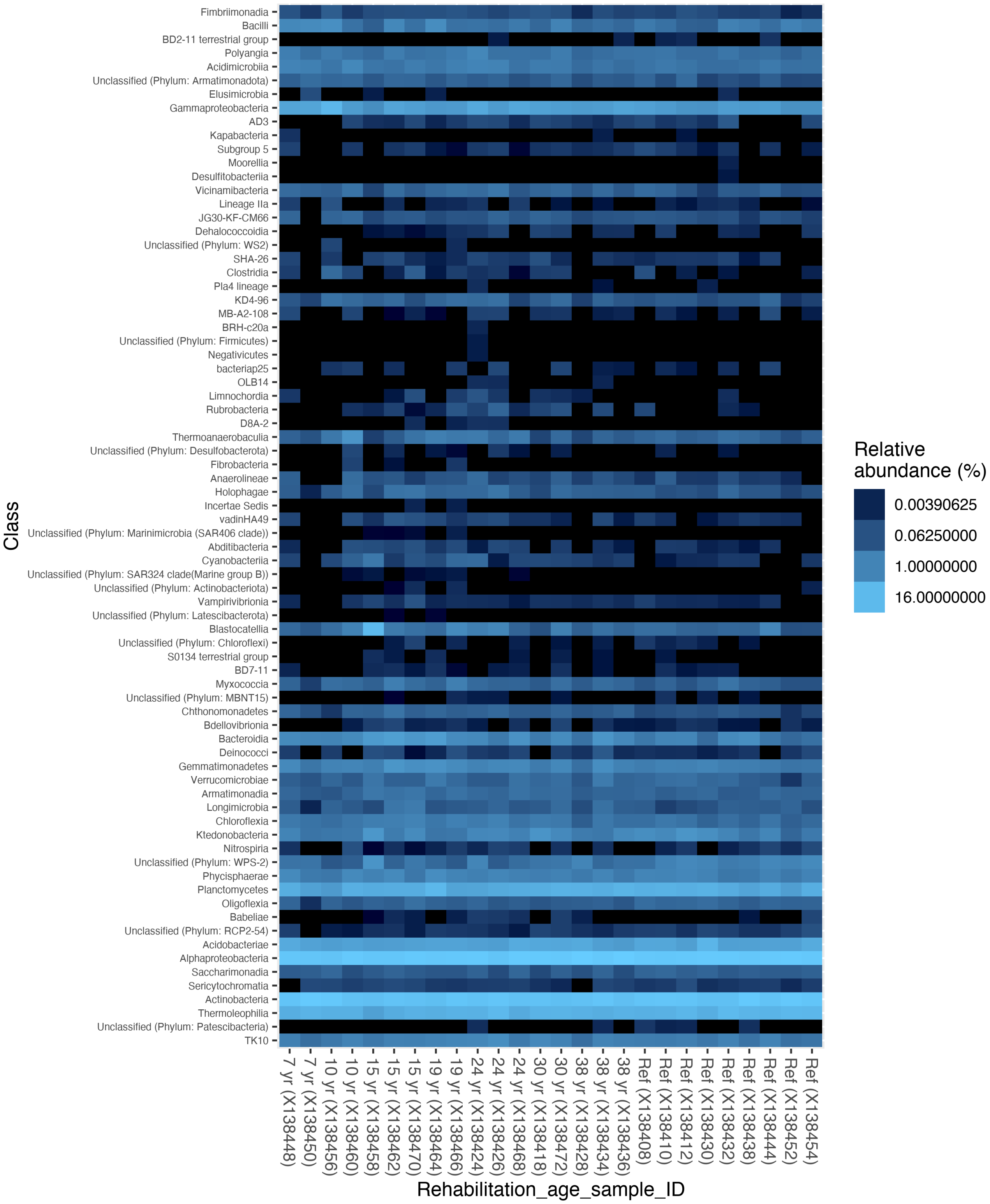


FIGURE S9. Eneabba class-level heatmap of sequence relative abundance (%). From a total of 76 class-level groups, 13 are not assigned to a class but are labelled at the next available taxonomic level.

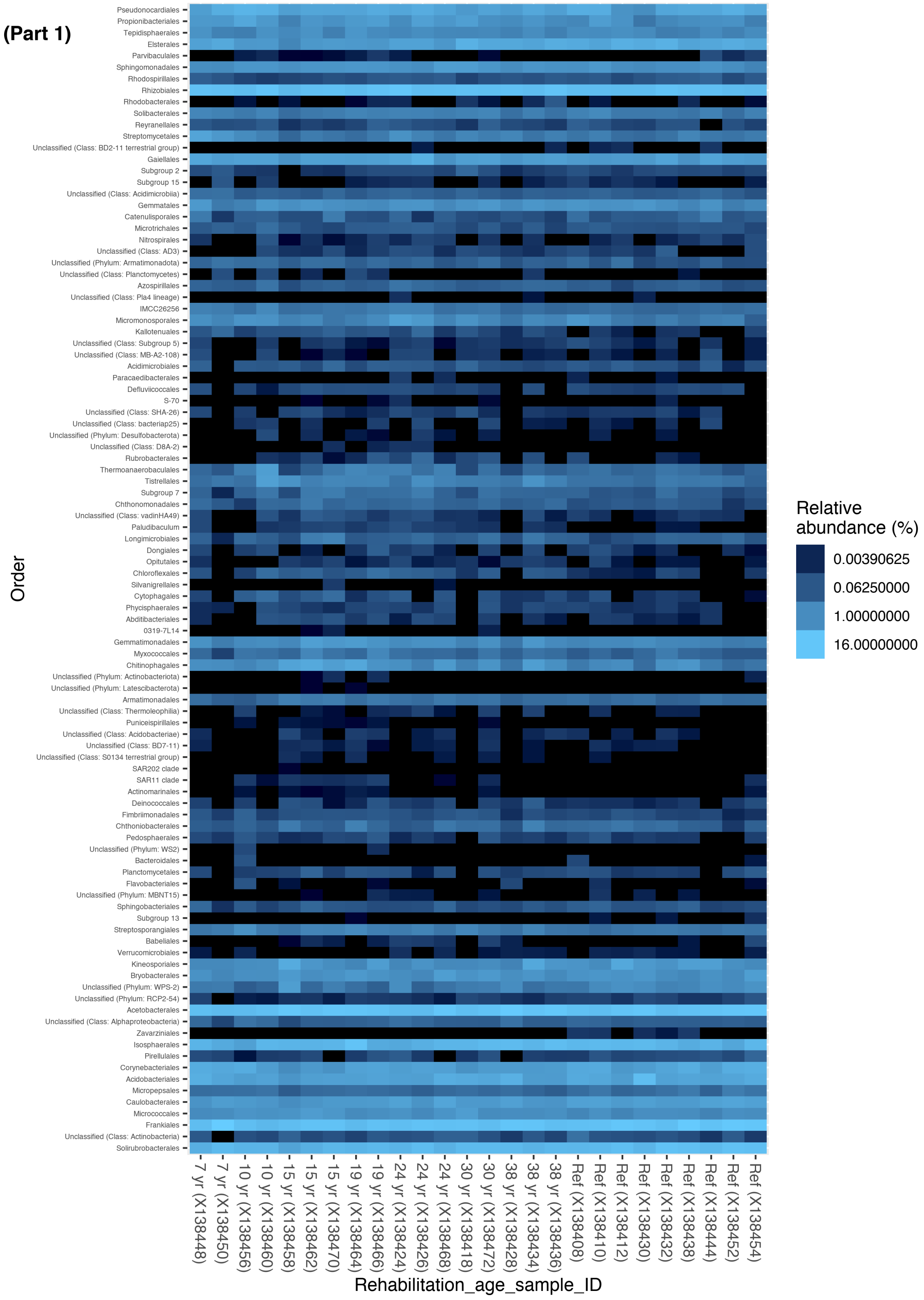


FIGURE S10. Eneabba order-level heatmap (Part 1) of sequence relative abundance (%). 194 order-level groups are identified, with taxa 1-100 in this panel, 101-194 in the Part 2 (next page). 43 taxonomic groups are not assigned to an order but are labelled at the next available taxonomic level.

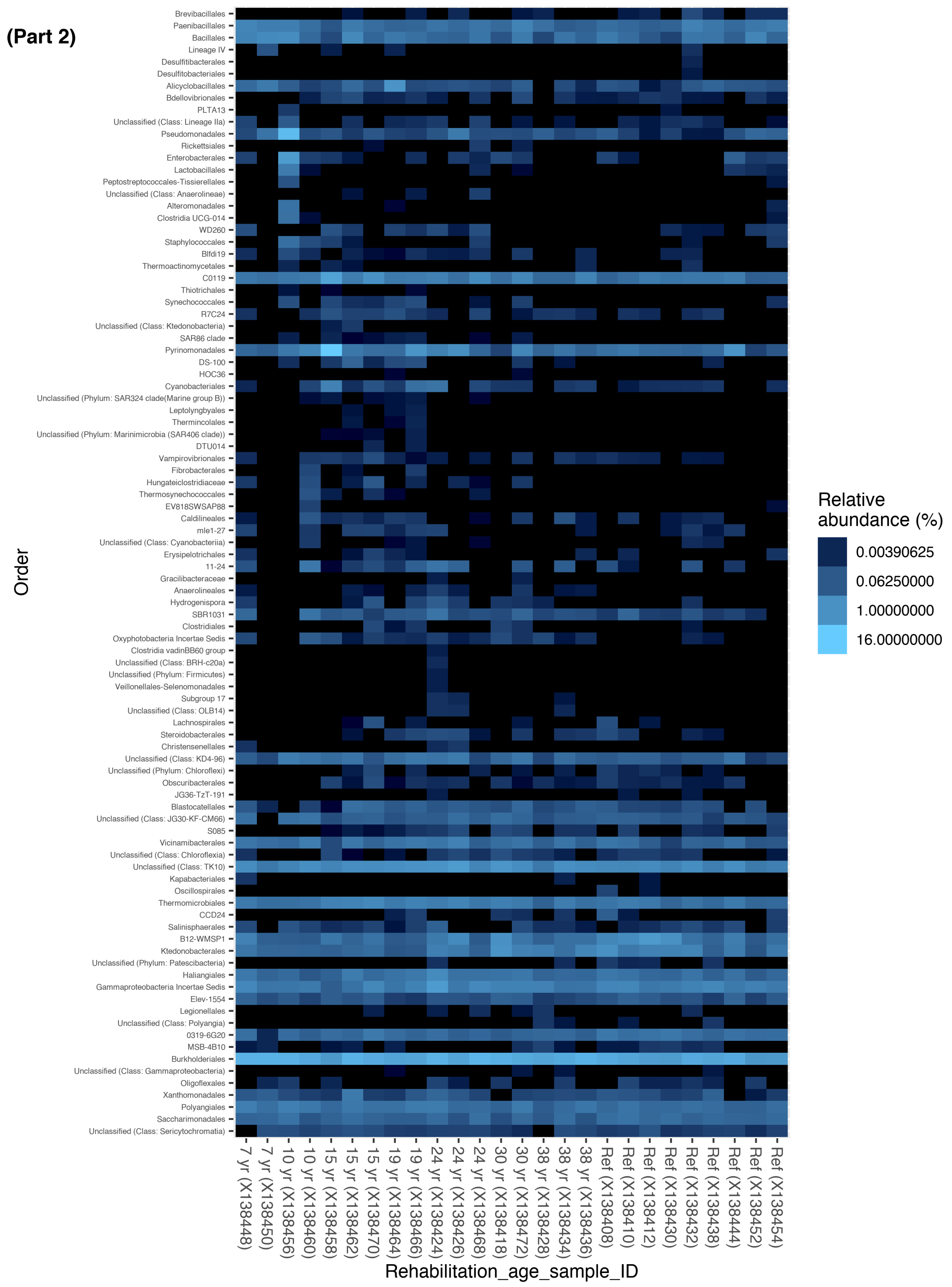


FIGURE S10 (Continued). Eneabba order-level heatmap (Part 2) of sequence relative abundance (%). Refer to description in Part 1 (previous page).

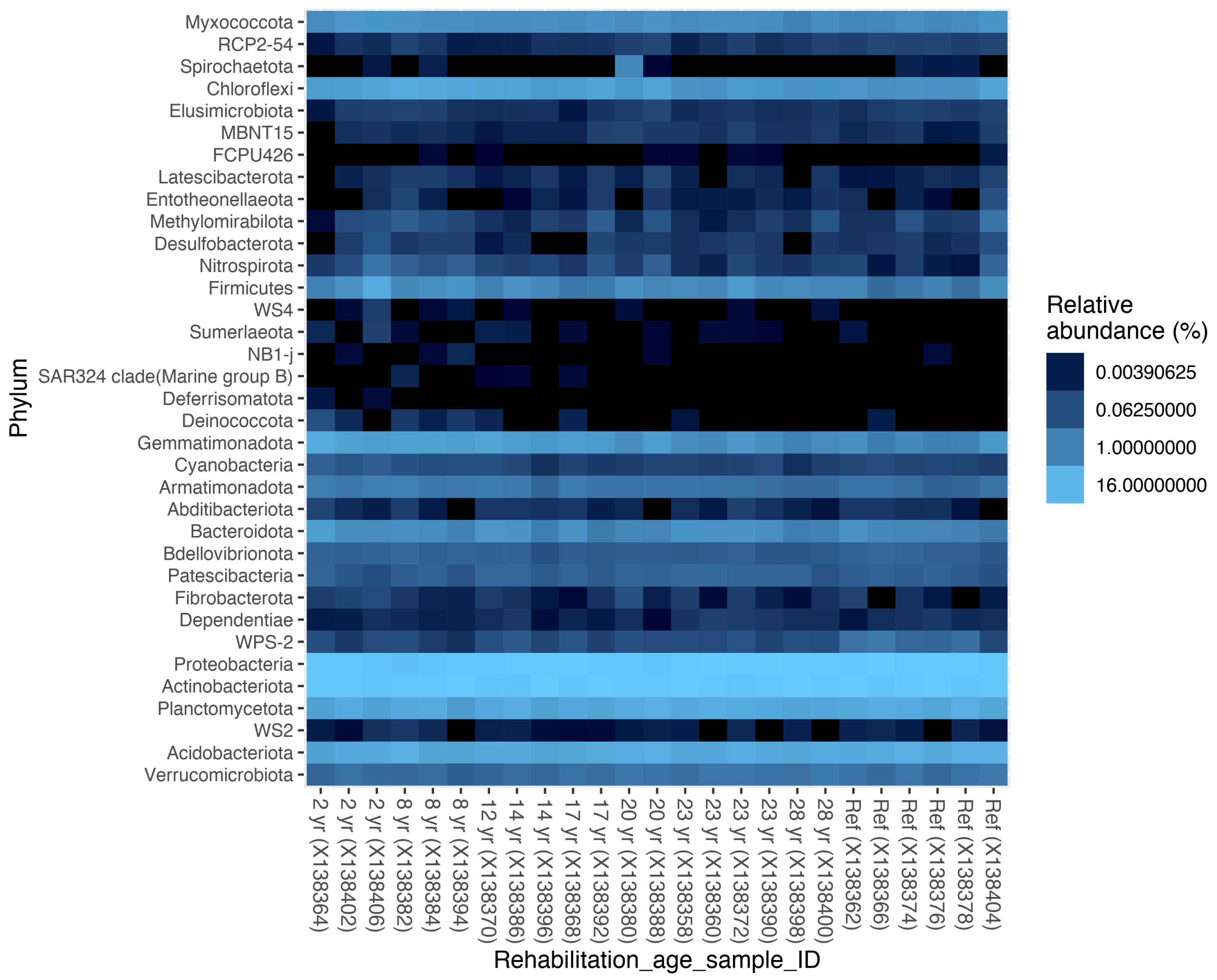


FIGURE S11. Worsley phylum-level (n = 35) heatmap of sequence relative abundance (%).

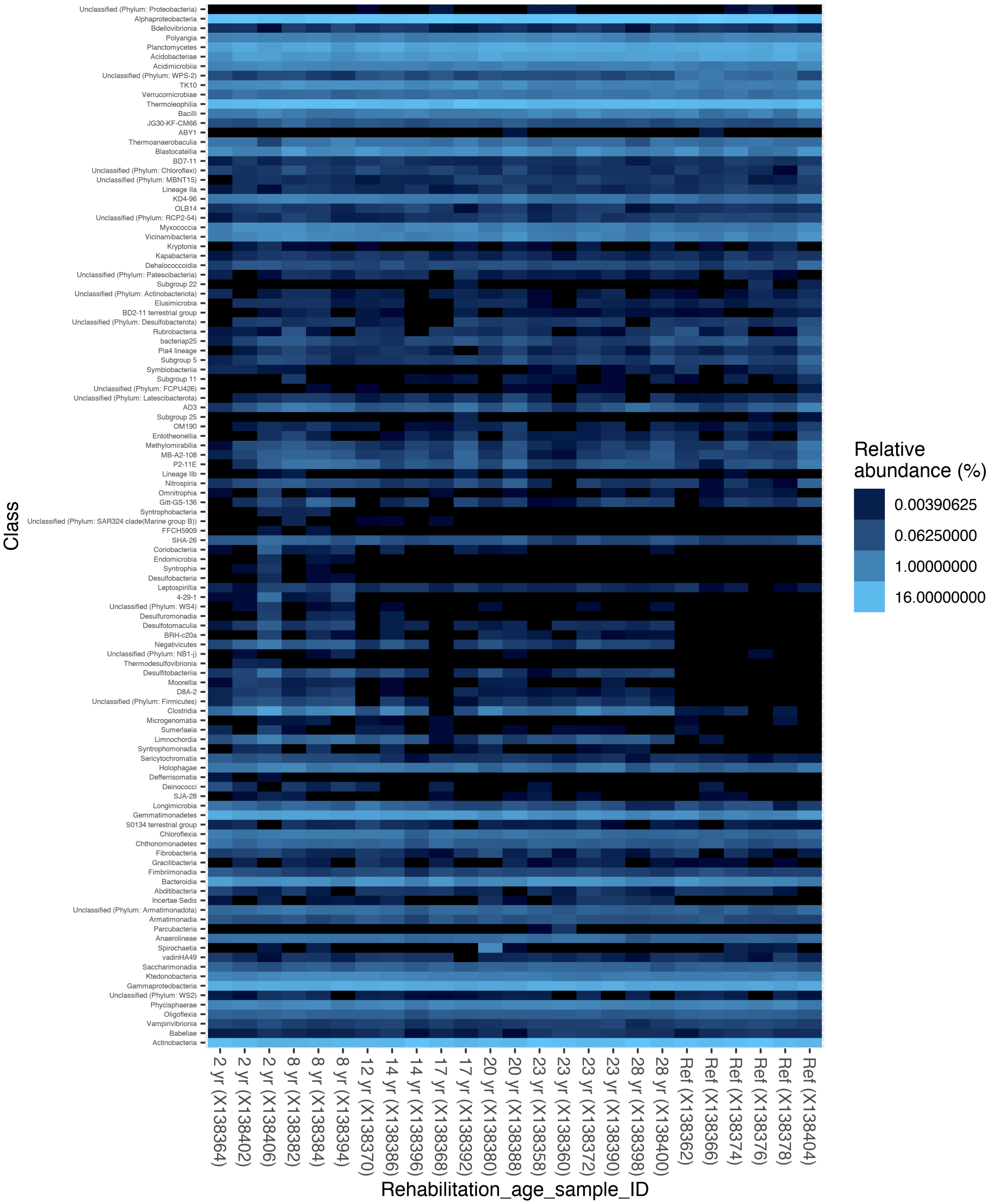


FIGURE S12. Worsley class-level heatmap of sequence relative abundance (%). From a total of 110 class-level groups, 16 are not assigned to a class but are labelled at the next available taxonomic level.

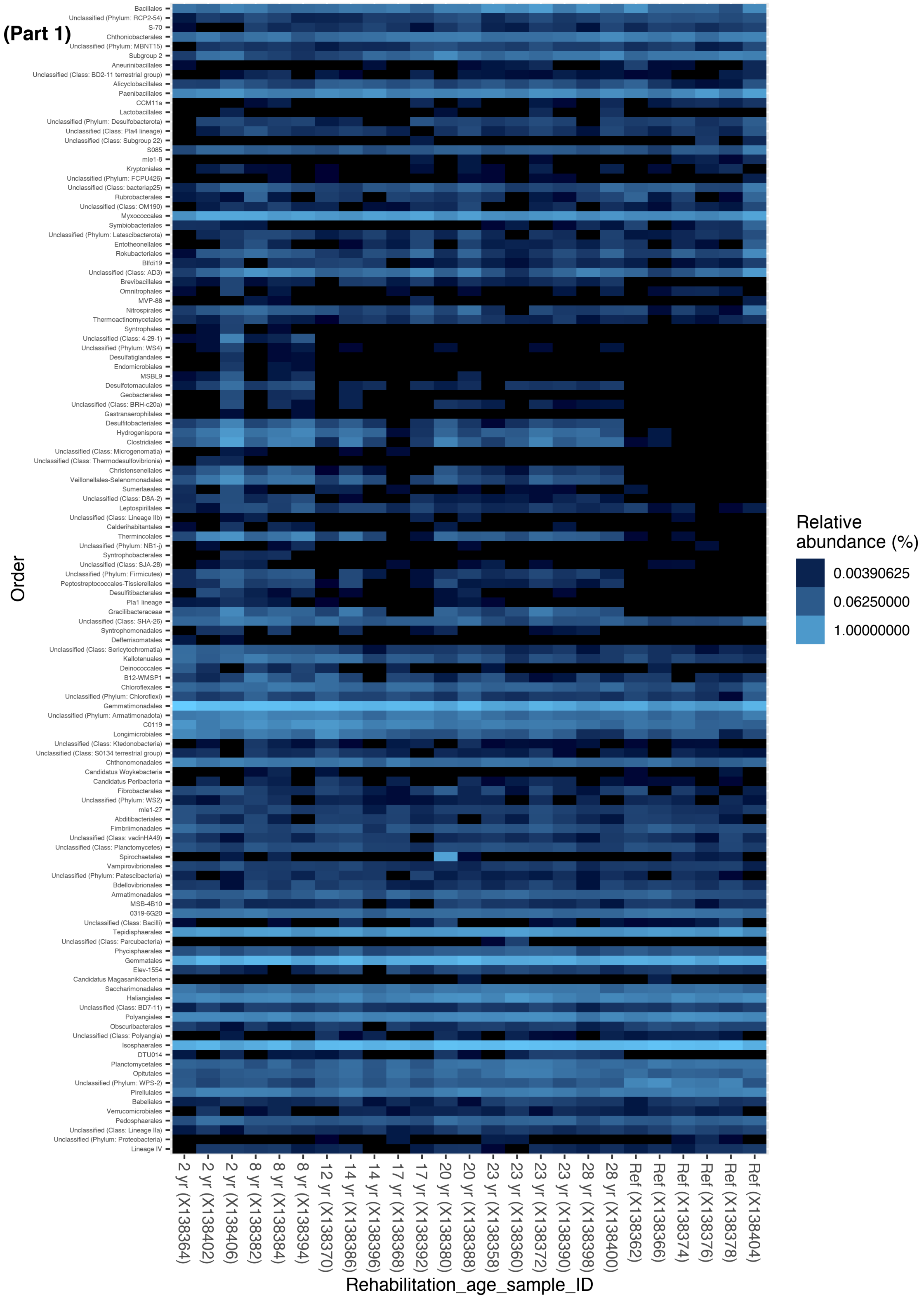


FIGURE S13. Heatmap (Part 1) of sequence relative abundance (%) grouped at the order-level for Worsley. 244 order-level groups are identified, with taxa 1-100 in this panel, 101-194 in the Part 2 (next page). 59 taxonomic groups are not assigned to an order but are labelled at the next available taxonomic level.

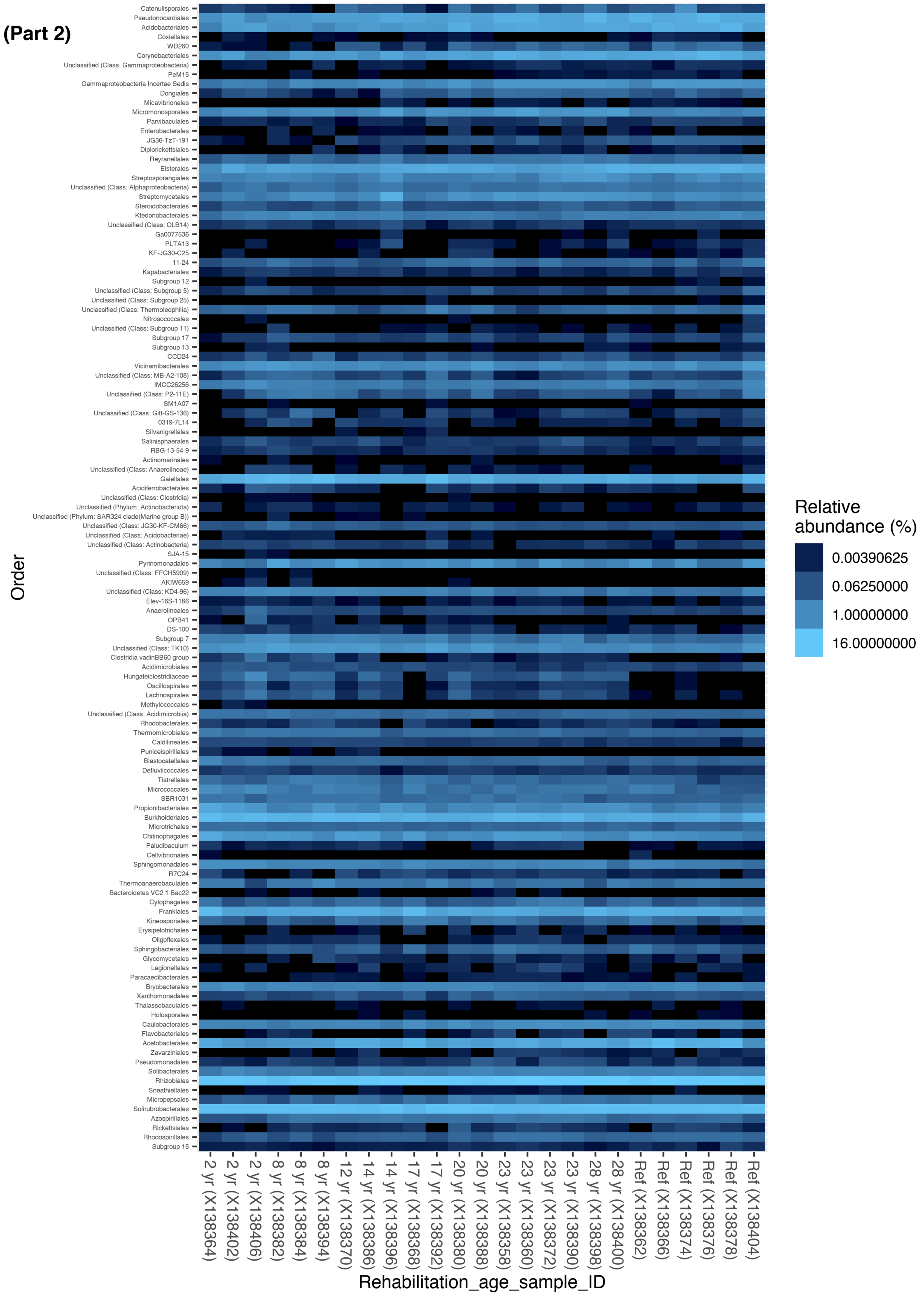


FIGURE S13 (Continued). Heatmap (Part 2) of sequence relative abundance (%) grouped at the order-level for Worsley. Refer to description in Part 1 (previous page).

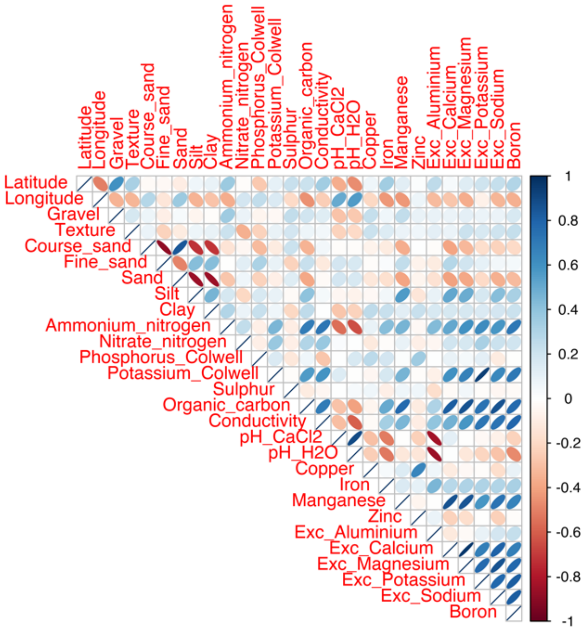


FIGURE S14. Correlation plot for soil sample/site variables at Huntly.

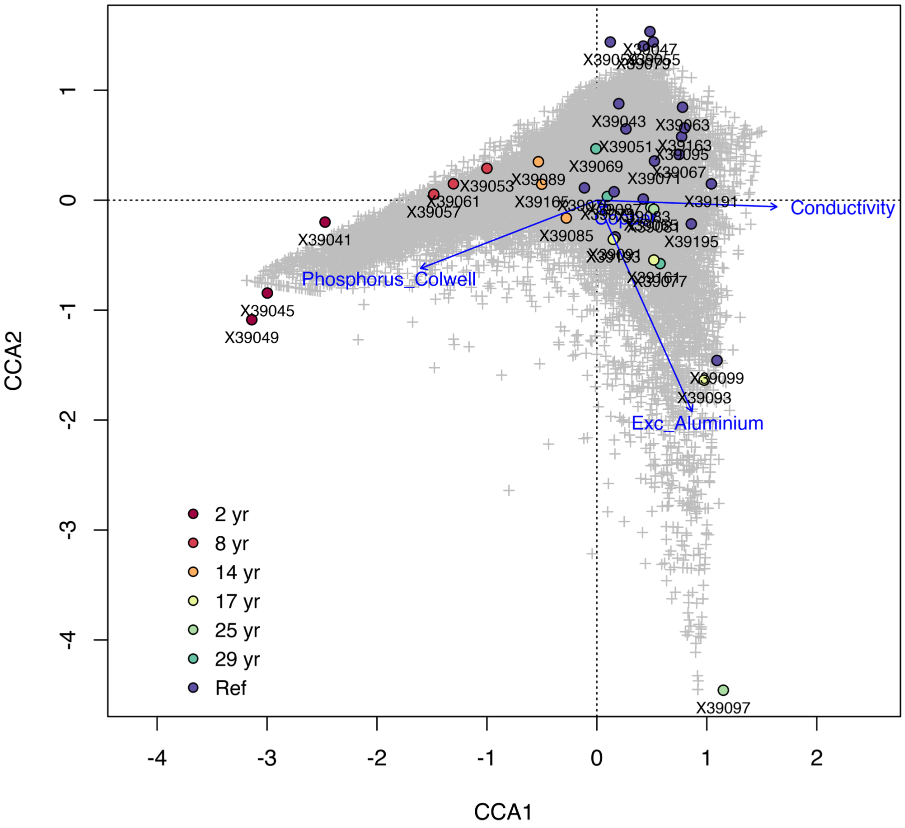


FIGURE S15. Constrained correspondence analysis (CCA) plot for Huntly displays variation in the bacterial community structure associated with scaled soil variables: Colwell-phosphorus (mg/kg), exchangeable aluminium (meq/100g), conductivity (dS/m), and copper (DTPA extraction; mg/kg). High exchangeable aluminium correlates with low pH. Grey points (+) denote taxa associated with variation in the soil variables.

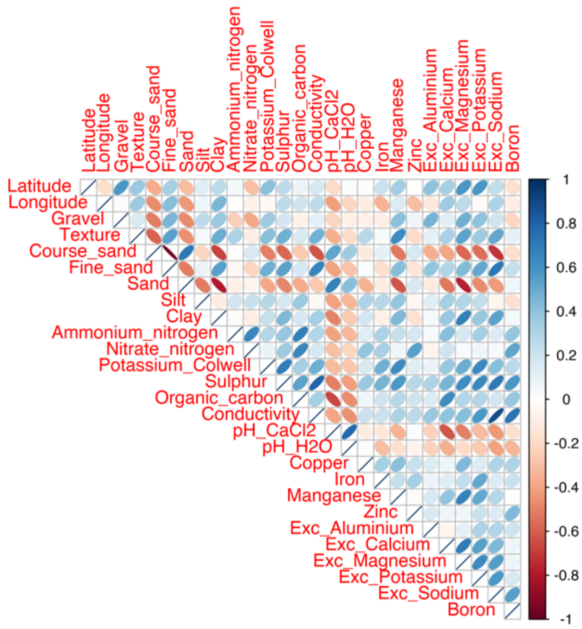


FIGURE S16. Correlation plot for soil sample/site variables at Eneabba.

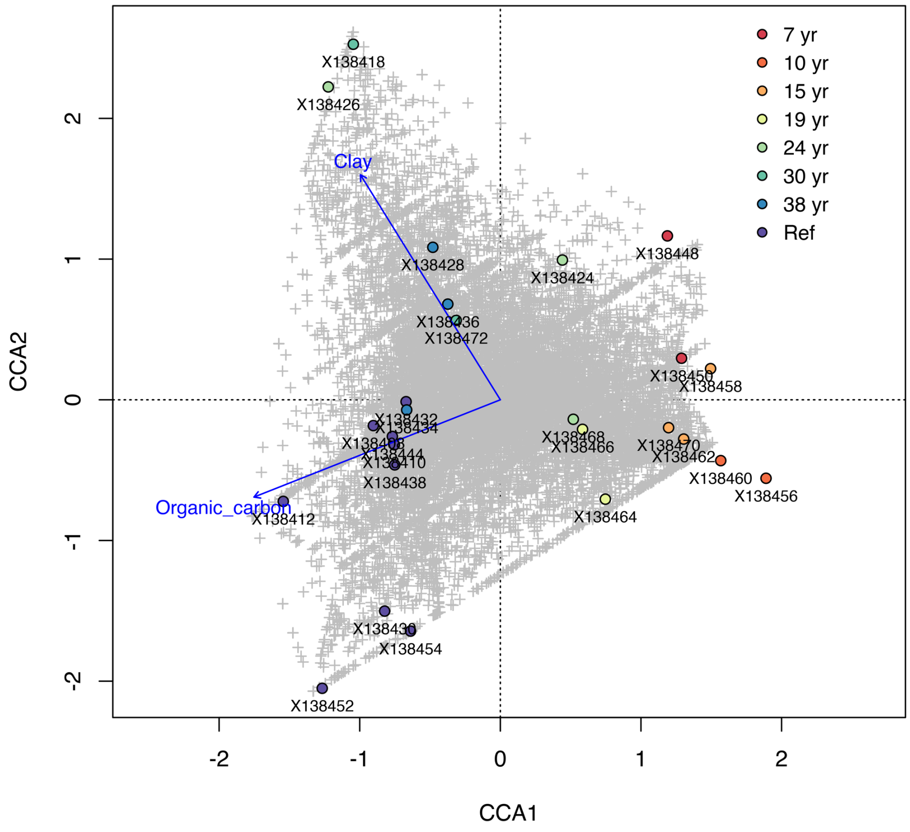


FIGURE S17. Constrained correspondence analysis (CCA) plot for Eneabba. Sample microbiota associate with scaled soil variables: organic carbon (mass %) and clay content (% in the fine earth fraction, < 2µm).

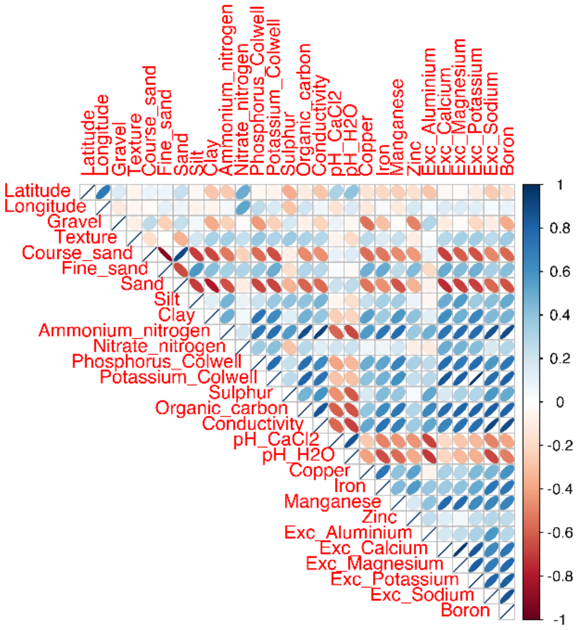


FIGURE S18. Correlation plot for soil sample/site variables at Worsley.

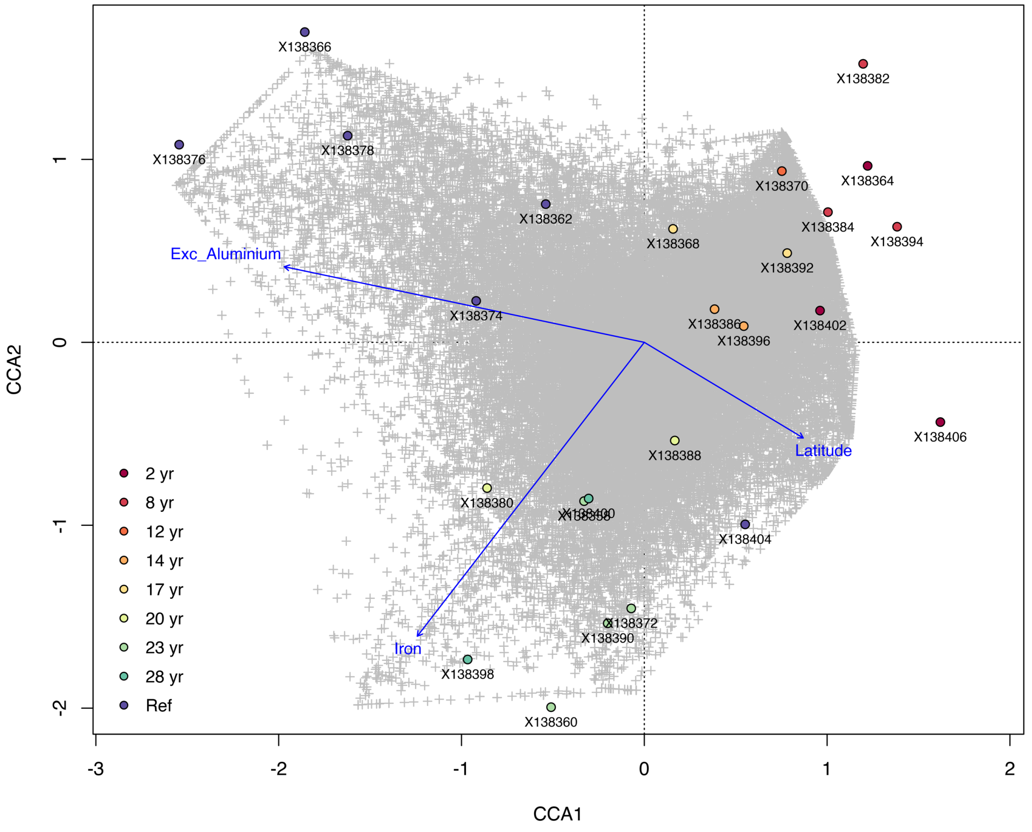


FIGURE S19. Constrained correspondence analysis (CCA) plot for Worsley. Sample bacterial communities associate with scaled soil and site variables: exchangeable aluminium (meq/100g), iron (DTPA extraction; mg/kg), and latitude (decimal degrees).

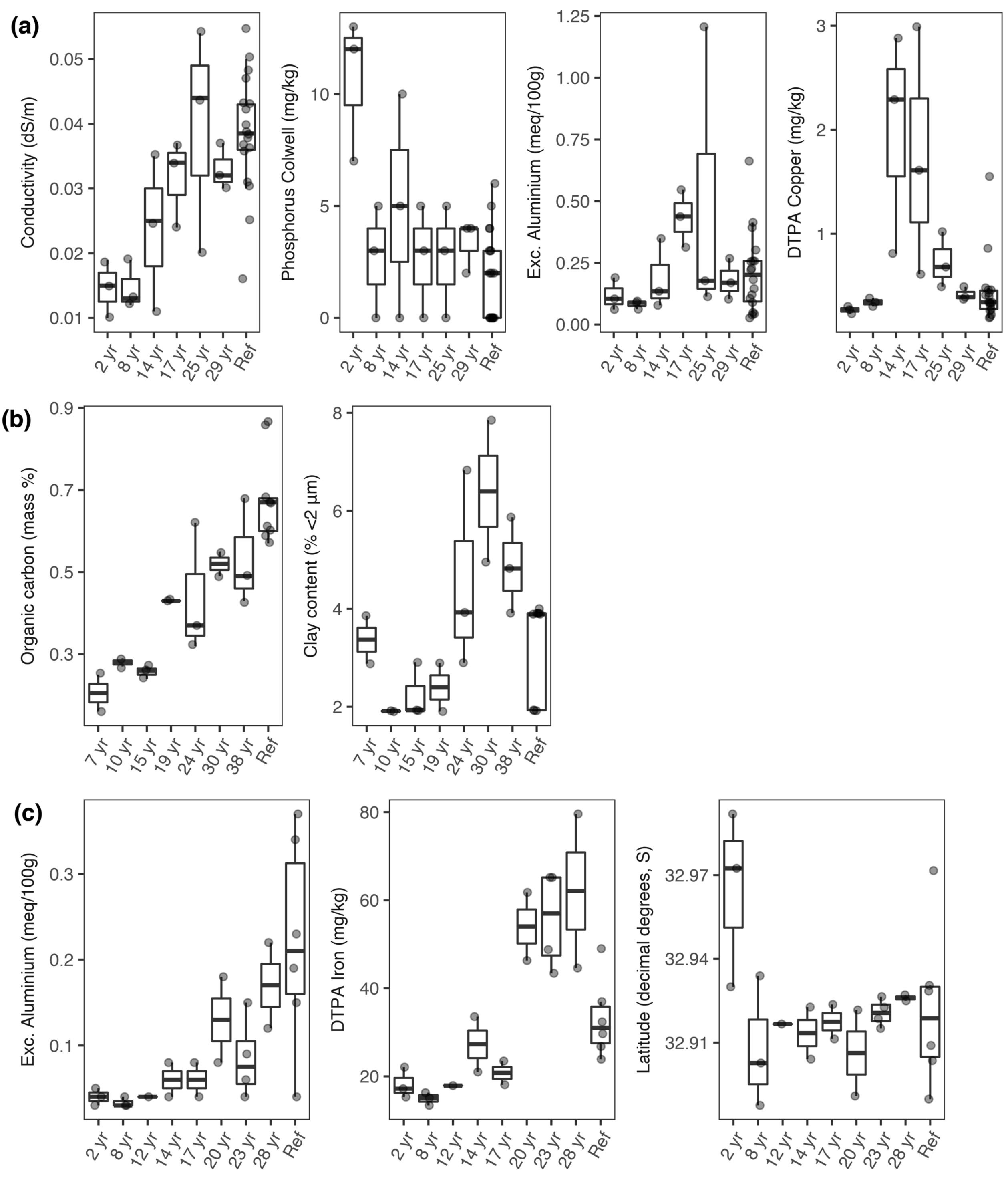


FIGURE S20. Variation in soil and landscape variables that associated with bacterial communities at (a) Huntly, (b) Eneabba, and (c) Worsley, as identified from constrained correspondence analysis (see Figures S15, S17, S19).

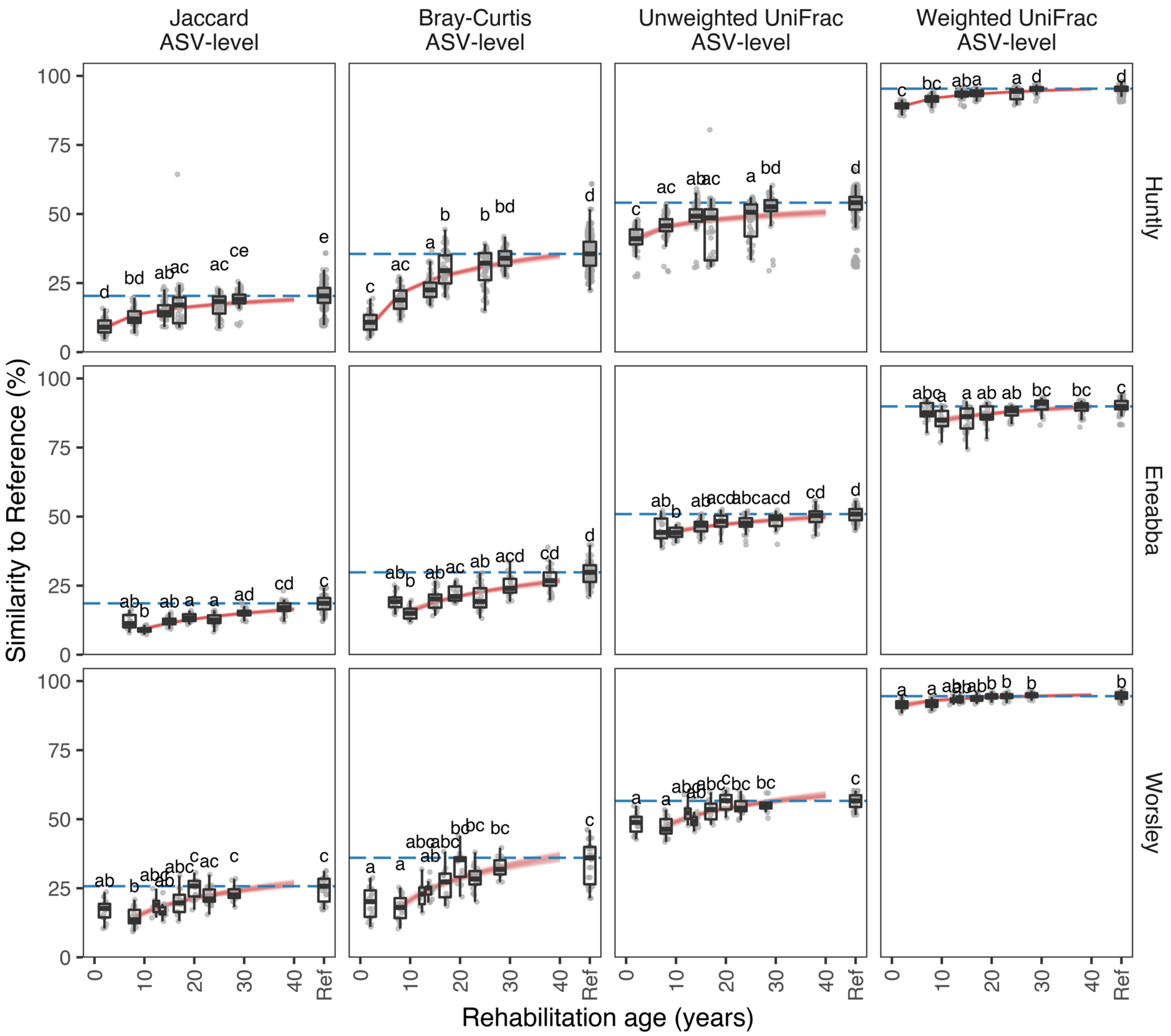


FIGURE S21. Modelled rehabilitation trajectories based on surface soil bacterial community similarity to reference data using Jaccard, Bray-Curtis, Unweighted UniFrac and Weighted UniFrac measures, for the Huntly, Eneabba, and Worsley minesites. Boxplots display the distribution of similarity values across rehabilitation ages (groups not sharing a letter are significantly different). Blue dotted lines denote the median similarity among references. Red lines represent logarithmic models for the change in similarity to reference with rehabilitation age, based on bootstrap resampling and modelling (B=100).

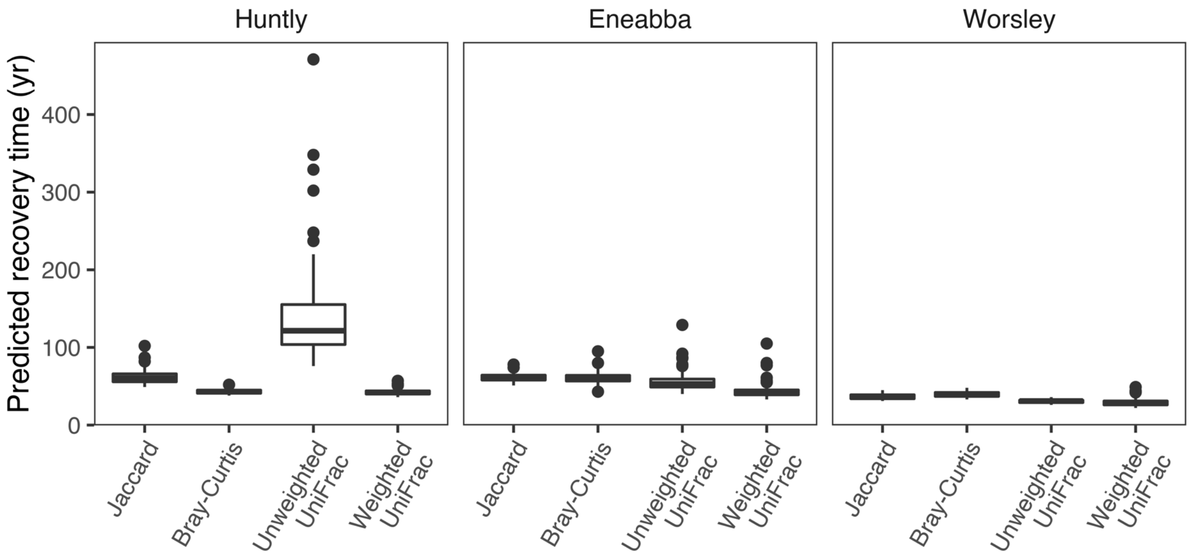


FIGURE S22. Predicted recovery times for soil bacterial ASVs to reach the target similarity to reference (= median of among-reference similarity values), for Huntly, Enneabba, and Worsley, considering Jaccard, Bray-Curtis, Unweighted UniFrac and Weighted UniFrac measures, based on bootstrap (B=100) logarithmic models (see SI Appendix, Table S6 for values).

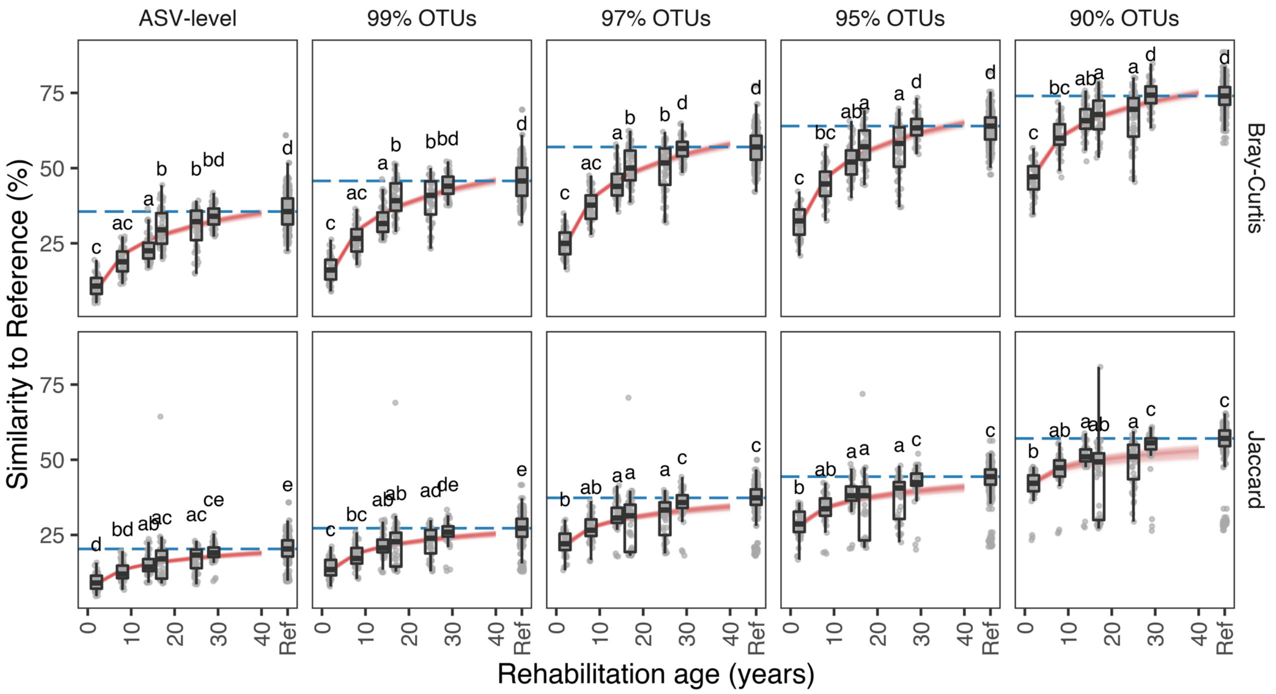


FIGURE S23. Modelled rehabilitation trajectories based on surface soil bacterial community similarity to reference samples for the Huntly minesite, comparing ASV-level data with 99%, 97%, 95%, and 90%-identity clustered OTUs for Jaccard and Bray-Curtis measures. Boxplots display the distribution of similarity values (groups not sharing a letter are significantly different) across a range of rehabilitation ages. Blue dotted lines denote the assumed target, equal to the median % similarity among reference soils. Red lines represent logarithmic models for the change in % similarity to reference with rehabilitation age based on bootstrap resampling and modelling (B=100). Sample sizes for the rehabilitation age groups (used to produce distance data) are described in section 2.1.

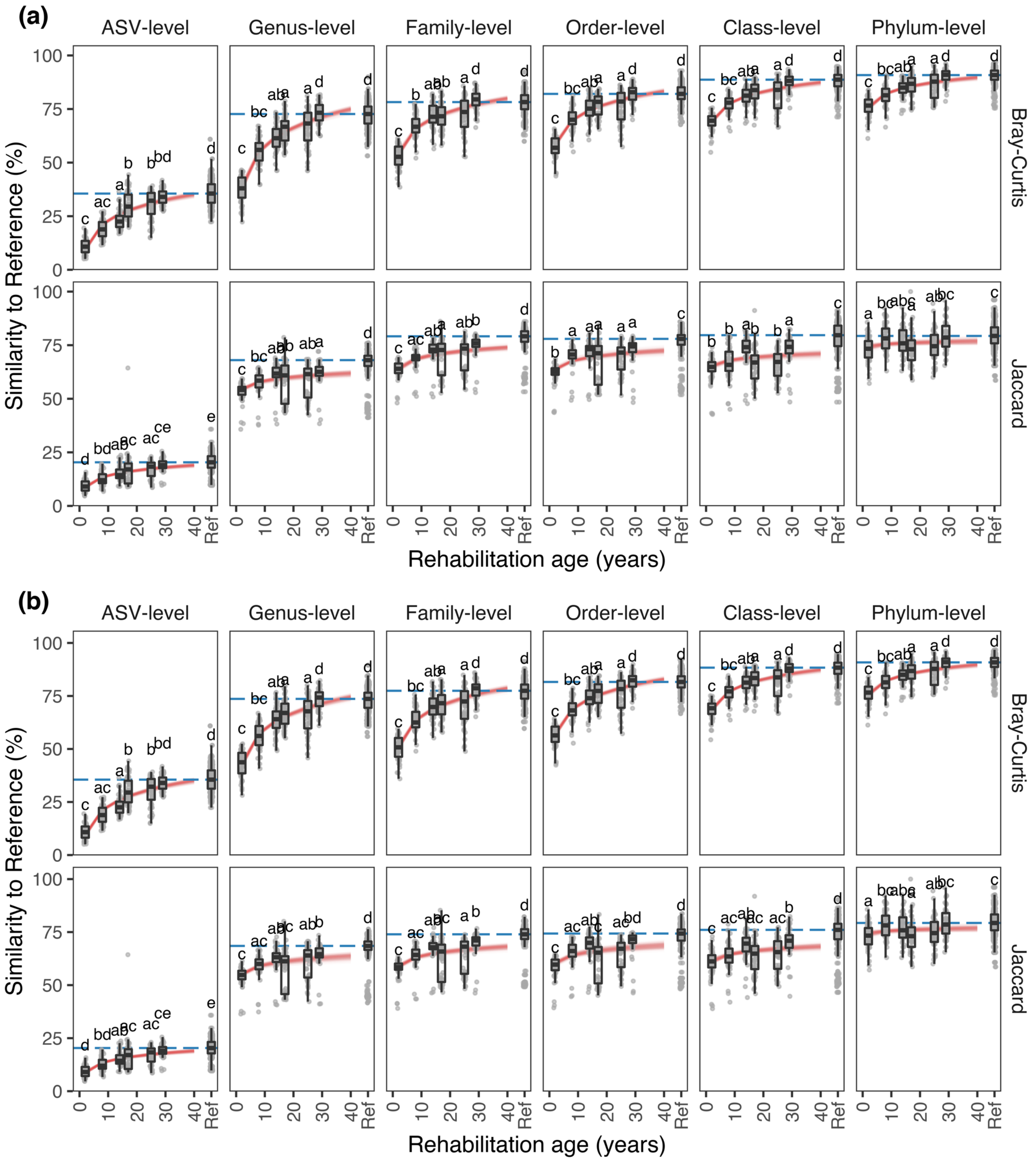


FIGURE S24. Modelled rehabilitation trajectories based on surface soil bacterial community similarity to reference samples for the Huntly minesite, comparing ASV-level data with taxa grouped at higher taxonomic ranks, with (A) pruning and (B) non-pruning of unclassified taxa within respective ranks, for Jaccard and Bray-Curtis measures. Boxplots display the distribution of similarity values (groups not sharing a letter are significantly different) across a range of rehabilitation ages. Blue dotted lines denote the assumed target, equal to the median % similarity among reference soils. Red lines represent logarithmic models for the change in % similarity to reference with rehabilitation age based on bootstrap resampling and modelling (B=100). Sample sizes for the rehabilitation age groups (used to produce distance data) are described in section 2.1. The relative abundance of phyla, classes, and orders present are indicated in heatmaps, see Figures S5-S13.

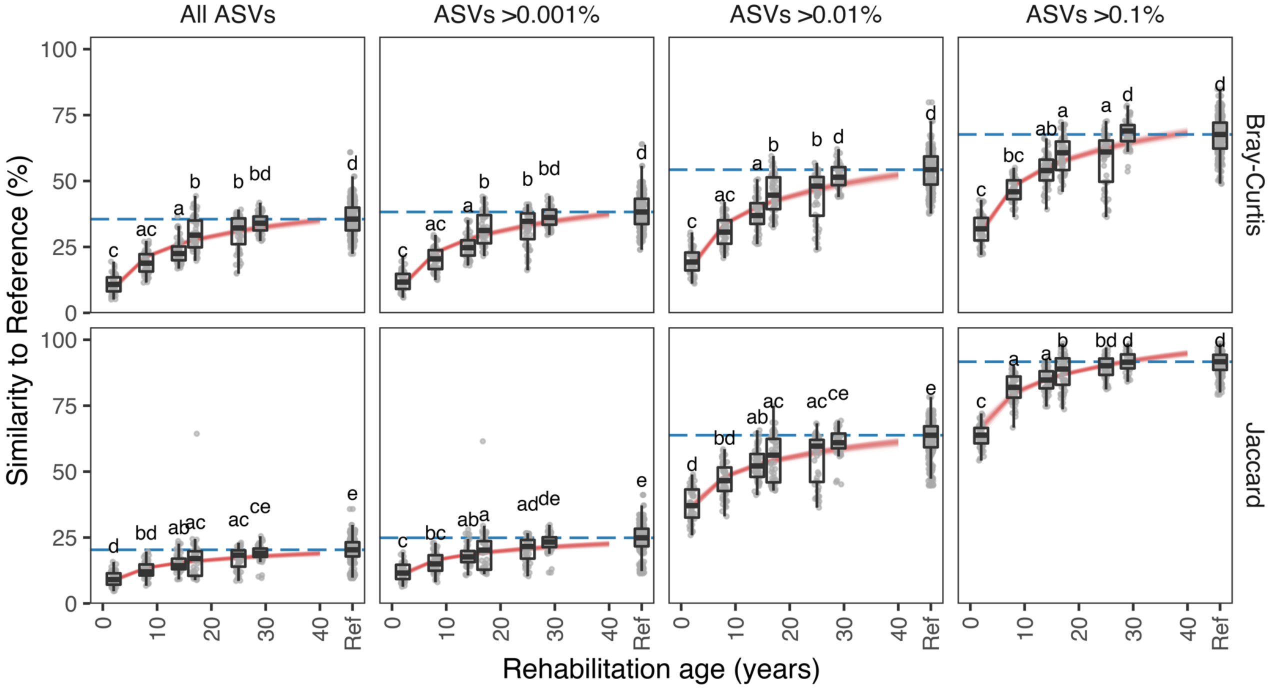


FIGURE S25. Modelled rehabilitation trajectories based on surface soil bacterial community similarity to reference samples for the Huntly minesite, comparing all ASV-level data with filtered data for ASVs > 0.001%, > 0.01%, and > 0.1% relative abundance, for Jaccard and Bray-Curtis measures. Boxplots display the distribution of similarity values (groups not sharing a letter are significantly different) across a range of rehabilitation ages. Blue dotted lines denote the assumed target, equal to the median % similarity among reference soils. Red lines represent logarithmic models for the change in % similarity to reference with rehabilitation age based on bootstrap resampling and modelling (B=100). Sample sizes for the rehabilitation age groups (used to produce distance data) are described in section 2.1.

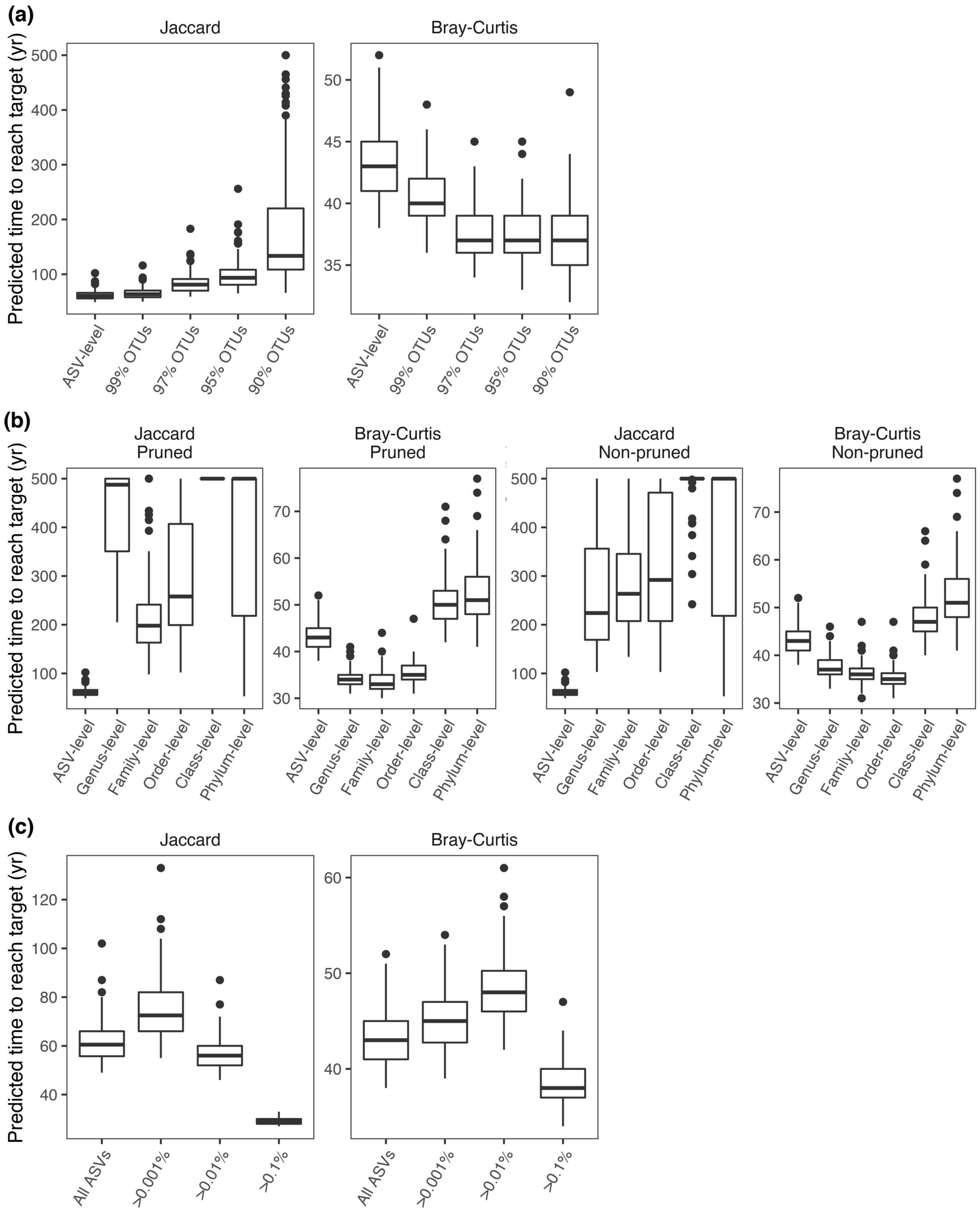


FIGURE S26. Predicted time to reach target bacterial community similarity to references, for alternative pre-processing options (a) grouping by sequence similarity (i.e. clustered OTUs); (b) grouping by taxonomic ranks, with pruning and non-pruning of unclassified taxa at each rank; and (c) filtering ASVs present at >0.001%, >0.01%, >0.1% relative abundance. Note the panels have different y-axis limits to highlight variability in outcomes.

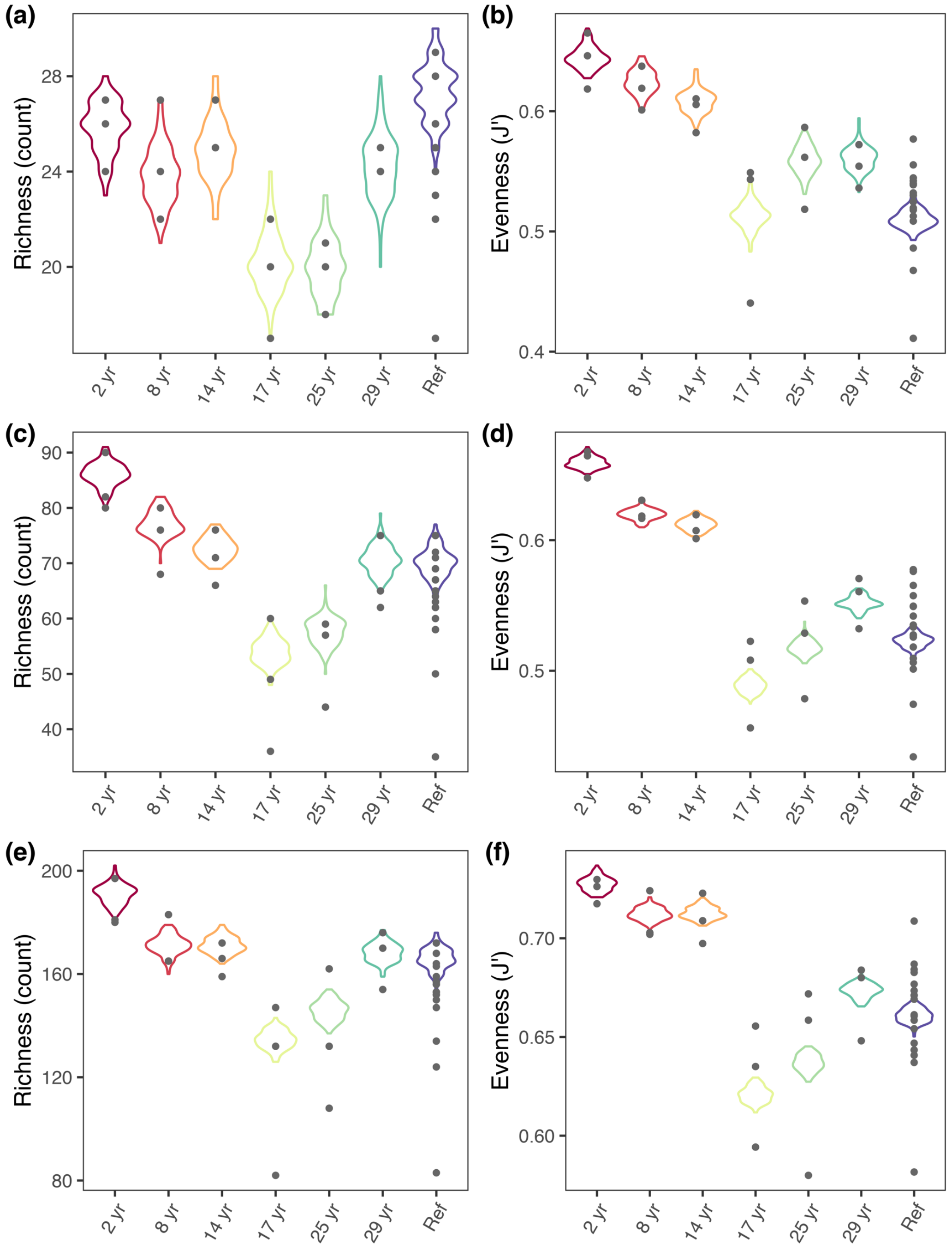


FIGURE S27. Huntly minesite data: phyla-level (a) richness, and (b) evenness; class-level (c) richness, and (d) evenness; order-level (e) richness, and (f) evenness. Violin-plot distributions represent the overall richness or evenness respectively, within a rehabilitation age group, calculated using merged-sample bootstrap resampling (B = 100). Grey dots represent richness or evenness respectively within each sample calculated from one-off rarefying to the minimum sample sequence read depth (i.e., 17,485 sequences).

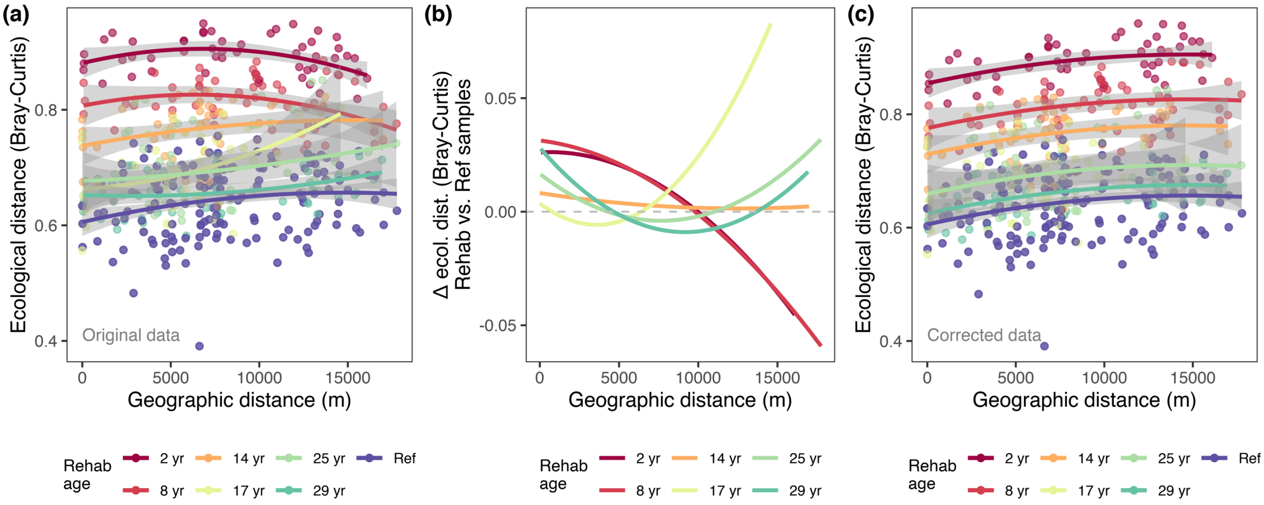


FIGURE S28. Exploring spatial autocorrelation in the Huntly dataset based on Bray-Curtis distance measures. (a) Ecological distance to reference versus geographic distance to reference for rehabilitation age groups. (b) Mean-centred difference in ecological distance to reference between rehabilitation age groups and among references. (c) Corrected ecological distance to reference versus geographic distance for rehabilitation age groups, to match the slope-trend of ecological to geographic distances as found among references.

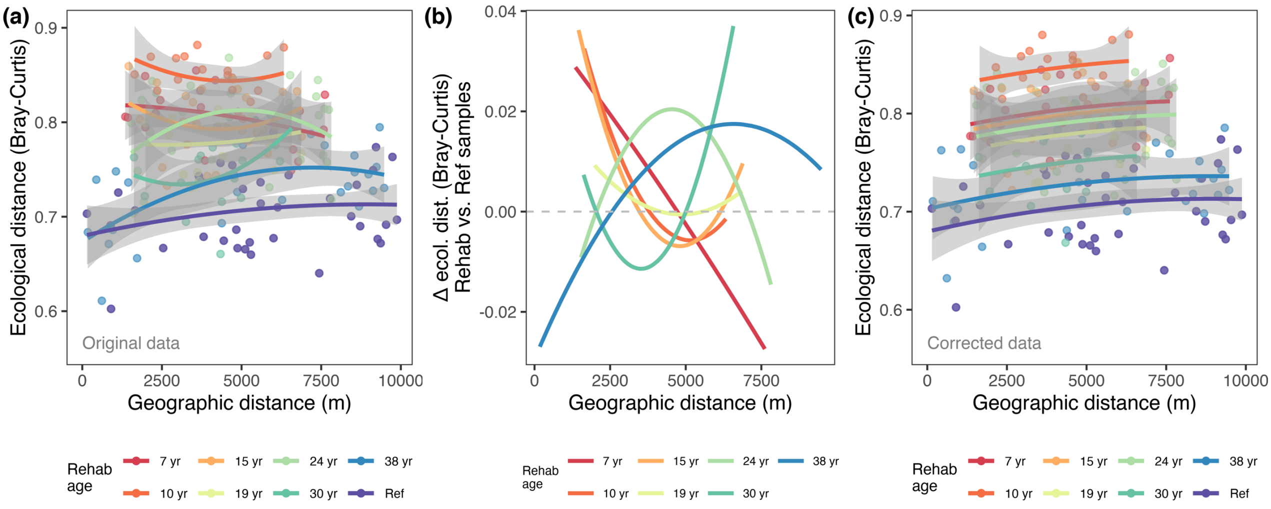


FIGURE S29. Exploring spatial autocorrelation in the Eneabba dataset based on Bray-Curtis distance measures. (a) Ecological distance to reference versus geographic distance to reference for rehabilitation age groups. (b) Mean-centred difference in ecological distance to reference between rehabilitation age groups and among references. (c) Corrected ecological distance to reference versus geographic distance for rehabilitation age groups, to match the slope-trend of ecological to geographic distances as found among references.

TABLE S1. Huntly site data. Further detailed data is available via the Australian Microbiome (AM) initiative data portal (https://data.bioplatforms.com/organization/about/australian-microbiome)

| Sample ID[†] | AMI sample ID | Longitude (Decimal degrees; GDA94) | Latitude (Decimal degrees; GDA94) | Rehabilitation age |
|---|---|---|---|---|
| 39041 | 102.100.100/39041 | 116.14246 | -32.51827 | 2 yr |
| 39043 | 102.100.100/39043 | 116.14239 | -32.51761 | Ref |
| 39045 | 102.100.100/39045 | 116.18371 | -32.52046 | 2 yr |
| 39047 | 102.100.100/39047 | 116.18426 | -32.51976 | Ref |
| 39049 | 102.100.100/39049 | 116.17192 | -32.52362 | 2 yr |
| 39051 | 102.100.100/39051 | 116.1718 | -32.5227 | Ref |
| 39053 | 102.100.100/39053 | 116.15891 | -32.53487 | 8 yr |
| 39055 | 102.100.100/39055 | 116.1582 | -32.53369 | Ref |
| 39057 | 102.100.100/39057 | 116.22374 | -32.57229 | 8 yr |
| 39059 | 102.100.100/39059 | 116.22408 | -32.5729 | Ref |
| 39061 | 102.100.100/39061 | 116.18417 | -32.58888 | 8 yr |
| 39063 | 102.100.100/39063 | 116.18457 | -32.58827 | Ref |
| 39065 | 102.100.100/39065 | 116.08824 | -32.61953 | 25 yr |
| 39067 | 102.100.100/39067 | 116.0879 | -32.62009 | Ref |
| 39069 | 102.100.100/39069 | 116.06979 | -32.62008 | 29 yr |
| 39071 | 102.100.100/39071 | 116.06943 | -32.61966 | Ref |
| 39073 | 102.100.100/39073 | 116.05283 | -32.60527 | 29 yr |
| 39075 | 102.100.100/39075 | 116.0519 | -32.60501 | Ref |
| 39077 | 102.100.100/39077 | 116.0462 | -32.59863 | 29 yr |
| 39079 | 102.100.100/39079 | 116.04491 | -32.59757 | Ref |
| 39081 | 102.100.100/39081 | 116.03674 | -32.59458 | 25 yr |
| 39083 | 102.100.100/39083 | 116.03629 | -32.59432 | Ref |
| 39085 | 102.100.100/39085 | 116.04454 | -32.55292 | 14 yr |
| 39087 | 102.100.100/39087 | 116.04415 | -32.55298 | Ref |
| 39089 | 102.100.100/39089 | 116.04864 | -32.55939 | 14 yr |
| 39091 | 102.100.100/39091 | 116.04851 | -32.55975 | Ref |
| 39093 | 102.100.100/39093 | 116.06857 | -32.57056 | 17 yr |
| 39095 | 102.100.100/39095 | 116.06907 | -32.57004 | Ref |
| 39097 | 102.100.100/39097 | 116.08275 | -32.59874 | 25 yr |
| 39099 | 102.100.100/39099 | 116.08204 | -32.59924 | Ref |
| 39161 | 102.100.100/39161 | 116.12112 | -32.56681 | 17 yr |
| 39163 | 102.100.100/39163 | 116.11913 | -32.56602 | Ref |
| 39165 | 102.100.100/39165 | 116.11904 | -32.56591 | 14 yr |
| 39191 | 102.100.100/39191 | 116.11904 | -32.56591 | Ref |
| 39193 | 102.100.100/39193 | 116.11904 | -32.56591 | 17 yr |
| 39195 | 102.100.100/39195 | 116.11904 | -32.56591 | Ref |

[†]Note: SI figures may include sample IDs with the prefix ‘X’ for data handling purposes.

TABLE S2. Eneabba site data. Further detailed data is available via the Australian Microbiome (AM) initiative data portal (https://data.bioplatforms.com/organization/about/australian-microbiome)

| Sample ID† | AMI sample ID | Longitude (Decimal degrees; GDA94) | Latitude (Decimal degrees; GDA94) | Rehabilitation age |
|---|---|---|---|---|
| 138408 | 102.100.100/138408 | 115.30685 | -29.85854 | Ref |
| 138410 | 102.100.100/138410 | 115.30768 | -29.85744 | Ref |
| 138412 | 102.100.100/138412 | 115.30451 | -29.90079 | Ref |
| 138418 | 102.100.100/138418 | 115.29352 | -29.91182 | 30 yr |
| 138424 | 102.100.100/138424 | 115.28249 | -29.92179 | 24 yr |
| 138426 | 102.100.100/138426 | 115.27987 | -29.92265 | 24 yr |
| 138428 | 102.100.100/138428 | 115.28059 | -29.93926 | 38 yr |
| 138430 | 102.100.100/138430 | 115.27763 | -29.94226 | Ref |
| 138432 | 102.100.100/138432 | 115.27332 | -29.93523 | Ref |
| 138434 | 102.100.100/138434 | 115.2729 | -29.9337 | 38 yr |
| 138436 | 102.100.100/138436 | 115.27376 | -29.92973 | 38 yr |
| 138438 | 102.100.100/138438 | 115.26441 | -29.93275 | Ref |
| 138444 | 102.100.100/138444 | 115.26823 | -29.85698 | Ref |
| 138448 | 102.100.100/138448 | 115.27285 | -29.87432 | 7 yr |
| 138450 | 102.100.100/138450 | 115.27433 | -29.8739 | 7 yr |
| 138452 | 102.100.100/138452 | 115.26273 | -29.86594 | Ref |
| 138454 | 102.100.100/138454 | 115.25452 | -29.88762 | Ref |
| 138456 | 102.100.100/138456 | 115.2714 | -29.88568 | 10 yr |
| 138458 | 102.100.100/138458 | 115.26952 | -29.88615 | 15 yr |
| 138460 | 102.100.100/138460 | 115.27662 | -29.89009 | 10 yr |
| 138462 | 102.100.100/138462 | 115.26917 | -29.89772 | 15 yr |
| 138464 | 102.100.100/138464 | 115.26154 | -29.90438 | 19 yr |
| 138466 | 102.100.100/138466 | 115.27548 | -29.90391 | 19 yr |
| 138468 | 102.100.100/138468 | 115.26772 | -29.91854 | 24 yr |
| 138470 | 102.100.100/138470 | 115.27269 | -29.91134 | 15 yr |
| 138472 | 102.100.100/138472 | 115.28455 | -29.89748 | 30 yr |

†Note: SI figures may include sample IDs with the prefix ‘X’ for data handling purposes.

TABLE S3. Worsley site data. Further detailed data is available via the Australian Microbiome (AM) initiative data portal (https://data.bioplatforms.com/organization/about/australian-microbiome)

| Sample ID† | AM sample ID | Longitude (Decimal degrees; GDA94) | Latitude (Decimal degrees; GDA94) | Rehabilitation age |
|---|---|---|---|---|
| 138358 | 102.100.100/138358 | 116.46573 | -32.9227 | 23 yr |
| 138360 | 102.100.100/138360 | 116.47172 | -32.92646 | 23 yr |
| 138362 | 102.100.100/138362 | 116.47382 | -32.92846 | Ref |
| 138364 | 102.100.100/138364 | 116.45609 | -32.92999 | 2 yr |
| 138366 | 102.100.100/138366 | 116.45549 | -32.93049 | Ref |
| 138368 | 102.100.100/138368 | 116.45389 | -32.91135 | 17 yr |
| 138370 | 102.100.100/138370 | 116.45506 | -32.91668 | 12 yr |
| 138372 | 102.100.100/138372 | 116.44543 | -32.91866 | 23 yr |
| 138374 | 102.100.100/138374 | 116.44527 | -32.90898 | Ref |
| 138376 | 102.100.100/138376 | 116.4481 | -32.90357 | Ref |
| 138378 | 102.100.100/138378 | 116.43892 | -32.88975 | Ref |
| 138380 | 102.100.100/138380 | 116.44391 | -32.89086 | 20 yr |
| 138382 | 102.100.100/138382 | 116.4481 | -32.88755 | 8 yr |
| 138384 | 102.100.100/138384 | 116.42808 | -32.90269 | 8 yr |
| 138386 | 102.100.100/138386 | 116.4337 | -32.9041 | 14 yr |
| 138388 | 102.100.100/138388 | 116.43233 | -32.92172 | 20 yr |
| 138390 | 102.100.100/138390 | 116.43416 | -32.9151 | 23 yr |
| 138392 | 102.100.100/138392 | 116.43931 | -32.9237 | 17 yr |
| 138394 | 102.100.100/138394 | 116.42781 | -32.93388 | 8 yr |
| 138396 | 102.100.100/138396 | 116.44985 | -32.92279 | 14 yr |
| 138398 | 102.100.100/138398 | 116.47131 | -32.92499 | 28 yr |
| 138400 | 102.100.100/138400 | 116.46324 | -32.92707 | 28 yr |
| *138402 | 102.100.100/138402 | 116.48289 | -32.97245 | 2 yr |
| *138404 | 102.100.100/138404 | 116.48354 | -32.97165 | Ref |
| *138406 | 102.100.100/138406 | 116.48699 | -32.99188 | 2 yr |

Notes: †SI figures may include sample IDs with the prefix 'X' for data handling purposes.
*These southernmost sites were excluded in the 'filtered' analysis of spatial autocorrelation.

TABLE S4. Total cleaned and rarefied numbers of taxa and sequences for minesites and pre-processing options.

| Pre-processing option | Taxa (level of grouping) | Cleaned data No. of taxa [No. of sequences] | Rarefied data No. of taxa [No. of sequences] |
|---|---|---|---|
| Huntly minesite (n = 36 samples) | | | |
| Standard measures (Jaccard, Bray-Curtis, UniFrac) | ASVs | 31,746 taxa [1,772,249 sequences] | 30,751 taxa [629,460 sequences] |
| Compositional / Aitchison measure | ASVs | 25,720 taxa [1,723,759 sequences] | N/A |
| i) Grouping by sequence similarity (i.e. clustering into OTUs) | OTUs based on 99% identity clusters | 18,062 taxa [1,772,249 sequences] | 17,584 taxa [629,460 sequences] |
| | OTUs based on 97% identity clusters | 8663 taxa [1,772,249 sequences] | 8461 taxa [629,460 sequences] |
| | OTUs based on 95% identity clusters | 4868 taxa [1,772,249 sequences] | 4772 taxa [629,460 sequences] |
| | OTUs based on 90% identity clusters | 1512 taxa [1,772,249 sequences] | 1489 taxa [629,460 sequences] |
| iia) Grouping at higher taxonomic levels (with retention of unclassified taxa at the next available level) | Genera (non-pruned) | 771 taxa [1,772,249 sequences] | 767 taxa [629,460 sequences] |
| | Families (non-pruned) | 404 taxa [1,772,249 sequences] | 400 taxa [629,460 sequences] |
| | Orders (non-pruned) | 266 taxa [1,772,249 sequences] | 265 taxa [629,460 sequences] |
| | Classes (non-pruned) | 110 taxa [1,772,249 sequences] | 109 taxa [629,460 sequences] |
| | Phyla (non-pruned) | 33 taxa [1,772,249 sequences] | 33 taxa [629,460 sequences] |
| iib) Grouping at higher taxonomic levels (with removal of unclassified taxa) | Genera (pruned) (42.3 % of sequences excluded) | 457 taxa [1,022,280 sequences] | 454 taxa [373,608 sequences] |
| | Families (pruned) (20.2 % of sequences excluded) | 243 taxa [1,413,678 sequences] | 242 taxa [507,924 sequences] |
| | Orders (pruned) (3.2 % of sequences excluded) | 202 taxa [1,716,291 sequences] | 200 taxa [622,764 sequences] |
| | Classes (pruned) (0.6 % of sequences excluded) | 94 taxa [1,762,184 sequences] | 92 taxa [628,128 sequences] |
| | Phyla (pruned) (Unclassified phyla were already removed to obtain cleaned data) | 33 taxa [1,772,249 sequences] | 33 taxa [629,460 sequences] |
| iii) Eliminating rare taxa | ASVs >0.001% | 17943 taxa [1,645,918 sequences] | 17941 taxa [603,072 sequences] |
| | ASVs >0.01% | 1338 taxa [868,441 sequences] | 1338 taxa [348,876 sequences] |
| | ASVs >0.1% | 72 taxa [320,029 sequences] | 72 taxa [49,752 sequences] |
| Eneabba minesite (n = 26 samples) | | | |
| Standard measures (Jaccard, Bray-Curtis, UniFrac) | ASVs | 33,636 taxa [2,155,211 sequences] | 27,115 taxa [263,692 sequences] |
| Compositional / Aitchison measure | ASVs | 24117 taxa [2,042,214 sequences] | N/A |
| Worsley minesite (n = 25 samples) | | | |
| Standard measures (Jaccard, Bray-Curtis, UniFrac) | ASVs | 54,671 taxa [2,049,625 sequences] | 54,327 taxa [1,353,050 sequences] |
| Worsley minesite (filtered, excl. southern) (n = 22 samples) | | | |
| Standard measures (Jaccard, Bray-Curtis, UniFrac) | ASVs | 54,671 taxa [1,839,173 sequences] | 53404 taxa [1,190,684 sequences] |
| Compositional / Aitchison measure | ASVs | 43,598 taxa [1,782,724 sequences] | N/A |

TABLE S5. Similarity values among reference samples within each minesite, showing median and interquartile range of similarity to reference (%)

(a) Based on ASV-level data and comparison of Jaccard, Bray-Curtis, Unweighted UniFrac, Weighted UniFrac, Aitchison measures

| Study location | Jaccard ASV-level (%) | Bray-Curtis ASV-level (%) | Unweighted UniFrac ASV-level (%) | Weighted UniFrac ASV-level (%) | Aitchison ASV-level (%) |
|---|---|---|---|---|---|
| Huntly | 20.3 (17.8-23.2) | 35.6 (31.3-39.9) | 54.1 (51.6-56.2) | 95.4 (94.6-96.1) | 22.1 (18.1-27.8) |
| Eneabba | 18.6 (16.4-20.5) | 29.8 (26.3-32.4) | 50.9 (48.7-52.7) | 89.9 (88.9-91.8) | 20.7 (17.3-22.3) |
| Worsley | 25.7 (20.1-28.4) | 36.0 (26.5-40.0) | 56.6 (54.4-58.7) | 94.5 (93.6-95.9) | NA |
| Worsley (*filtered; excl. southern) | N/A | 38.2 (36.0-42.9) | N/A | N/A | 21.6 (18.4-24.7) |

*Worsley filtered data have the southernmost sites 138402, 138404, 138406 excluded.

(b) Based on ASV-level, 99%, 97%, 95%, 90%-identity clustered OTUs, with Jaccard and Bray-Curtis measures.

| Distance measure | ASV-level (%) | 99% OTUs (%) | 97% OTUs (%) | 95% OTUs (%) | 90% OTUs (%) |
|---|---|---|---|---|---|
| Bray-Curtis | 35.6 (31.3-39.9) | 45.7 (40.8-50.1) | 57.0 (52.8-60.8) | 64.0 (59.5-66.9) | 74.0 (71.1-76.9) |
| Jaccard | 20.3 (17.8-23.2) | 27.2 (24.4-30.4) | 37.3 (35.0-40.1) | 44.4 (41.7-46.8) | 57.1 (54.5-59.7) |

(c) Based on ASV, genus, family, order, class, and phylum-level grouping (showing both pruning and non-pruning of unclassified taxa), with Jaccard and Bray-Curtis measures.

| Distance measure | ASV (%) | Genus (%) | Family (%) | Order (%) | Class (%) | Phylum (%) |
|---|---|---|---|---|---|---|
| Bray-Curtis (Pruned) | 35.6 (31.3-39.9) | 72.7 (68.2-76.2) | 78.2 (74.9-81.5) | 82.1 (79.7-85.2) | 88.7 (85.6-91.0) | 90.8 (88.7-93.0) |
| Bray-Curtis (Non-pruned) | 35.6 (31.3-39.9) | 73.6 (69.4-76.5) | 77.4 (73.7-80.4) | 81.6 (79.0-84.3) | 88.3 (85.4-90.7) | 90.8 (88.7-93.0) |
| Jaccard (Pruned) | 20.3 (17.8-23.2) | 68.0 (65.2-70.1) | 79.1 (76.6-81.5) | 77.9 (75.2-79.7) | 79.7 (74.6-84.1) | 79.3 (75.9-83.3) |
| Jaccard (Non-pruned) | 20.3 (17.8-23.2) | 68.5 (66.6-70.6) | 74.0 (71.7-76.4) | 74.3 (71.0-76.2) | 76.1 (71.6-79.0) | 79.3 (75.9-83.3) |

(d) Based on all ASVs and filtered ASVs present at >0.001%, >0.01%, >0.1% relative abundance, with Jaccard and Bray-Curtis measures.

| Distance measure | All ASVs (%) | >0.001% (%) | >0.01% (%) | >0.1% (%) |
|---|---|---|---|---|
| Bray-Curtis | 35.6 (31.3-39.9) | 38.3 (33.7-43.3) | 54.3 (48.7-59.3) | 67.7 (62.3-72.1) |
| Jaccard | 20.3 (17.8-23.2) | 24.9 (21.7-28.3) | 63.8 (59.2-67.2) | 91.7 (88.6-94.3) |

TABLE S6. Predicted times for bacteria in rehabilitated soils to reach the target % similarity to reference (taken as the median of within-reference % similarity values)—based on comparison data as described below, with Jaccard and Bray-Curtis similarity measures. The tables displays median ($2.5^{th}$ percentile, $97.5^{th}$ percentile) predicted years based on results from bootstrap (B=100) logarithmic models.

(a) Based on ASV-level, Jaccard, Bray-Curtis, Unweighted-UniFrac, Weighted-UniFrac, and Aitchison measures.

| Minesite | Jaccard ASV-level (yr) | Bray-Curtis ASV-level (yr) | Unweighted UniFrac ASV-level (yr) | Weighted UniFrac ASV-level (yr) | Aitchison ASV-level (yr) |
|---|---|---|---|---|---|
| Huntly | 61 (51, 81) | 43 (39, 50) | 122 (78, 316) | 42 (38, 51) | 31 (27, 36) |
| Eneabba | 61 (53, 73) | 60 (51, 73) | 53 (41, 87) | 41 (34, 69) | 50 (44, 59) |
| Worsley | 36 (31, 43) | 39 (34, 46) | 31 (27, 36) | 29 (23, 43) | N/A |
| Worsley (*filtered; excl. southern) | N/A | 44 (37, 60) | N/A | N/A | 53 (43, 71) |

*Worsley filtered data have the southernmost sites 138402, 138404, 138406 excluded

(b) Huntly only, based on ASV-level, 99%, 97%, 95%, 90%-identity clustered OTUs.

| Distance measure | ASV-level (yr) | 99% OTUs (yr) | 97% OTUs (yr) | 95% OTUs (yr) | 90% OTUs (yr) |
|---|---|---|---|---|---|
| Bray-Curtis | 43 (39, 50) | 40 (37, 46) | 37 (34, 43) | 37 (34, 42) | 37 (32, 43) |
| Jaccard | 61 (51, 81) | 64 (52, 89) | 81 (61, 130) | 94 (66, 177) | 134 (71, 483) |

(c) Huntly only, based on ASV, genus, family, order, class, and phylum-level grouping (with results for pruning and non-pruning of unclassified taxa).

| Distance measure | ASV-level (yr) | Genus-level (yr) | Family-level (yr) | Order-level (yr) | Class-level (yr) | Phylum-level (yr) |
|---|---|---|---|---|---|---|
| Bray-Curtis (Pruned) | 43 (39, 50) | 34 (31, 39) | 33 (30, 39) | 35 (32, 40) | 50 (43, 63) | 51 (43, 68) |
| Bray-Curtis (Non-pruned) | 43 (39, 50) | 37 (33, 44) | 36 (32, 42) | 35 (32, 40) | 47 (40, 58) | 51 (43, 68) |
| Jaccard (Pruned) | 61 (51, 81) | 488 (219, >500) | 198 (118, 421) | 258 (142, >500) | >500 (>500, >500) | >500 (61, >500) |
| Jaccard (Non-pruned) | 61 (51, 81) | 224 (111, >500) | 264 (146, >500) | 292 (124, >500) | >500 (361, >500) | >500 (61, >500) |

TABLE S6. (Continued next page)

TABLE S6. (Continued)

(d) Huntly only, based on all ASVs and filtered ASVs present at >0.001%, >0.01%, >0.1% relative abundance.

| Distance measure | All ASVs (yr) | >0.001% (yr) | >0.01% (yr) | >0.1% (yr) |
|---|---|---|---|---|
| Bray-Curtis | 43 (39, 50) | 45 (40, 52) | 48 (43, 57) | 38 (34, 43) |
| Jaccard | 61 (51, 81) | 73 (59, 106) | 56 (48, 71) | 29 (27, 32) |

(e) Based on original and spatial-autocorrelation-corrected ASV-level data.

| Study location | Original data Bray-Curtis (yr) | Corrected data Bray-Curtis (yr) |
|---|---|---|
| Huntly | 43 (39, 50) | 41 (37, 47) |
| Eneabba | 60 (51, 73) | 56 (48, 68) |
| Worsley | 39 (34, 46) | 50 (37, 87) |
| Worsley (*filtered; excl. southern) | 44 (37, 60) | 44 (36, 61) |

*Worsley filtered data have the southernmost sites 138402, 138404, 138406 excluded

Bolyen, E., Rideout, J.R., Dillon, M.R., Bokulich, N.A., Abnet, C.C., Al-Ghalith, G.A., Alexander, H., Alm, E.J., Arumugam, M., Asnicar, F., Bai, Y., Bisanz, J.E., Bittinger, K., Brejnrod, A., Brislawn, C.J., Brown, C.T., Callahan, B.J., Caraballo-Rodríguez, A.M., Chase, J., Cope, E.K., Da Silva, R., Diener, C., Dorrestein, P.C., Douglas, G.M., Durall, D.M., Duvallet, C., Edwardson, C.F., Ernst, M., Estaki, M., Fouquier, J., Gauglitz, J.M., Gibbons, S.M., Gibson, D.L., Gonzalez, A., Gorlick, K., Guo, J., Hillmann, B., Holmes, S., Holste, H., Huttenhower, C., Huttley, G.A., Janssen, S., Jarmusch, A.K., Jiang, L., Kaehler, B.D., Kang, K.B., Keefe, C.R., Keim, P., Kelley, S.T., Knights, D., Koester, I., Kosciolek, T., Kreps, J., Langille, M.G.I., Lee, J., Ley, R., Liu, Y.-X., Loftfield, E., Lozupone, C., Maher, M., Marotz, C., Martin, B.D., McDonald, D., McIver, L.J., Melnik, A.V., Metcalf, J.L., Morgan, S.C., Morton, J.T., Naimey, A.T., Navas-Molina, J.A., Nothias, L.F., Orchanian, S.B., Pearson, T., Peoples, S.L., Petras, D., Preuss, M.L., Pruesse, E., Rasmussen, L.B., Rivers, A., Robeson, M.S., Rosenthal, P., Segata, N., Shaffer, M., Shiffer, A., Sinha, R., Song, S.J., Spear, J.R., Swafford, A.D., Thompson, L.R., Torres, P.J., Trinh, P., Tripathi, A., Turnbaugh, P.J., Ul-Hasan, S., van der Hooft, J.J.J., Vargas, F., Vázquez-Baeza, Y., Vogtmann, E., von Hippel, M., Walters, W., Wan, Y., Wang, M., Warren, J., Weber, K.C., Williamson, C.H.D., Willis, A.D., Xu, Z.Z., Zaneveld, J.R., Zhang, Y., Zhu, Q., Knight, R. & Caporaso, J.G. (2019) Reproducible, interactive, scalable and extensible microbiome data science using QIIME 2. *Nature Biotechnology,* **37,** 852-857.

Borrett, R.J. (2020) The secrets in our soils: Using soil bacteria to monitor mine site restoration across a 40-year chronosequence in an arid environment in Western Australia. Honours thesis, Murdoch University.

Brundrett, M.C. & Abbott, L.K. (1991) Roots of Jarrah Forest Plants .I. Mycorrhizal Associations of Shrubs and Herbaceous Plants. *Australian journal of botany,* **39,** 445-457.

Callahan, B.J., McMurdie, P.J., Rosen, M.J., Han, A.W., Johnson, A.J.A. & Holmes, S.P. (2016) DADA2: High-resolution sample inference from Illumina amplicon data. *Nature Methods,* **13,** 581.

Churchward, H.M. & Dimmock, G.M. (1989) The soils and landforms of the northern jarrah forest. *The Jarrah Forest: A complex mediterranean ecosystem* (eds B. Dell, J.J. Havel & N. Malajczuk). Springer, Dordrecht, The Netherlands.

Cowling, R.M., Rundel, P.W., Lamont, B.B., Kalin Arroyo, M. & Arianoutsou, M. (1996) Plant diversity in mediterranean-climate regions. *Trends Ecol Evol,* **11,** 362-366.

Ducki, L.C. (2020) Soil fungi, but not bacteria, track vegetation reassembly across a 30-year restoration chronosequence in the northern jarrah forest, Western Australia. Honours thesis, Murdoch University.

Edgar, R.C. (2010) Search and clustering orders of magnitude faster than BLAST. *Bioinformatics,* **26,** 2460-2461.

Edgar, R.C. (2016) UNOISE2: improved error-correction for Illumina 16S and ITS amplicon sequencing. *bioRxiv***,** 081257.

Enright, N.J., Keith, D.A., Clarke, M.F. & Miller, B.P. (2012) Fire regimes in Australian sclerophyllous shrubby ecosystems: Heathlands, heathy woodlands and mallee woodlands. *Flammable Australia: Fire Regimes and Biodiversity in a Changing World* (eds R.J. Williams, A.M. Gill & R.A. Bradstock), pp. 215-235. CSIRO Publishing, Melbourne, Australia.

Enright, N.J. & Lamont, B.B. (1992) Survival, Growth and Water Relations of Banksia Seedlings on a Sand Mine Rehabilitation Site and Adjacent Scrub-Heath Sites. *The Journal of applied ecology,* **29,** 663-671.

Golos, P.J. & Dixon, K.W. (2014) Waterproofing Topsoil Stockpiles Minimizes Viability Decline in the Soil Seed Bank in an Arid Environment. *Restor Ecol,* **22,** 495-501.

Jasper, D.A. (2007) Beneficial soil microorganisms of the Jarrah forest and their recovery in bauxite Southwestern Australia. *Restoration Ecology,* **15,** S74-S84.

Jost, L. (2006) Entropy and diversity. *Oikos,* **113,** 363-375.

Katoh, K. & Standley, D.M. (2013) MAFFT Multiple Sequence Alignment Software Version 7: Improvements in Performance and Usability. *Molecular Biology and Evolution,* **30,** 772-780.

Koch, J.M. (2007) Alcoa's Mining and Restoration Process in South Western Australia. *Restoration Ecology,* **15,** S11-S16.

Koch, J.M. & Samsa, G.P. (2007) Restoring Jarrah Forest Trees after Bauxite Mining in Western Australia. *Restoration Ecology,* **15,** S17-S25.

Koch, J.M. & Ward, S.C. (1994) Establishment of Understorey Vegetation for Rehabilitation of Bauxite-mined Areas in the Jarrah Forest of Western Australia. *Journal of Environmental Management,* **41,** 1-15.

Kuhn, M. (2020) caret: Classification and Regression Training. R package version 6.0-86.

Lane, D.J. (1991) 16s/23s rrna sequencing. *Nucleic acid techniques in bacterial systematics* (eds E. Stackbrandt & M. Goodfellow). John Wiley and Sons, New York.

Lane, D.J., Pace, B., Olsen, G.J., Stahl, D.A., Sogin, M.L. & Pace, N.R. (1985) Rapid determination of 16 s ribosomal rna sequences for phylogenetic analyses. *Pro Natl Acad Sci,* **82**.

Liddicoat, C., Weinstein, P., Bissett, A., Gellie, N., Mills, J., Waycott, M. & Breed, M. (2019) Can bacterial indicators of a grassy woodland restoration inform ecosystem assessment and microbiota-mediated human health? *Environment International,* **129,** 105-117.

Magoč, T. & Salzberg, S.L. (2011) FLASH: fast length adjustment of short reads to improve genome assemblies. *Bioinformatics,* **27,** 2957-2963.

Manero, A., Standish, R. & Young, R. (2021) Mine completion criteria defined by best-practice: A global meta-analysis and Western Australian case studies. *Journal of Environmental Management,* **282,** 111912.

McMurdie, P.J. & Holmes, S. (2013) phyloseq: An R Package for Reproducible Interactive Analysis and Graphics of Microbiome Census Data. *PLoS One,* **8,** e61217.

Minh, B.Q., Schmidt, H.A., Chernomor, O., Schrempf, D., Woodhams, M.D., von Haeseler, A. & Lanfear, R. (2020) IQ-TREE 2: New Models and Efficient Methods for Phylogenetic Inference in the Genomic Era. *Molecular Biology and Evolution,* **37,** 1530-1534.

Mucina, L., Laliberte, E., Thiele, K.R., Dodson, J.R. & Harvey, J. (2014) Biogeography of kwongan: Origins, diversity, endemism and vegetation patterns. *Plant life on the sandplains in*

*southwest Australia, a global biodiversity hotspot* (ed. H. Lambers). UWA Publishing, Crawley, Western Australia.
Norman, M.A., Koch, J.M., Grant, C.D., Morald, T.K. & Ward, S.C. (2006) Vegetation Succession After Bauxite Mining in Western Australia. *Restoration Ecology,* **14,** 278-288.
Peddle, S. (2020) Changes to soil bacterial communities following post-mining rehabilitation in the Western Australian Jarrah forest. Honours thesis, Flinders University.
Pielou, E.C. (1966) The measurement of diversity in different types of biological collections. *Journal of Theoretical Biology,* **13,** 131-144.
Pruesse, E., Peplies, J. & Glöckner, F.O. (2012) SINA: Accurate high-throughput multiple sequence alignment of ribosomal RNA genes. *Bioinformatics,* **28,** 1823-1829.
R-Core-Team (2020) R: A language and environment for statistical computing. R Foundation for Statistical Computing, Vienna, Austria.
Rognes, T., Flouri, T., Nichols, B., Quince, C. & Mahé, F. (2016) VSEARCH: a versatile open source tool for metagenomics. *PeerJ,* **4,** e2584.
Schloss, P.D., Westcott, S.L., Ryabin, T., Hall, J.R., Hartmann, M. & Hollister, E.B. (2009) Introducing mothur: open-source, platform-independent, community-supported software for describing and comparing microbial communities. *Appl Environ Microbiol,* **75,** 7537-7541.
Specht, R.L., Roe, E.M. & Boughton, V.H. (1974) *Conservation of major plant communities in Australia and Papua New Guinea*. East Melbourne : Commonwealth Scientific and Industrial Research Organization, East Melbourne.
Standish, R.J., Daws, M.I., Gove, A.D., Didham, R.K., Grigg, A.H., Koch, J.M., Hobbs, R.J. & Gibson, D. (2015) Long-term data suggest jarrah-forest establishment at restored mine sites is resistant to climate variability. *The Journal of ecology,* **103,** 78-89.
Talavera, G. & Castresana, J. (2007) Improvement of Phylogenies after Removing Divergent and Ambiguously Aligned Blocks from Protein Sequence Alignments. *Systematic Biology,* **56,** 564-577.
Tibbett, M. (2010) Large-scale mine site restoration of Australian eucalypt forests after bauxite mining: soil management and ecosystem development. *Ecology of Industrial Pollution*, pp. 309-326.
Tibbett, M., Daws, M.I., George, S.J. & Ryan, M.H. (2020) The where, when and what of phosphorus fertilisation for seedling establishment in a biodiverse jarrah forest restoration after bauxite mining in Western Australia. *Ecological engineering,* **153,** 105907.
